# I like coffee with cream and *dog?* Change in an implicit probabilistic representation captures meaning processing in the brain

**DOI:** 10.1101/138149

**Authors:** Milena Rabovsky, Steven S. Hansen, James L. McClelland

## Abstract

The N400 component of the event-related brain potential has aroused much interest because it is thought to provide an online measure of meaning processing in the brain. Yet, the underlying process remains incompletely understood and actively debated. Here, we present a computationally explicit account of this process and the emerging representation of sentence meaning. We simulate N400 amplitudes as the change induced by an incoming stimulus in an implicit and probabilistic representation of meaning captured by the hidden unit activation pattern in a neural network model of sentence comprehension, and we propose that the process underlying the N400 also drives implicit learning in the network. The model provides a unified account of 16 distinct findings from the N400 literature and connects human language processing with successful deep learning approaches to language processing.

---

The N400 component of the event-related brain potential (ERP) has received a great deal of attention, as it promises to shed light on the brain basis of meaning processing. The N400 is a negative deflection recorded over centro-parietal areas peaking around 400 ms after the presentation of a potentially meaningful stimulus. The first report of the N400 showed that it occurred on presentation of a word violating expectations established by context: given “I take my coffee with cream and …” the anomalous word *dog* produces a larger N400 than the congruent word *sugar*^1^. Since this study, the N400 has been used as a dependent variable in over 1000 studies and has been shown to be modulated by a wide range of variables including sentence context, category membership, repetition, and lexical frequency, amongst others^2^. However, despite the large amount of data on the N400, its functional basis continues to be debated: various competing verbal descriptive theories have been proposed^3–8^, but their capacity to capture all the relevant data is difficult to determine unambiguously due to the lack of implementation, and none has yet offered a generally accepted account^2^.

Here, we provide both support for and formalization of the view that the N400 reflects the input-driven update of a representation of sentence meaning – one that implicitly and probabilistically represents all aspects of meaning as it evolves in real time during comprehension^2^. We do so by presenting an explicit computational model of this process. The model is trained and tested using materials generated by a simplified artificial microworld (see below) in which we can manipulate variables that have been shown to affect the N400, allowing us to explore how these factors affect processing. The use of these synthetic materials prevents us from simulating N400 responses to the specific sentences used in empirical experiments. Nevertheless, using these artificial materials, we are able to show that the model can capture the effects of a broad range of factors on N400 amplitudes.

The model does not exactly correspond to any existing account of the N400, as it implements a distinct perspective on language comprehension. Existing accounts are often grounded, at least in part, in modes of theorizing based on constructs originating in the 1950’s^9^, in which symbolic representations (e.g., of the meanings of words) are retrieved from memory and subsequently integrated into a compositional representation – an annotated structural description thought to serve as the representation of the meaning of a sentence^10–12^. Even though perspectives on language processing have evolved in a variety of ways, many researchers maintain the notion that word meanings are first retrieved from memory and subsequently assigned to roles in a compositional representation. The account we offer here does not employ these constructs and thus may contribute to the effort to rethink aspects of several foundational issues: What does it mean to understand language? What are the component parts of the process? Do we construct a structural description of a spoken utterance in our mind, or do we more directly construct a representation of the speaker’s meaning? Our work suggests different answers than those often given to these questions.

Our model, called the *Sentence Gestalt* (SG) model, was initially developed 30 years ago^13,14^ with the goal of illustrating how language understanding might occur without relying on the traditional mode of theorizing described above. The model sought to offer a functional-level characterization of language understanding in which each word in a sentence someone hears or reads provides clues that constrain the formation of an implicit representation of the event or situation described by the sentence. The initial work with the model^14^ established that it could capture several core aspects of language, including the ability to resolve ambiguities of several kinds; to use word order and semantic constraints appropriately; and to represent events described by sentences never seen during the network’s training. A subsequent model using a similar approach successfully mastered a considerably more complex linguistic environment^15^. The current work extending the SG model to address N400 amplitudes complements efforts to model neurophysiological details underlying the N400^16–18^.

The design of the model (Fig. 1) reflects the principle that listeners continually update their representation of the situation or event being described as each incoming word of a sentence is presented. The representation is an internal representation (putatively corresponding to a pattern of neural activity, modeled in an artificial neural network) called the *sentence gestalt* (SG) that depends on connection-based knowledge in the *update* part of the network. The SG pattern can be used to guide responses to potential queries about the event or situation being described by the sentence (see *Implicit probabilistic theory of meaning* section in *online methods*). The model is trained with sentences and queries about the events the sentences describe, so that it can, if probed, provide responses to such queries. Although we focus on a very simple microworld of events and sentences that can describe them, the model exemplifies a wider conception of a neural activation state that represents a person’s subjective understanding of a broad range of situations and of the kinds of inputs that can be used to update this understanding. The input could be in the form of language expressing states of affairs (e.g., “Her hair is red.”) or even non-declarative language. For example, the question “What time is it?” communicates that the speaker would like to know the time. Though we focus only on linguistic input here, the input guiding the formation of these representations could also come from witnessing events directly; from pictures, sounds, or movies; or from any combination of linguistic or other forms of input.

**Figure 1.**
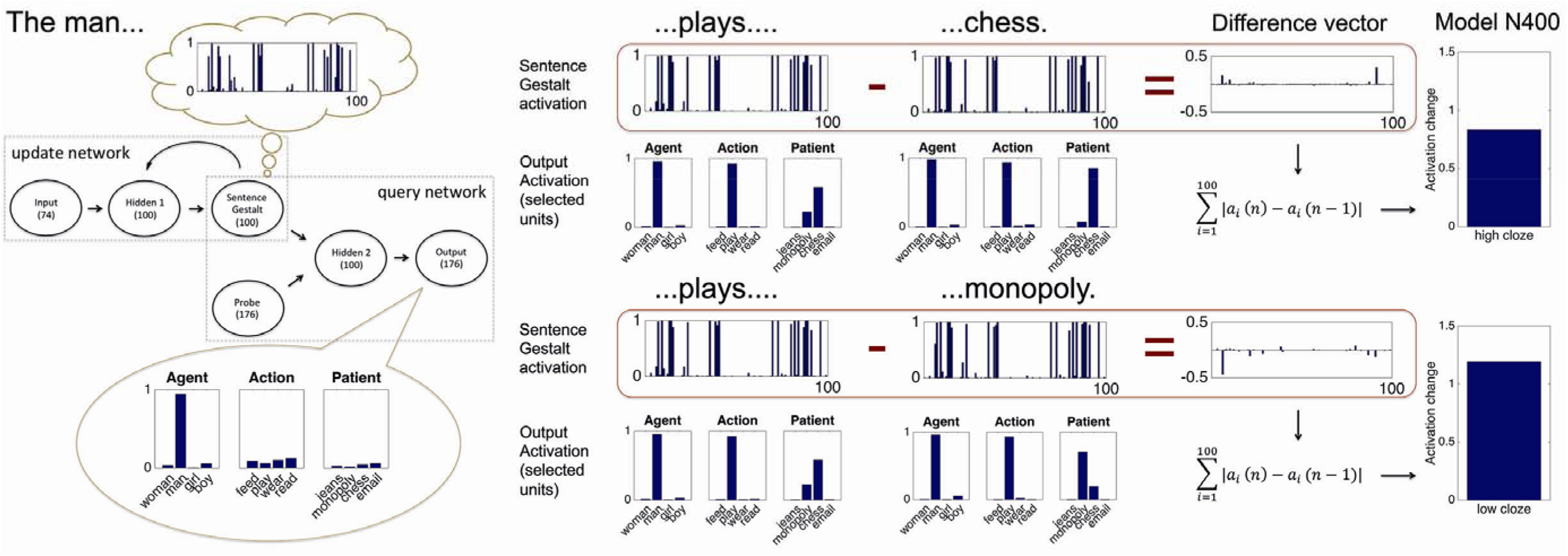
The Sentence Gestalt (SG) model architecture, shown processing a sentence with a high or low cloze probability ending, and the model’s N400 correlate. The model (gray boxes on the left) consists of an update network and a query network. Ovals represent layers of units (and number of units in each layer). Arrows represent all-to-all modifiable connections; each unit applies a sigmoid transformation to its summed inputs, where each input is the product of the activation of the sending unit times the weight of that connection. In the update part of the model, each incoming word is processed through layer Hidden 1 where it combines with the previous activation of the SG layer to produce the updated SG pattern corresponding to the updated implicit representation of the event described by the sentence. During training, after each presented word, the model is probed concerning all aspects of the described event (e.g. agent, “man”, action, “play”, patient, “monopoly”, etc.) in the query part of the network. Here, the activation from the probe layer combines via layer Hidden 2 with the current SG pattern to produce output activations. Output units for selected query response units activated in response to the agent, action, and patient probes are shown; each query response includes a distinguishing event feature (e.g. ‘man’, ‘woman’, as shown) as well as other features (e.g., ‘person’, ‘adult’, not shown) that capture semantic similarities among event participants; see Supplementary Table 1). After presentation of “The man”, the SG representation (thought bubble at top left) supports activation of the correct event features when probed for the agent and estimates the probabilities of action and patient features consistent with this agent. After the word “plays” (shown twice in the middle of the figure) the SG representation is updated and the model now activates the correct features given the agent and action probes, and estimates the probability of alternative possible patients. These estimates reflect the model’s experience, since the man plays chess with higher probability than monopoly. If the next word is “chess” (top), the change in the pattern of activation on the SG layer(summed magnitudes of changes shown in ‘Difference vector’) is smaller than if the next word is “monopoly” (bottom). The change signal, called the Semantic Update (SU) is the proposed N400 correlate (right). It is larger for the less probable ending (monopoly, bottom) as compared to the more probable ending (chess, top).

The magnitude of the update produced by each successive word of a sentence corresponds to the change in the model’s implicit representation that is produced by the word, and it is this change, we propose, that is reflected in N400 amplitudes. Specifically, the *semantic update* (SU) induced by the current word *n* is defined as the sum across the units in the SG layer of the absolute value of the change in each unit’s activation produced by the current word *n*. For a given unit (indexed below by the subscript *i*), the change is simply the difference between the unit’s activation after word *n* and after word *n*-1:

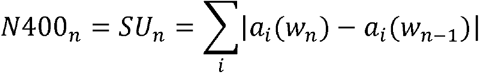

This measure can be related formally to a Bayesian measure of surprise^19^ and to the signals that govern learning in the network (see *online methods* and below). Indeed, we propose a new learning rule driven by the semantic update, allowing the model to address how language processing even in the absence of external event information can drive learning about events and about how speakers use language to describe them.

How does the semantic update capture the N400? After a listener has heard “I take my coffee with cream and …” our account holds that the activation state already implicitly represents a high subjective probability that the speaker takes her coffee with cream and sugar, so the representation will change very little when the final word “sugar” is presented, resulting in little or no change in activation, and thus a small N400 amplitude. In contrast, the representation will change much more if “dog” is presented instead, corresponding to a much larger change in subjective probabilities of the characteristics of the event being described, reflected in a larger change in the pattern of activation and thus a larger N400 amplitude.

Learning takes place in the model over an extended time course thought of as loosely corresponding to the time course of human development into early adulthood, based on the gradual accumulation of experience about events and the sentences speakers use to describe them. Among other things, this means that the extent of the semantic update that occurs upon the presentation of a particular word in a particular context depends not only on the statistics of the environment, but also on the extent of the model’s training – thereby allowing it to address changes in N400 responses as a function of experience.

## Distinctive Features of the Sentence Gestalt Model

Several aspects of the model’s design and behavior are worth understanding in order to see how it accounts for the findings we apply it to below. First, the model is designed to form a representation of the *situation or event* described by the sentence that it hears, rather than a representation of the sentence itself. This contrasts with other models of language processing that focus primarily on the updating of *linguistic* expectations. Such expectations include expectations about the occurrence of specific words or of specific structural relationships that may be tied closely to the sentence itself^11,12^. In this regard our approach is similar to approaches in which the constructed representation specifies entities and their relations rather than relationships among words^20^. Furthermore, unlike most other models, the SG model does not contain separate modules that implement distinct stages of lexical access or syntactic parsing on the way to the formation of a representation of the event. Instead the model simply maps from word forms to an implicit probabilistic representation of the overall meaning of the sentence.

Second, we as modelers make no stipulations of the form or structure of the model’s internal representations. Rather, these representations are shaped by the statistics of the experiences it is trained on, as in some language representation models developed by other groups in recent years^21,22^ To train the model, we require a way of providing it with information about the event described by the sentence. We follow the choice made in the original implementation, in which events are described in terms of an action, a location, a situation (such as ‘at breakfast’), the actor or agent in the event, and the object or patient to which the action is applied. Critically, the event description is not the model’s internal representation, but is instead a characterization of those aspects of the event described by the sentence that the representation should be capable of describing if probed. In this way our model is similar to contemporary deep learning models such as Google’s Neural Machine Translation (GNMT) system^23^. Our model is simpler than this system, which employs more layers of neuron-like processing units, but the models are similar in avoiding representational commitments. Though GNMT is focused on sequences of words rather than sentence meanings, it likewise makes no stipulations of the form or structure of the internal representation generated from an input sentence; instead the representation is shaped by the process of learning to predict the translation of a sentence in one language in other languages. The relative success of such systems where other approaches have struggled can be seen as supporting the view that a commitment to *any* stipulated form of internal representation is an impediment to capturing the nuanced, *quasiregular* nature of language^24,25^.

Third, the model responds to any words presented to it, independently of whether they form sentences, as implemented in the *update* part of the network (Fig 1). This allows the model to address N400’s evoked by words presented in pairs or even in isolation. We view this process as a largely automatic process that proceeds independently of the intention of the listener. Whether they are in sentences or not, the SG activity produced by words will reflect aspects of events in which they occur, in line with embodied or grounded approaches to the representation of meaning^26,27^. The explicit computation of responses to queries about events is used during training to allow the model learn to map from sentences to meaning, but this process is not thought of as contributing to the N400. Indeed, this process would not ordinarily occur during an N400 experiment, when external sources of information about events are not generally available.

Finally, we do not see the process reflected in the N400 and captured by the SG as the only process that contributes to on-line language understanding. While we see this process as lying at the heart of comprehension, other processes also play important roles. This view is consistent with the fact that other ERP components appear to reflect different aspects of language processing. Specifically, we see the N400 as reflecting an implicit process that operates quickly and automatically as linguistic input is presented. Language processing may also involve other components that might form expectations about specific word-forms and their sequencing that are not captured by the SG model or the N400. Furthermore, the initial representation that the model forms as it processes language in real time may not always correspond to the final understood meaning of a sentence. Other processes may come into play in understanding sentences describing implausible events or sentences with unusual structure (such as garden-path sentences), and these processes may result in changes to the meaning representation that is ultimately derived from reading or listening to a linguistic input. In the *Discussion* below we consider how the formation of an initial, implicit representation of meaning, as captured by the SG model, might fit into this broader picture, and how our findings may inform discussions of other aspects of human language processing.

## Advantages from Using a Controlled Micro-World Training Environment

As previously noted, we use an artificial corpus of {*sentence, event*} training examples produced by a generative model that embodies a simplified and controlled micro-world in which the statistics of events, the properties of the objects that occur in them, and the words used in sentences about these events are completely controlled by the modeler (see *online methods*). Thus, we simulate the influence of specific variables (such as frequency or cloze probability) that are manipulated in experiments by manipulating these variables in our synthetic materials. It may eventually be possible to train a successor to our model on a much larger corpus of real sentences, allowing modeling of the semantic update produced by the actual materials used in empirical experiments. Such a success would still leave open the question of what factors were responsible for the model’s behavior. Our approach, relying on a synthetic corpus, allows us to build into the training materials manipulations of variables corresponding to those explored in the designs of the experiments we are modeling. For example, we can separately manipulate how frequently an object designated by a particular word appears in an event of a particular type (e.g. how often a knife is used for spreading butter on bread) and the extent to which the properties of the object signaled by a word are consistent with the properties of the objects that typically appear in events of this type (e.g. an axe, though never used in spreading, is more semantically similar to a knife than a chair is). Thus we are able to separate predictability from semantic similarity more cleanly than might be possible using a large corpus of real sentences.

## Results

We report sixteen simulations of well-established findings in the N400 literature. We chose these findings to illustrate how the model can address a broad range of empirical findings taken as supporting diverse and sometimes conflicting descriptive theories of the functional basis of the N400 (see Table 1). We focus on language-related effects but note that both linguistic and non-linguistic information contribute to changes in semantic activation as reflected by the N400. Throughout what follows, we define an *N400 effect* as the difference in negativity between an experimental condition and a control or baseline condition, simulated as the difference in the size of the SU between corresponding conditions in our simulations.

**Table 1.**
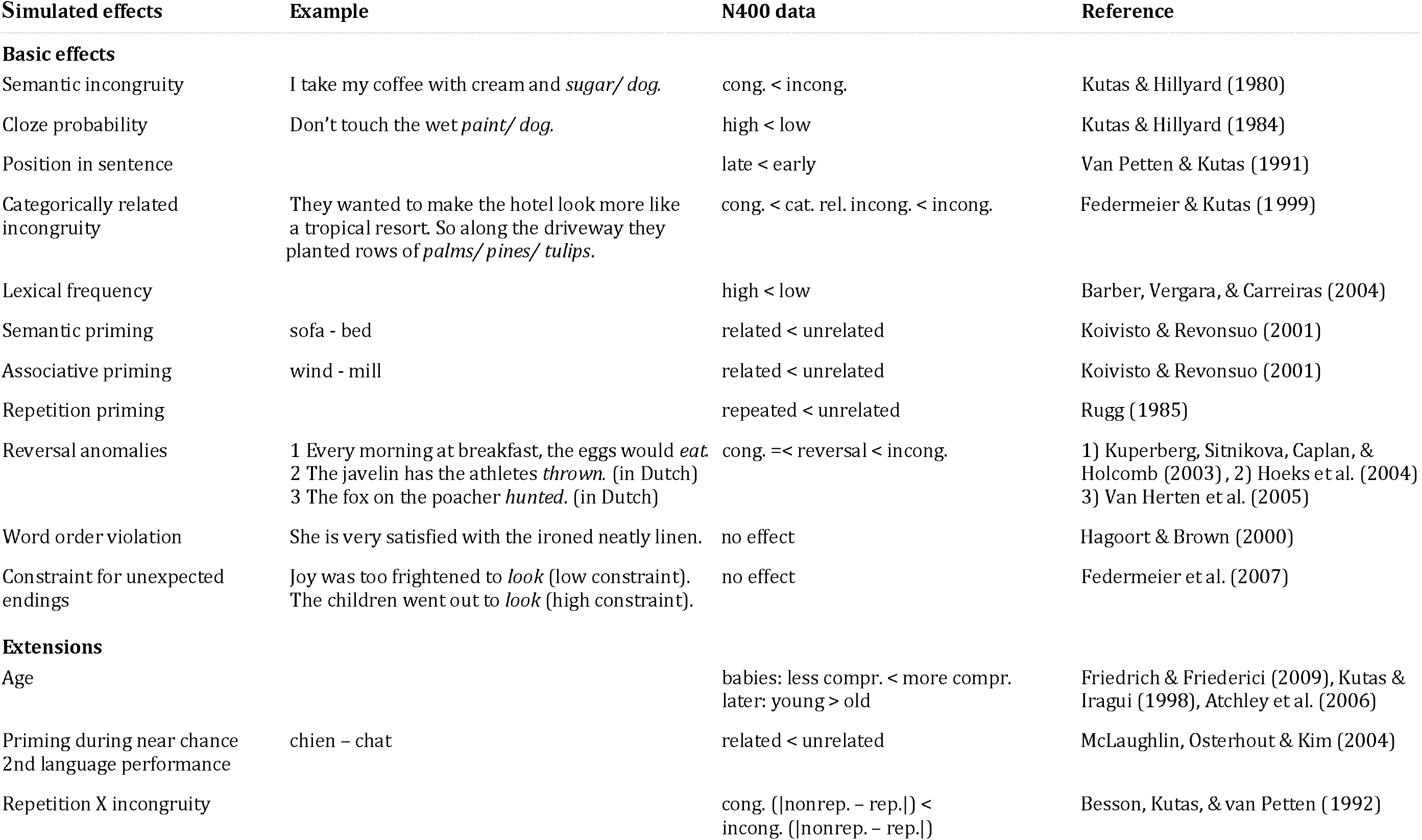
Overview of simulated effects, cong: congruent; incong.: incongruent; cat rel.: categorically related; rev. anom.: reversal anomaly; compr.: comprehension; rep.: repeated; nonrep.: nonrepeated.

### Basic effects

#### From “violation signal” to graded reflection of surprise

The N400 effect was first observed as a large negative deflection in the EEG after a semantically anomalous sentence completion such as e.g. “He spread the warm bread with *sods*”^1^ as compared to a high probability congruent completion (*butter*). Correspondingly, in our model, SU is significantly larger for sentences with endings that are both semantically and statistically inconsistent with the training corpus compared to semantically consistent, high-probability completions (Fig. 2a and Supplementary Fig. 1a). Soon after the initial study it became clear that the N400 is graded, with larger amplitudes for acceptable sentence continuations with lower cloze probability (defined as the percentage of participants that continue a sentence fragment with that specific word in an offline sentence completion task), as in the example “Don’t touch the wet *dog* (low cloze)/*paint* (high cloze)”^28^. This is also captured by the model: it exhibits larger SU for sentence endings presented with a low (.3) as compared to a high probability (.7) during training^1^ (Fig. 2b, Fig. 1, and Supplement Fig. 1b).

**Figure 2.**
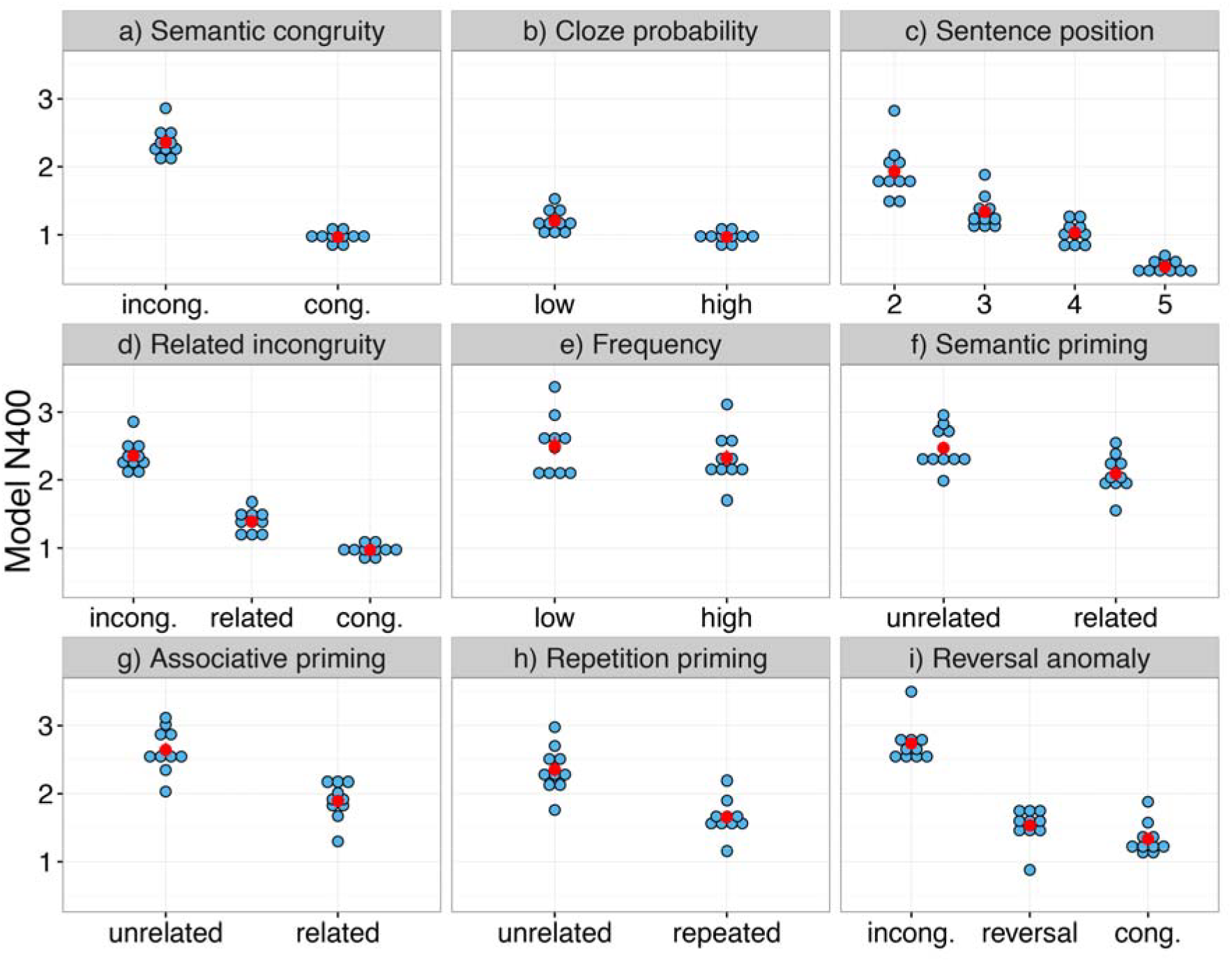
Simulation results for the basic effects. Displayed is the model’s N400 correlate, i.e. the update of the Sentence Gestalt layer activation – the model’s probabilistic representation of sentence meaning - induced by the new incoming word. Cong, congruent; incong., incongruent. See text for details of each simulation. Each blue dot represents the results for one independent run of the model, averaged across items per condition; the red dots represent the means for each condition, and red error bars represent +/− SEM (sometimes invisible because bars may not exceed the area of the red dot). Statistical results (t_1_ from the model analyses, t_2_ from the item analyses, Cohen’s d [please note that effect sizes might be larger in simulations than in empirical experiments due to the high level of noise present in EEG signals], 95% confidence interval for the condition difference, CI): a, semantic incongruity: t_1(9)_ = 25.00, p < .0001, d = 7.91, 95% CI[1.26, 1.51]; t_2(9)_ = 11.24, p < .0001, d = 3.55, 95% CI [1.11, 1.67]; b, cloze probability: t_1(9)_ = 8.56, p < .0001, d = 2.71, 95% CI[.18, .30]; t_2(9)_ = 6.42, p = .0001, d = 2.03, CI95% [.15, .32]; c, position in sentence: t_1(9)_ = 8.17, p <.0001, d = 2.58, 95% CI[.43, .76]; t_2(11)_ = 43.54, p <.0001, d = 12.57, 95% CI [.57, .63] from the second to the third sentence position; t_1(9)_ = 4.73, p =.003, d = 1.50, 95% CI[.16, .44], t_2(11)_ = 4.66, p =.0018, d = 1.34, 95% CI[.16, .44] from the third to the fourth position; t_1(9)_ = 17.15, p < .0001, d = 5.42, 95% CI [.44, .58]; t_2(11)_ = 12.65, p <.0001, d = 3.65, 95% CI [.42, .60] from the fourth to the fifth position; d, categorically related incongruities were larger than congruent, t_1(9)_ = 10.63, p < .0001, d = 3.36, 95% CI[.33, .51]; t_2(9)_ = 3.31, p = .018, d = 1.05, 95% CI [.13, .71], and smaller than incongruent continuations, t_1(9)_ = 14.69, p < .0001, d = 4.64, 95% CI[.82, 1.11]; t_2(9)_ = 12.44, p < .0001, d = 3.94, 95% CI [.79, 1.14]; e, lexical frequency: t_1(9)_ = 3.13, p = .012, d = .99, 95% CI [.05, .31]; t_2(13)_ = 3.26, p = .0062, d = .87, 95% CI [.06, .30]; f, semantic priming: t_1(9)_ = 14.55, p < .0001, d = 4.60, 95% CI[.32.44]; t_2(9)_ = 8.92, p < .0001, d = 2.82, 95% CI[.28, .48]; g, associative priming: t_1(9)_ = 14.75, p < .0001, d = 4.67, 95% CI [.63, .86]; t_2(9)_ = 18.42, p < .0001, d = 5.82, 95% CI [.65, .84]; h, immediate repetition priming: t_1(9)_ = 16.0, p < .0001, d = 5.07, 95% CI[.60, .80]; t_2(9)_ = 18.93, p < .0001, d = 5.99, 95% CI[.62, .79]; i, reversal anomaly: t_1(9)_ = 2.09, p = .199, d = .66, 95% CI[.02, .41]; t_2(7)_ = 5.67, p = .0023, d = 2.0, 95% CI [.12, .28], for the comparison between congruent condition and reversal anomaly; t_1(9)_ = 10.66, p < .0001, d = 3.37, 95% CI[.95, 1.46]; t_2(7)_ = 3.56, p = .028, d = 1.26, 95% CI [.40, 2.0] for the comparison between reversal anomaly and incongruent, and t_1(9)_ = 28.39, p < .0001, d = 8.98, 95% CI[1.29, 1.51]; t_2(7)_ = 4.21, p = .012, d = 1.49, 95% CI[.61, 2.19].

All Supplementary Figures currently appear at the end of the manuscript.

The graded character of the underlying process is further supported empirically by the finding that N400s gradually decrease across the sequence of words in normal congruent sentences^29^. SU in the model correspondingly shows a gradual decrease across successive words in sentences (Fig. 2c and Supplementary Fig. 1c; see *online methods* for details).

#### Expectancy for words or semantic features?

The findings discussed above would be consistent with the view that N400s reflect the inverse probability of a word in a specific context (i.e. word surprisal^30^), and indeed, a recent study observed a significant correlation between N400 and word surprisal measured at the output layer of a simple recurrent network (SRN) trained with a naturalistic corpus to predict the next word based on the preceding context^31^. However, there is evidence that N400s may not be a function of word probabilities *per se* but rather of probabilities of aspects of meaning signaled by words: N400s are smaller for incongruent completions that are closer semantically to the correct completion than those that are semantically more distant. For example, consider the sentence: “They wanted to make the hotel look more like a tropical resort. So, along the driveway they planted rows of …”.

The N400 increase relative to *palms* (congruent completion) is smaller for *pines* (incongruent completion from the same basic level category as the congruent completion) than for *tulips* (incongruent completion not from the same basic level category as the congruent completion)”^32^. Our model captures these results: We compared SU for sentence completions that were presented with a high probability during training and two types of never-presented completions. SU was lowest for high probability completions, as expected; crucially, among never-presented completions, SU was smaller for those that shared semantic features with high probability completions compared to those that did not (Fig. 2d and Supplementary Fig. 1d).

#### Semantic integration versus lexical access?

The sentence-level effects considered above have often been taken to indicate that N400 amplitudes reflect the difficulty or effort required to integrate an incoming word into the preceding context^7,33^. However, a sentence context is not actually needed: N400 effects can also be obtained for words presented in pairs or even in isolation. Specifically, N400s are smaller for isolated words with a high as compared to a low lexical frequency^34^; for words (e.g. “bed”) presented after a categorically related prime (e.g., “sofa”) or an associatively related prime (e.g., “sleep”) as compared to an unrelated prime^35^; and for an immediate repetition of a word compared to the same word following an unrelated prime^36^. Such N400 effects outside of a sentence context, especially the influences of repetition and lexical frequency, have led some researchers to suggest that N400 amplitudes do not reflect the formation of a representation of sentence meaning but rather lexical access to individual word meaning^3,16^. As previously noted, the SG pattern probabilistically represents the meaning of a sentence if one is presented, but the model will also process words presented singly or in pairs. This corresponds to the assumption that there is no separate language processing system to process single words or word pairs. Instead the system used to process sentences will deal with whatever linguistic input it is presented with in a rather automatic fashion. As noted in the introduction, the model will presumably co-activate implicit representations of the semantic features of the events in which words typically occur, in line with embodied/grounded approaches to the representation of word meaning^26,27^, but no explicit event representation (in the sense of producing responses to queries) is assumed to be computed during N400 experiments.

The model captures all four of the above-mentioned effects: First, SU was smaller for isolated words that occurred relatively frequently during training (Fig. 2e and Supplementary Fig. 1e). Furthermore, SU was smaller for words presented after words from the same semantic category as compared to words from a different category (Fig. 2f and Supplementary Fig. 1f), and smaller for words presented after associatively related words (objects presented after a typical action as in “chess” following “play”) as compared to unrelated words (objects presented after an unrelated action as in “chess” following “eat”) (Fig. 2g and Supplementary Fig. 1g). Finally, SU was smaller for immediately repeated words as compared to words presented after unrelated words (Fig. 2h and Supplementary Fig. 1h). Thus, the model demonstrates that the same mechanism that captures N400 effects in sentences – and that supports correct responses to probes about these sentences – also shows the effects seen with word pairs and single words.

#### Reversal anomalies and the N400

A finding that has puzzled the N400 community is the lack of a robust N400 effect in reversal anomaly sentences: That is, only a very small N400 increase occurs at the verb in sentences such as “Every morning at breakfast, the eggs would *eat*…” when compared with corresponding congruent sentences such as “Every morning at breakfast, the boys would *eat*…”^37^. The result is seen as surprising since English syntactic conventions typically map nouns preceding an action verb (presented in active voice) to the agent rather than an object or patient role (though this is not always the case, as in usages such as ‘the book reads well’). The small size of the N400 effect in reversal anomalies is typically accompanied by an increase of the P600, a subsequent positive potential. In contrast, N400 but not P600 amplitudes are considerably larger in sentence variations such as “Every morning at breakfast, the boys would *plant*…”^37^.

This pattern of results has sparked considerable theoretical uncertainty. Many researchers have taken these findings to indicate that the word *eggs* in the given context is easily integrated into a representation of sentence meaning because *eggs* is (at least temporarily) interpreted as the patient instead of the agent of eating. Such a situation has been called a temporary “semantic illusion”^38^. Our account is partly in line with this account, though we describe such a state of mind as an *event-probability based interpretation* to avoid the implication that syntax must always be treated as the definitive cue when syntax and other considerations conflict. Others^39^ have argued that the N400 is not related to semantic integration but instead reflects the difficulty the reader experiences in activating the meaning of the critical word. The idea is that the retrieval of the meaning of *eat* is facilitated by priming from *breakfast* and *eggs*, which both occur prior to *eggs* in the reversal anomaly sentence; in contrast *plant* would not be facilitated by the prior context. On this view, the process of understanding sentence meaning is associated with the P600 rather than the N400. In what follows, we show that our model’s SU at the occurrence of the verb in reversal anomaly sentences is similar in size to the SU at the verb in the corresponding control sentences, consistent with the small N400 effect seen in the empirical experiments.

The first relevant simulation used materials in our standard training corpus to simulate the reversal anomaly experiment described above (see *online methods* for details). We found that the semantic update in the SG model reproduces the pattern seen in the human N400 data. That is, the model exhibited only a very slight increase in SU for reversal anomalies (e.g., “At breakfast, the eggs *eat…*”) as compared to congruent continuations (e.g., “At breakfast, the man *eats*…’), and a substantial increase in SU for incongruent continuations (e.g., “At breakfast, the man *plants*…”) (Fig. 2i and Supplementary Fig. 1i). Analysis of the query network’s response to relevant probes suggests that the model indeed maintains an event probability-based interpretation, in that these responses continue to favor the eggs as the patient instead of the agent of eating even after the word *eat* is presented (Supplementary Fig. 2).

The second simulation addresses the absence of a robust N400 effect in reversal anomaly sentences in which both participants were animate beings that could occur as agents (this was not the case in the first simulation and another we describe in *online methods*). As examples, consider these materials used in the relevant experiment^40^: “De vos die op de stroper *joeg*…” (lit: The fox who on the poacher *hunted*…; paraphrase: The fox who hunted the poacher) and “De zieke die in de chirurg sneede…” (lit: The patient who into the surgeon cut…; paraphrase: The patient who cut into the surgeon…). Here, the syntactically supported interpretations are inconsistent with event probabilities^2^, yet both the participants are animate. In addition, both can be agents in events involving the other participant (a fox could watch a poacher, and a patient could stand in front of a surgeon), and both can engage in the relevant action (hunt something or cut into something). What makes these cases anomalous is that in hunting events involving poachers and foxes, it is always the poachers that hunt the foxes; and in events involving surgeons and patients where one is cutting into the other, it is always the surgeons that cut into the patients.

To address such cases, we conducted an additional simulation involving reversal anomaly sentences, focusing on the experiment that used the cited examples among others^40^. The experiment was done in Dutch with Dutch word order conventions, and this is critical because it means that both noun phrases are presented prior to the occurrence of the verb. We therefore trained an additional model with sentences using Dutch word order, using event scenarios involving a central action (e.g. hunting or cutting into something) and two central animate participants (poacher and fox, or surgeon and patient) as well as alternative actions and participants (see *online methods* for details). The scenarios were set up to align with the characteristics of the materials used in the target experiment, which we obtained from the authors (though we used simple one-clause sentences rather than sentences with embedded clauses as in the experiment). Specifically, when both central participants occur in the same event, the action is usually, but not always, the central action. When the action is the central action, one of the central participants (called the *central agent*) is always the agent and the other (called the *central patient*) is always the patient (e.g. poacher can hunt fox, but fox never hunts poacher, and surgeon can cut into patient, but patient never cuts into surgeon). When both central participants participate in one of the alternative actions, the central patient can also be the agent (e.g., fox can watch the poacher, patient can stand in front of the surgeon). Furthermore, when only one of the central participants occurs in an event, either can be the agent of the central action (foxes can hunt chickens, patients can cut into turkeys). The central patient (e.g., the fox or the patient) was overall a very likely agent, but was less likely to be the agent of the central action (hunting or cutting into something) than the central agent (poacher or surgeon; see *online methods* for full details). The materials also reflect the fact that word order is a less reliable cue to role assignment in Dutch than in English^41^.

Using sentences from these scenarios, the model again successfully captures the relatively small N400 effect at the presentation of the verb in reversal anomaly sentences. The model exhibited only a very slight increase in SU for reversal anomalies such as “The fox on the poacher *hunted*” as compared to congruent control sentences such as “The poacher on the fox *hunted*” and a substantial increase in SU for incongruent continuations such as “The poacher on the fox *planted*” (see Fig. 3 and Supplementary Fig. 4).

**Figure 3.**
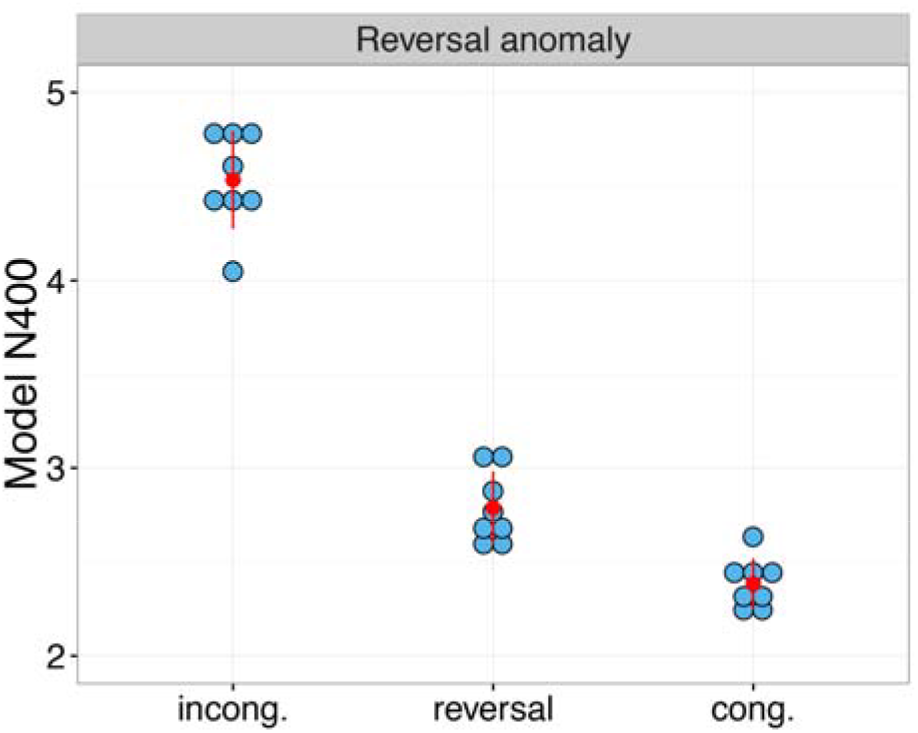
Simulation results for a type of reversal anomaly where both event participants can be agents and can perform the action of interest (see text for details). Cong., congruent; incong., incongruent; reversal, reversal anomaly. Each blue dot represents the results for one independent run of the model, averaged across items per condition. In line with the empirical data, SU in the reversal anomaly condition is only slightly increased as compared to the congruent condition, while it is considerably larger in the incongruent condition. For the comparison between congruent condition and reversal anomaly: t_1(9)_ = 4.15, p = .008, d = 1.31, 95% CI [.18, .62]; t_2(7)_ = 4.71, p = .007, d = 1.67, 95% CI[.20, .61]; for the comparison between congruent and incongruent conditions: t_1(9)_ = 9.90, p < .0001, d = 3.13, 95% CI [1.66, 2.64], t_2(7)_ = 27.34, p < .0001, d = 9.67, 95% CI[1.96, 2.34]; for the comparison between reversal anomaly and incongruent condition: t_1(9)_ = 7.91, p < .0001, d = 2.50, 95% CI[1.25, 2.25], t_2(7)_ = 17.95, p < .0001, d = 6.35, 95% CI[1.52, 1.98].

It may seem surprising that the model does not exhibit a substantially larger SU in the role-reversed sentences compared to the corresponding control sentences. To understand this pattern, we probed the network’s responses to Action, Agent and Patient probes in both types of sentences. While the model’s interpretation of the congruent sentence was unambiguous and clear, it exhibited uncertainty in its role assignments when processing the reversal anomaly sentences, due to conflicting constraints imposed by word order and event probabilities. This conflict was not reflected in a large SU at the occurrence of the verb because it already started at the occurrence of the second noun and was not resolved by the presentation of the verb (for details, see *online methods* and Suppl. Fig. 14).

In summary, the model shows that the small N400 effect in reversal anomalies is consistent with the view that the N400 reflects the updating of an implicit representation of sentence meaning as implemented in the SG model. The model is somewhat in line with previous accounts favoring a role for plausibility constraints (based largely on previously experienced event probabilities) in sentence processing^38^. However, in our model, the initial heuristic comprehension process underlying N400 amplitudes is not purely based on event probabilities. Instead, the model is sensitive to both event probabilities and syntactic constraints, and from this perspective the small N400 effect in reversal anomalies does not necessarily reflect a clear-cut event probability based representation (of, for instance, the poacher hunting the fox; see Suppl. Fig. 14). Instead, the finding may reflect a state of unresolved conflict between different cues. As discussed in more detail in the discussion section, below, other processes, which may be associated with the P600, could resolve the conflict between competing interpretations in situations where the initial implicit representation is uncertain and undetermined. We note that the specific pattern of role expectations exhibited in the model depend on the event probabilities and syntactic conventions that we have adopted in constructing our training and testing materials. The details of the model’s estimates of role assignments in situations of cue conflict will strongly depend on the statistics of its training environment. Further research, involving parametric variation of the relevant dimensions in the training environment and in empirical experiments, as well as analysis of event probabilities as reflected in actual human experiences, will be necessary to examine the influence of these factors on the model’s internal representations and to determine if the model can capture the presence or absence of N400 effects in the full range of relevant situations (see *discussion*, below).

#### Specificity of the N400 to violations of semantic rather than syntactic factors

While the N400 is sensitive to a wide range of semantic variables, amplitudes are not influenced by syntactic factors such as for instance violations of word order (e.g., “The girl is very satisfied with the ironed neatly linen.”) which instead elicit P600 effects^42^. Because the model is representing the event described by the sentence, and this event itself is not necessarily affected by a change in word order, the model is likewise insensitive to such violations. To demonstrate this, we examined the model’s response to changes in the usual word order (e.g., “On Sunday, the man *the robin* feeds” compared to “On Sunday, the man *feeds* the robin), examining the size of the semantic update at the highlighted position, where the standard word-order is violated. We found that, if anything, SU was actually slightly larger in the condition with the normal as compared to the changed word order (please see Fig. 4 and Supplementary Fig. 5; significant over models but not items). This is because changes in word order also entail changes in the amount of information a word provides about the event being described; it turns out that the amount of semantic update was on average slightly larger in the sentences with normal compared to changed word order (see *online methods* for details).

**Figure 4.**
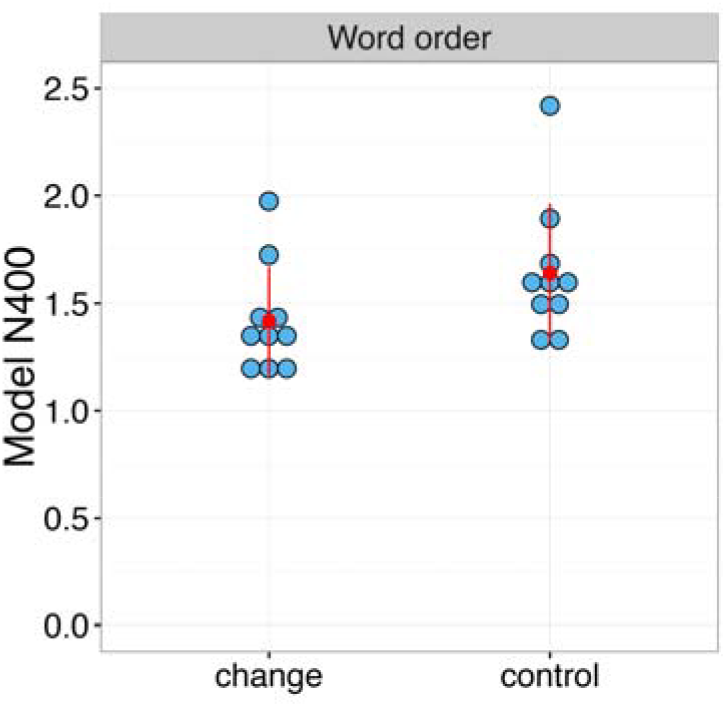
Simulation of the influence of a change in normal word order. Change, changed word order; control, normal word order. Each blue dot represents the results for one independent run of the model, averaged across items per condition; the red dots represent the means for each condition, and red error bars represent +/− SEM. Semantic update was slightly larger for normal compared to changed word order; the main effect was significant over models, t_1(9)_ = 5.94, p = .0002, d = 1.88, 95% CI [.14, .31], but not over items, t_2(9)_ = 1.56, p = .14, d = .39, 95% CI[-.08, .53].

#### No influence of constraint for unexpected endings

Another factor that does not influence the N400 but instead modulates later P600 like positivities is the constraint of a context sentence when completed with an unexpected ending^43^. Specifically, N400 amplitude is the same independent of whether the target is unpredictable because the context predicts no specific word (e.g., “Joy was too frightened to *look*.”) or because the context predicts a specific word that does not arrive (e.g., “The children went outside to *look*.” where *play* would be highly expected). Correspondingly, in the model SU was equally large for words that are unexpected because the context is unconstraining (e.g., “The man likes the *email*.”) as for words that are unexpected because they violate a specific expectation (e.g., “The man eats the *email*.”; see Fig. 5 and Supplementary Fig. 6). This finding highlights the fact discussed above that the N400 (and SU in the model) correspond to the amount of unexpected semantic information (in the sense of Bayesian surprise) and do not constitute a violation signal per se.

**Figure 5.**
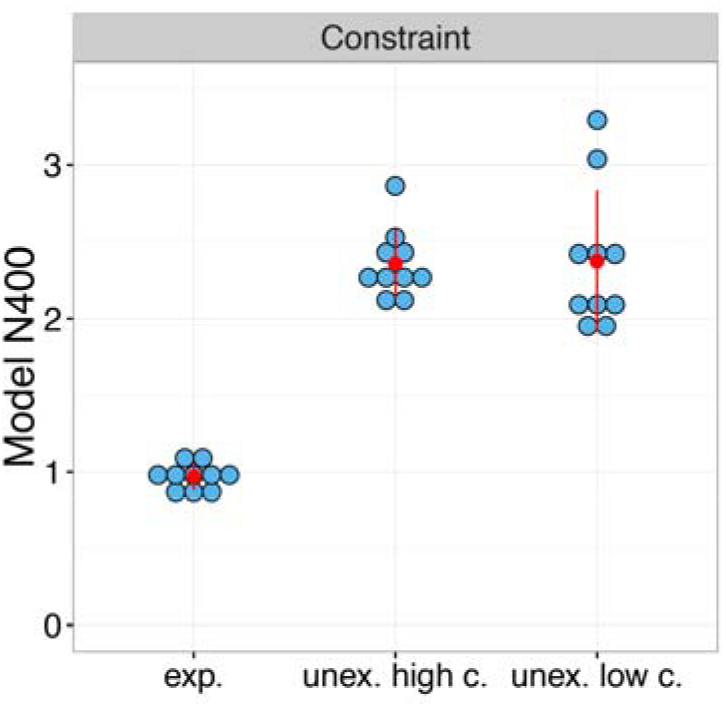
Simulation of the influence of constraint for unexpected endings. Exp., expected; unex., unexpected; c., constraint. Each blue dot represents the results for one independent run of the model, averaged across items per condition; the red dots represent the means for each condition, and red error bars represent +/− SEM. Semantic update did not differ between unexpected high constraint endings and unexpected low constraint endings, t_1(9)_ = 0.13, p = .90, d = .04, 95% CI[-.24, .27]; t_2(9)_ = 0.12, p = .91, d = .04, 95% CI[-.27, .30], while for expected endings it was considerably lower than both, for unexpected high constraint, t_1(9)_ = 25.00, p < .0001, d = 7.91, 95% CI[1.26, 1.52]; t_2(9)_ = 11.24, p < .0001, d = 3.55, 95% CI [1.11, 1.67], and for unexpected low constraint endings, t_1(9)_ = 10.21, p < .0001, d = 3.23, 95% CI[1.09, 1.72]; t_2(9)_ = 23.33, p < .0001, d = 7.38, 95% CI[1.27, 1.54].

### Extensions

In all of the simulations above, it would have been possible to model the phenomena by treating the N400 as a direct reflection of change in estimates of event-feature probabilities, rather than as reflecting the update of an implicit internal representation that latently represents these estimates in a way that only becomes explicit when queried. In this section, we show that the implicit semantic update (measured at the hidden SG layer) and the change in the networks’ explicit estimates of feature probabilities in response to probes (measured at the output layer) can pattern differently, with the implicit semantic update patterning more closely with the N400, supporting a role for the update of the learned implicit representation rather than explicit estimates of event-feature probabilities or objectively true probabilities in capturing neural responses (see *online methods* for details of these measures). We then consider how the implicit semantic update can drive connection-based learning in the update network, accounting for a final observed pattern of empirical findings.

#### Development

N400s change with increasing language experience and over developmental time. The examination of N400 effects in different age groups has shown that N400 effects increase with comprehension skills in babies^44^ but later decrease with age^45,46^. A comparison of the effect of semantic congruity on SU at different points in training shows a developmental pattern consistent with these findings (Fig. 6, top, and Supplementary Fig. 7, top): the size of the congruity effect on SU first increased and then decreased as training proceeded. Interestingly, the decrease in the effect on SU over the second half of training was accompanied by a continuing increase in the effect of semantic congruity on the change in output activation (Fig. 6, bottom, and Supplementary Fig. 7, bottom). The activation pattern at the output layer directly reflects explicit estimates of semantic feature probabilities in that units at the output layer explicitly specify semantic features, such as for instance “can grow”, “can fly” etc., and network error (across the training environment) is minimized when the activation of each feature unit in each situation corresponds to the conditional probability of this feature in this situation (e.g., an activation state of .7 in a situation where the conditional probability of the feature is .7). Thus, in the trained model, changes in output activation induced by an incoming word approximate changes in explicit estimates of semantic feature probabilities induced by that word. The continuing increase of the congruity effect across training displayed at the bottom of Fig. 6 thus shows that changes in explicit estimates of semantic feature probabilities do not pattern with the developmental trajectory of N400 effects.

**Figure 6.**
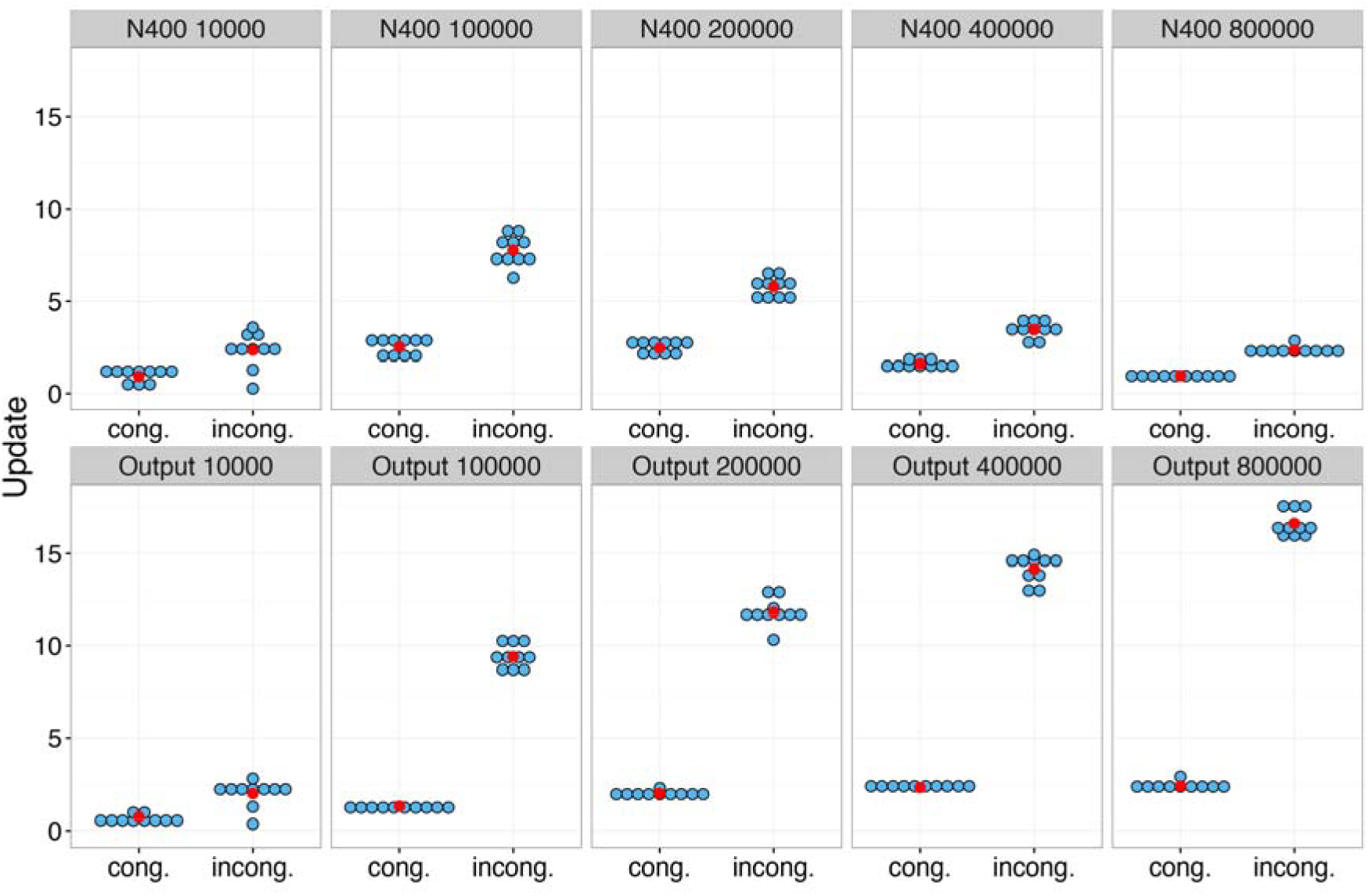
Development across training. Semantic incongruity effects as a function of the number of sentences the model has been exposed to. Top. Semantic update at the model’s hidden Sentence Gestalt layer shows at first an increase and later a decrease with additional training, in line with the developmental trajectory of the N400. Each blue dot represents the results for one independent run of the model, averaged across items per condition; the red dots represent the means for each condition, and red error bars represent +/− SEM. The size of the effect (i.e. the numerical difference between the congruent and incongruent condition) differed between all subsequent time points: t_1(9)_ = 17.02, p < .0001, d = 5.38, 95% CI[3.28, 4.29], t_2(9)_ = 6.94, p = .00027, d = 2.19, 95% CI[2.55, 5.02] between 10000 and 100000 sentences; t_1(9)_ = 7.80, p = .00018, d = 2.47, 95% CI[1.33, 2.41], t_2(9)_ = 10.05, p < .0001, d = 3.18, 95% CI[1.45, 2.29] between 100000 and 200000 sentences; t_1(9)_ = 14.69, p < .0001, d = 4.65, 95% CI[1.24, 1.69], t_2(9)_ = 6.87, p = .00029, d = 2.17, 95% CI[.98, 1.95] between 200000 and 400000 sentences; t_1(9)_ = 7.70, p = .00012, d = 2.43, 95% CI[.34, .62], t_2(9)_ = 3.70, p = .02, d = 1.17, 95% CI[.19, .78] between 400000 and 800000 sentences. Bottom. Activation update at the output layer steadily increases with additional training, reflecting closer and closer approximation to the true conditional probability distributions embodied in the training corpuss.

Instead, the change in hidden SG layer activation patterns with the N400 (Fig. 6, top), showing that the implicit and ‘hidden’ character of the model’s N400 correlate is crucial to account for the empirical data. The *decrease* in the amount of activation change at the hidden SG layer and the corresponding *increase* in the amount of activation change at the output layer over the later phase of learning shows that, as learning proceeds, less change in activation at the SG layer is needed to effectively support larger changes in explicit probability estimates.

This pattern is possible because, as noted above, the activation pattern at the SG layer does not explicitly represent the probabilities of semantic features per se; instead it provides a basis (together with the connection weights in the query network) for estimating these probabilities when probed. As connection weights in the query network get stronger throughout the course of learning, smaller changes in SG activations become sufficient to produce big changes in output activations. This shift of labor from activation to connection weights is interesting in that it might underlie the common finding that neural activity often decreases as practice leads to increases in speed and accuracy of task performance^47^.

#### Early sensitivity to a new language

A second language learning study showed robust influences of semantic priming on N400s while overt lexical decision performance in the newly trained language was still near chance^48^. We leave it to future work to do full justice to the complexity of second language learning, but as a first approximation we tested the model at a very early stage in training (Fig. 7a). Even at this early stage, SU was significantly influenced by semantic priming, associative priming, and semantic congruity in sentences (Fig. 7b and Supplementary Fig. 8) while overt estimates of feature probabilities were only weakly modulated by the words presented.

**Figure 7.**
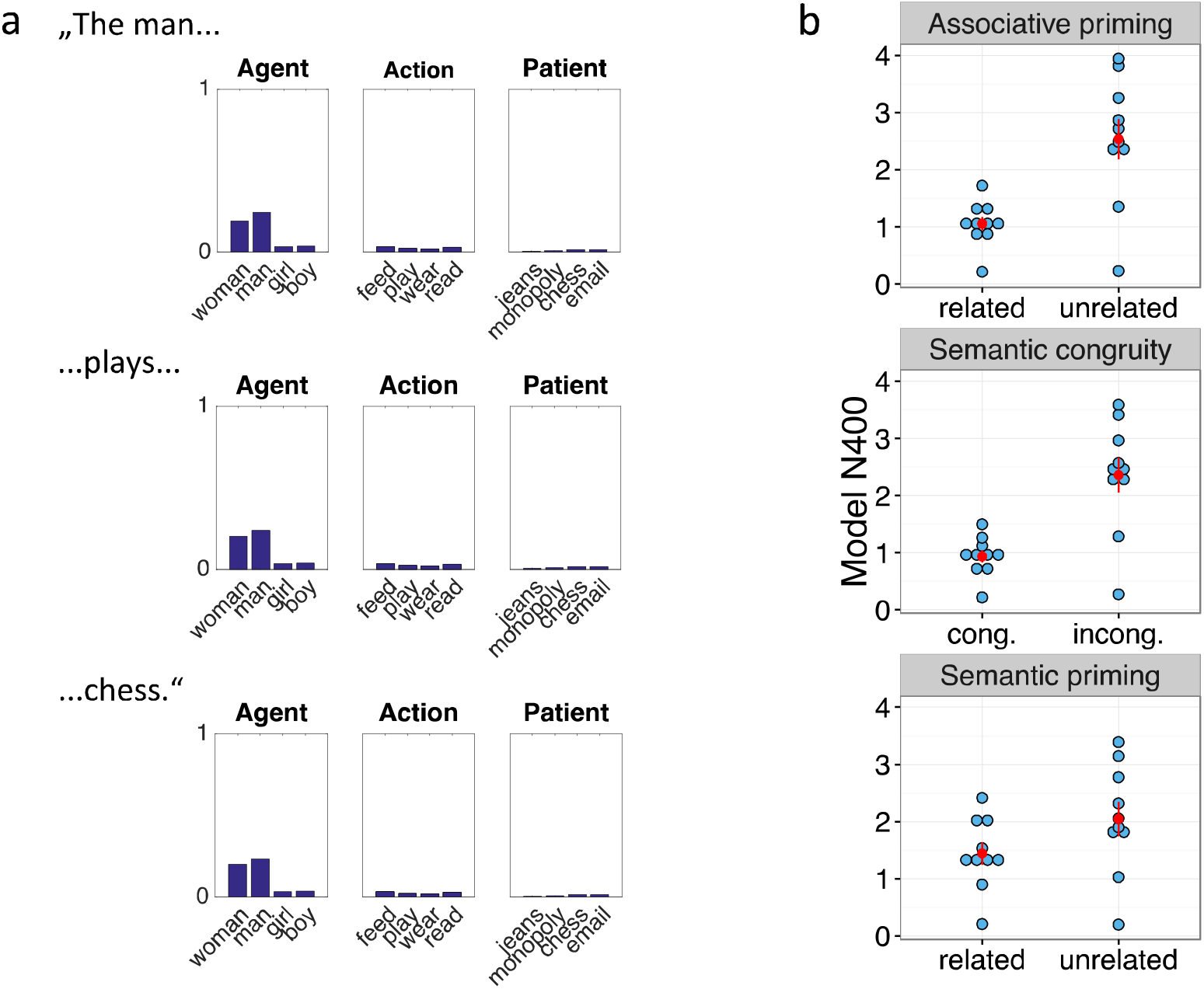
Comprehension performance and semantic update effects at a very early stage in training. Cong., congruent; incong., incongruent. a. Activation of selected output units while the model is presented with the sentence “The man plays chess.”. It can be seen that the model fails to activate the corresponding units at the output layer. The only thing that it has apparently learned at this point is which concepts correspond to possible agents, and it activates those in a way that is sensitive to their base rate frequencies (in the model’s environment, woman and man are more frequent than girl and boy; see online methods), and with a beginning tendency to activate the correct agent (“man”) most. b. Even at this low level of performance, there are robust effects of associative priming (t_1(9)_ = 6.12, p = .00018, d = 1.94, 95% CI[.93, 2.03], t_2(9)_ = 7.31, p < .0001, d = 2.31, 95% CI[1.02, 1.94], top), semantic congruity in sentences (t_1(9)_ = 6.85, p < .0001, d = 2.16, 95% CI [.95, 1.90]; t_2(9)_ = 5.74, p = .00028, d = 1.81, 95% CI [.86, 1.99], middle), and semantic priming (t_1(9)_ = 5.39, p = .0004, d = 1.70, 95% CI[.35, .85], t_2(9)_ = 3.79, p = .0043, d = 1.20, 95% CI[24, .96], bottom), on the size of the semantic update, the model’s N400 correlate. Each blue dot represents the results for one independent run of the model, averaged across items per condition; the red dots represent the means for each condition, and red error bars represent +/− SEM.

#### The relationship between activation update and adaptation in a predictive system

The change induced by the next incoming word that we suggest underlies N400 amplitudes can be seen as reflecting the ‘error’ (difference or divergence) between the model’s implicit probability estimate based on the previous word, and the updated estimate based on the next word in the sentence (see *online methods* for details). If the estimate after word *n* is viewed as a *prediction*, then this difference can be viewed as a kind of prediction error. It is often assumed that learning is based on such temporal difference or prediction errors^49–51^ so that if N400 amplitudes reflect the update of a probabilistic representation of meaning, then larger N400s should be related to greater adaptation, i.e., larger adjustments to future estimates. Here we implement this idea, using the semantic update to drive learning: The SG layer activation at the next word serves as the target for the SG layer activation at the current word, so that the error signal that we back-propagate through the network to drive the adaptation of connection weights after each presented word becomes the difference in SG layer activation between the current and the next word, i.e. SG_n+1_ – SG_n_ (see *online methods* for more details). Importantly, this allows the model to learn just from listening or reading, when no separate event description is provided. We then used this approach to simulate the finding that the effect of semantic incongruity on N400s is reduced by repetition: the first presentation of an incongruent completion, which induces larger semantic update compared to a congruent completion, leads to stronger adaptation, as reflected in a larger reduction in the N400 during a delayed repetition compared to the congruent continuation^52^.

To simulate the observed interaction between repetition and semantic incongruity, we presented a set of congruent and incongruent sentences a first time, adapting the weights in the update network using the temporal difference signal on the SG layer to drive learning. We then presented all sentences a second time. Using this approach, we captured the greater reduction in the N400 with repetition of incongruent compared to congruent sentence completions (Fig. 8 and Supplementary Fig. 9). Notably, the summed magnitude of the signal that drives learning corresponds exactly to our N400 correlate, highlighting the relationship between semantic update, prediction error, and experience-driven learning. Thus, our account predicts that in general, larger N400s should induce stronger adaptation. Though further investigation is needed, there is some evidence consistent with this prediction: larger N400s to single word presentations during a study phase have been shown to predict enhanced implicit memory (measured by stem completion in the absence of explicit memory) during test^53^.

**Figure 8.**
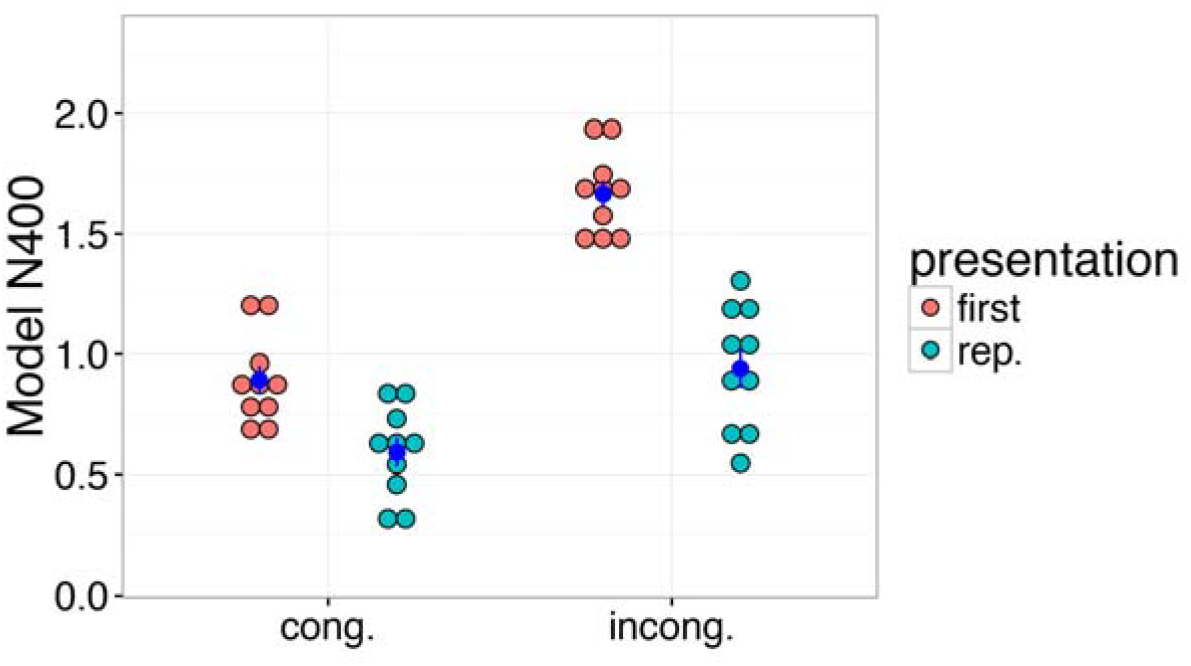
Simulation of the interaction between delayed repetition and semantic incongruity. Cong., congruent; incong., incongruent; rep., repeated. Each red or green dot represents the results for one independent run of the model, averaged across items per condition; the blue dots represent the means for each condition, and blue error bars represent +/− SEM. There were significant main effects of congruity. F_1_(1,9) = 214.13, p < .0001, 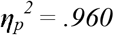, F_2(1,9)_ = 115.66, p < .0001, 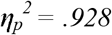 and repetition, F_1(1,9)_ = 48.47, p < .0001, 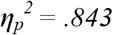, F_2(1,9)_ = 109.78, p < .0001, 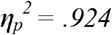, and a significant interaction between both factors, F_1(1,9)_ = 83.30, p < .0001, 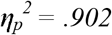, F_2(1,9)_ = 120.86, p < .0001, 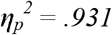; post-hoc comparisons showed that even though the repetition effect was larger for incongruent as compared to congruent sentence completions, it was significant in both conditions, t_1(9)_ = 4.21, p = .0046, d = 1.33, 95% CI[.14, .46], t_2(9)_ = 6.90, p < .0001, d = 2.18, 95% CI[.20, .40], for the congruent completions, and t_1(9)_ = 8.78, p < .0001, d = 2.78, 95% CI[.54, .91], t_2(9)_ = 12.02, p < .0001, d = 3.80, 95% CI[.59, .86] for the incongruent completions.

## Discussion

The N400 ERP component is widely used to investigate the neurocognitive processes underlying the processing of meaning in language. However, the component’s functional basis continues to be actively debated^2^. In the simulations presented above, we have shown that an implemented computational model of language comprehension, the *Sentence Gestalt* model, can provide a unified account that captures a wide range of findings (Table 1). The model treats N400 amplitudes as indexing the change induced by an incoming word in an implicit probabilistic representation of meaning. Our account has some similarities to existing descriptive accounts of the N400’s functional basis, while differing in important ways from these theories. Below we explain how the model’s distinctive characteristics, as described in the introduction, contribute to its ability to account for the findings we have considered.

First, our model does not assume separate stages for lexical access (retrieval of word meaning) and for subsequent integration of word meanings into a compositional representation. This is crucial because the two most prominent competing theories of the N400’s functional basis suggest that N400 amplitudes reflect either lexical access^3^ or integration (also referred to as unification) into a compositional (sometimes called combinatorial) representation of the meaning of the sentence^6,7^. In the SG model, incoming stimuli instead serve as ‘cues to meaning’^54^ which automatically change an activation pattern that implicitly represents estimates of conditional probabilities of all aspects of meaning construed as a description of the event or situation described by the sentence. Our account is similar to the lexical access perspective in that the process is assumed to be fast, automatic, and implicit, but differs from this view in that the resulting activation pattern represents not just the currently incoming word but an updated implicit representation of the event described by the sequence of words it has received as input. In this regard our account is similar to the integration view in that the resulting activation state is assumed to represent all aspects of the described event (including – though this aspect is not currently implemented – input from other modalities), though it differs from such accounts in avoiding a commitment to explicit compositional representation. Our perspective seems in line with a recent comprehensive review on the N400 ERP component^2^ which concluded that the N400 might best be understood as a “temporally delimited electrical snapshot of the intersection of a feedforward flow of stimulus-driven activity with a state of the distributed, dynamically active neural landscape that is semantic memory.” (p. 641). Crucially, the SG model provides a computationally explicit account of the nature and role of this distributed activation state and how it changes through stimulus-driven activity as meaning is dynamically constructed during comprehension. As noted in the introduction, the current implementation of the model focuses on sentences describing events, but the theory applies to sentence comprehension in general, and the model could learn to understand other types of sentences (including statives and questions) if trained with the appropriate materials. The model uses event probability information together with word order information to build a meaning representation instead of relying on syntax to build a linguistic representation in which word meanings are placed into syntactically specified slots. The model may override syntactic conventions when event probability information is very strong or it may enter a state of uncertainty when syntactic and event probability information conflict. These aspects of the model, taken together with the statistics of its relevant experiences with sentences and the events they describe, underlie the pattern of responses it exhibits when presented with sentences containing reversal anomalies (Fig. 2i, Fig. 3, & Supplementary Fig. S3) or violations of word order conventions (Fig. 4), allowing it to explain the absence of N400 responses when listeners or readers comprehend such sentences.

Second, the model’s representations result from a learning process and thus depend on the statistical regularities in the model’s environment as well the amount of training the model has received. These features allow the model to account for the pattern of N400 effects across development (Fig. 6) including N400 effects while behavioral performance is still near chance (Fig. 7) as well as the influence of delayed repetition on N400 congruity effects (Fig. 8).

Third, the model updates its activation pattern upon the presentation of a word, whether or not it actually occurs in a sentence, allowing it to capture N400 effects for single words (i.e., frequency effects; see Fig. 2e) and words presented in pairs (influences of repetition, Fig. 2h, semantic priming, Fig. 2f, and associative priming, Fig. 2g) as well as words presented in a sentence context (influences of semantic congruity, Fig. 2a, cloze probability, Fig. 2b, position in the sentence, Fig. 2c, semantically related incongruity, Fig. 2d, and no influence of constraint for unexpected endings, Fig. 5).

Fourth, the N400 as captured by the model is proposed to characterize one specific aspect of language comprehension, namely the automatic stimulus-driven update of an initial implicit representation of the event or situation described by the sentence. This characterization is in line with evidence for the N400’s anatomical localization in regions involved in semantic representation such as the medial temporal gyrus (MTG^3^) and anterior medial temporal lobe (AMTL^55,56^). The processes underlying the N400 may thus correspond to a type of language processing that others have characterized as shallow, incomplete, underspecified^57^ or plausibility based^38^, and sometimes described as “good enough”^58^. This kind of processing may be preserved in patients with lesions to frontal cortex (specifically left inferior prefrontal cortex, BA47)^59,60^. Thus, activity in temporal lobe regions MTG and AMTL may correspond to immediate, automatic, and implicit aspects of sentence processing as captured by the SG model. In contrast, the left, inferior frontal cortex has been proposed to support control processes in comprehension that are required only when processing demands are high^61,62^ such as in syntactically complex sentences^59^ and sentences which require selection among competing alternative interpretations^63^. This also holds for situations where event probability constraints must be overcome to correctly represent the specific meaning of certain sentences, including those describing implausible events and sentences with unusual structure such as reversal anomalies and garden-path sentences. These aspects of language comprehension may be reflected in other ERP components as discussed below. Delineating the boundary between situations where the automatic and heuristic processes captured by the SG model are sufficient for language understanding and situations requiring more controlled processes to accurately represent sentence meaning is an important topic for future investigation.

The pattern of activation in the model’s Sentence Gestalt (SG) layer latently predicts the attributes of the entire event described by a sentence, capturing base-rate probabilities (before sentence processing begins) and adjusting this pattern of activation as each word of the sentence is presented. While in the current implementation of the model, inputs are presented over a series of discrete time steps corresponding to each successive word in the sentence, this is just a simplification for tractability. We assume that in reality, the adjustment of the semantic activation occurs continuously in time as auditory or visual language input is processed, so that the earliest arriving information about a word (whether auditory or visual) immediately influences the evolving SG representation^64^. This assumption is in line with the finding that N400 effects in spoken language comprehension often begin to emerge before the spoken word has become acoustically unique^65,66^. It is important to underline the point that this kind of prediction does not refer to the active and intentional prediction of specific items but rather to a latent or implicit state such that the model (and presumably the brain) becomes tuned through experience to anticipate likely upcoming input and to respond to it with little additional change. This entails that semantic activation changes induced by new incoming input as revealed in the N400 reflect the discrepancy between probabilistically anticipated and encountered information about aspects of the state of the world conveyed by the sentence and at the same time correspond to the learning signal driving adaptation of connection-based knowledge representations. In this sense, our approach, first introduced almost 30 years ago, anticipates aspects of Bayesian approaches to understanding the dynamics of neural activity patterns in the brain^50,67^. Our simulations suggest that the semantic system may not represent probabilities of aspects of meaning explicitly but rather uses a summary representation that implicitly represents estimates of these probabilities, supporting explicit estimates when queried and becoming more and more efficient as learning progresses.

Recently, other studies have also begun to link the N400 to computational models. Most of these have concentrated on words presented singly or after a preceding prime, and therefore do not address processing in a sentence context^16–18,68^. Two modeling studies focus on sentence processing. One of these studies observed a correlation between N400s and word surprisal as estimated by a simple recurrent network (SRN) trained to predict the next word based on the preceding context^31^. Because this SRN’s predictions generalize across contexts and are mediated by a similarity-based internal representation, it can potentially account for effects of semantic similarity on word surprisal, and would thus share some predictions with the SG model. However, an account of N400s in terms of word surprisal faces some difficulties. To demonstrate this, we trained an SRN on the same training corpus as the SG model and repeated some of the critical simulations with this SRN (Fig. 9 and Supplementary Fig. 10; see *online methods* for details).

**Figure 9.**
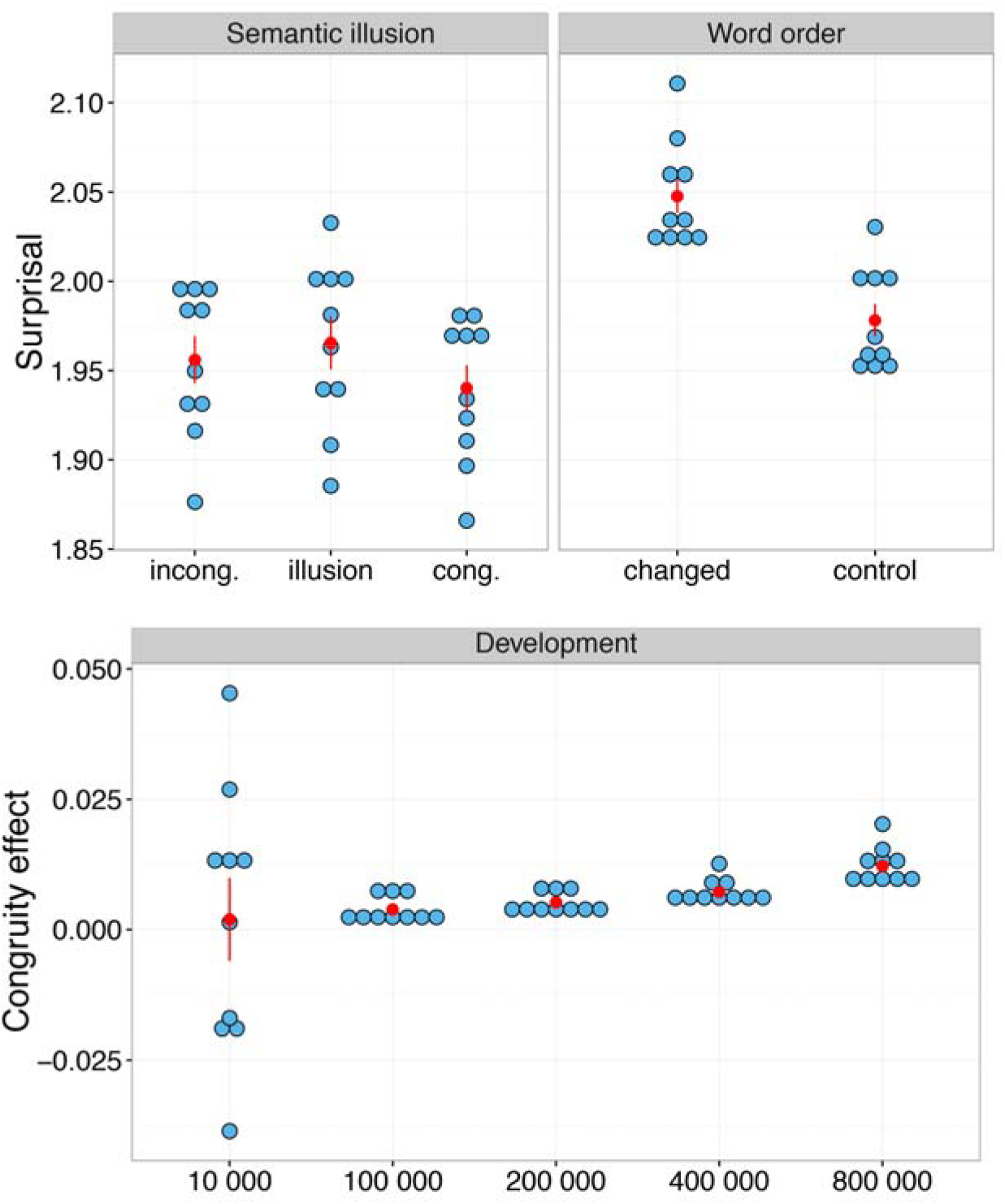
Simulation results from a simple recurrent network model (SRN) trained to predict the next word based on the preceding context. Each blue dot represents the results for one independent run of the model, averaged across items per condition; the red dots represent the means for each condition, and red error bars represent +/− SEM. Top left, reversal anomaly: t_1(9)_ = 4.55, p = .0042, d = 1.44, 95% CI[.013, .038], t_2(7)_ = 7.83, p = .0003, d = 2.77, 95% CI [.018, .033] for the comparison between congruent and reversal anomaly; t_1(9)_ = 12.28, p < .0001, d = 3.87, 95% CI[013, .019], t_2(7)_ = 2.98, p = .062, d = 1.05, 95% CI[003, .028] for the comparison between congruent and incongruent condition; t_1(9)_ = 1.52, p = .49, d = .48, 95% CI[-.005, .024], t_2(9)_ = 1.57, p = .48, d = .55, 95% CI[-.005, .024] for the comparison between incongruent and reversal anomaly condition. Top right, word order: t_1(9)_ = 29.78, p < .0001, d = 9.42, 95% CI[064, .075]; t_2(15)_ = 6.73, p < .0001, d = 1.68, 95% CI [.048, .092]. Bottom, congruity effect on surprisal as a function of the number of sentences the model has been exposed to: t_1(9_) =.26, p = 1.0, d = .082, 95% CI [-.015, .019]; t_2(9)_ = .15, p = 1.0, d = .048, 95% CI[-.027, .031] for the comparison between 10 000 and 100 000 sentences; t_1(9)_ = 6.74, p = .0003, d = 2.13, 95% CI[.0009, .0019]; t_2(9)_ = 1.08, p = 1.0, d = .34, 95% CI[-.0015, .0043] for the comparison between 100 000 and 200 000 sentences; t_1(9)_ = 7.45, p = .00015, d = 2.36, 95% CI[0014, .0026]; t_2(9)_ = 1.78, p = .44, d = .56, 95% CI[-.0005, .0045] for the comparison between 200 000 and 400 000sentences; t_1(9)_ = 10.73, p < .0001, d = 3.39, 95% CI[.0039, .0060]; t_2(9)_ = 1.93, p = .36, d = .61, 95% CI[-.0008, 011] for the comparison between 400 000 and 800 000 sentences.

First, word surprisal reflects both semantic and syntactic expectation violations, while the N400 is specific to semantic expectations as described above. Indeed, while SU in the SG model was insensitive to changes in word order (Fig. 4 and Supplementary Fig. 5), surprisal in the SRN was significantly larger for changed as compared to normal word order (see Fig. 9 and Supplementary Fig. 10). The lack of specificity of the word surprisal measure converges with the finding that the correlation between surprisal in the SRN and N400 observed in the above mentioned study^31^ was observed only for content words; the SRN surprisal measure when calculated over grammatical function words did not correlate with the N400 responses observed on these words.

Furthermore, the SRN did not account for the decrease of N400 effects with age, showing instead a slight increase with additional training (see Fig. 9 and Supplementary Fig. 10). This is because surprisal is measured in terms of the estimates of word probabilities, which become sharper as learning progresses. Finally, the SRN did not produce the small N400 in reversal anomalies: When presented with “At breakfast, the eggs *eat*…”, word surprisal was large, numerically even larger than an incongruent continuation (see Fig. 9 and Supplementary Fig. 10) while semantic update in the SG model shows only a very slight increase, in line with N400 data^37^ (see also Supplementary Fig. 11 and the accompanying text for relevant results from an SRN trained on a natural corpus by S. Frank, personal communication).

The other sentence-level model focuses specifically on reversal anomalies, assuming separate stages of lexical retrieval and semantic integration^39^. This retrieval-integration model is computationally explicit while following aspects of the classical framework for language processing, in which there is thought to be a distinct lexical-semantic processing module in which spreading activation can occur among related items, prior to integrating the retrieved word meanings into a compositional representation of sentence meaning^10,69^. The retrieval-integration model makes the further assumption that reversal anomalies such as ‘for breakfast the eggs would *eat*’ or ‘the fox on the poacher *hunted*’ must produce a large update in the representation of sentence meaning. This is because the sentences appear to describe events in which eggs are agents engaged in the act of eating and the fox is an agent engaged in the act of hunting the poacher. In this model, change in lexical activation (which is small in reversal anomalies due to priming, e.g. from *fox* and *poacher* to *hunting*) is linked to the N400; the change in activation representing sentence meaning is assigned to the later, P600 ERP component.

As discussed above, our model accounts for the small size of the N400 in reversal anomalies without separate mechanisms for lexical access and semantic interpretation, and addresses a wide range of N400 effects that traditional accounts would ascribe either to lexical access or to subsequent semantic integration. Crucially, our model accounts for the absence of an N400 effect in reversal anomalies because it takes both syntactic and semantic cues into account and can favor event statistics or remain uncertain and inconclusive when there is a conflict between different constraints.

While both the retrieval-integration model and the SG model account for the small N400 effects in reversal anomalies, the SG model does so within the context of a more complete account of the factors that do and do not influence the N400, while the retrieval-integration model has yet to be extended beyond accounting for a subset of the relevant findings. Further research will be required to determine whether the retrieval-integration model can encompass the range of N400 findings encompassed by the SG model.

The functional basis of the P600 component – which is increased in reversal anomalies - is not addressed by our model and requires further investigation to be more fully understood. It is true that P600 responses have been observed to a wide range of linguistic violations and irregularities, including reversal anomalies^37,40,70^, syntactic violations^42^, and garden path sentences^71^, as well as pragmatic processes such as the comprehension of irony (see review^72^). This has been taken to suggest that the P600 might reflect combinatorial aspects of language processing, either related to syntax^42^ or to semantic integration as assumed in the retrieval-integration model^39^. There is, however, an alternative perspective, in which the P600 is not treated as specific to language processing (either syntactic processes or semantic integration) *per se*, but to a more general process that may be associated with more conscious, deliberate, and effortful aspects of processing. Several researchers have pointed out that the P600 shares properties with the P3^73,74^ which is elicited by the occurrence of oddball stimuli (such as a rare high tone among much more frequent low tones), with the component’s latency depending on stimulus complexity. This component is thought to signal an explicit surprise response and a corresponding update in working memory^75^. This P600-as-P3 perspective naturally explains the observed sensitivity of P600 effects to task demands and attentional focus. Indeed, P600 effects are strongly reduced or absent when there is no active task or when the task is unrelated to the linguistic violation^76^. In contrast, N400 effects can be obtained during passive reading and even during unconscious processing such as within the attentional blink^77^. Thus, from this view, the P600 differs from the N400 in two ways. It belongs to a component family that responds to a wider range of expectation violations while the N400 is specific to the formation of a representation of meaning. Further, the N400 may reflect an automatic and implicit process which can result in representations that are plausibility based, possibly underspecified^57^, and not completely accurate representations of the linguistic input as determined by grammatical conventions^58^. On the other hand, the P600 may be associated with processes requiring a higher level of control and attention. These processes may be affected by factors beyond those affecting the semantic update process underlying N400 and may contribute to resolving situations in which there is a conflict between event probability and syntactic cues. These issues should be investigated in future research to be more fully understood. These investigations should include a consideration of individual differences in how complex syntactic constraints are combined with event probability considerations in order to make sense of linguistic input^78,79^.

As discussed above, there is indeed considerable evidence that representations formed during language comprehension are sometimes influenced by event probabilities. To directly investigate the claim that the N400 corresponds to the formation of such representations it would seem necessary to combine N400 measurements with comprehension questions, probing, for example, the comprehension of role-reversed sentences (e.g., “The dog was bitten by the *man*.”)^58^. Further consideration of responses to questions about such sentences, which have been shown to lead to a high rate of role assignment errors, would allow direct examination of the co-variation between N400 amplitudes and event probability based comprehension^3^. Another aspect which should be addressed in future research is the parametric variation of factors contributing to the effects obtained in reversal anomaly sentences. Relevant dimensions seem to be the relative frequency of both event participants to be an agent in an event, the relative likelihood to be an agent versus patient in events involving both participants, the relative likelihood to be able to perform the relevant action, the reliability of structural cues such as word order, etc. These factors should be systematically manipulated in model environments as well as in materials selected for empirical experiments, and experimental materials should be selected so that the values on the above-mentioned dimensions within the participant’s natural language environment correspond to the respective values in the model’s environment.

In general, the current work opens up an opportunity for extensive further investigations, addressing a wide range of behavioral as well as neural aspects of language processing. One key finding that needs to be addressed in future work is the finding that N400s were influenced by categorical relationship (i.e., semantic priming, see Fig. 2f) while being unaffected by sentence truth, at least in negated statements: The N400 is equally small in the false sentence “A robin is not a bird” and the true sentence “A robin is a bird”, and is equally large in the true sentence “A robin is not a vehicle” and the false sentence “A robin is a vehicle”^80^. It is important to note that sentence truth is not the same as expected sentence meaning, and that to understand the influence of negation on meaning expectations, one needs to take into account the pragmatics of negation^81,82^. Specifically, negation is typically used to deny a supposition, and in the absence of discourse context, this supposition must be grounded in general knowledge^81^. Thus, when used in short and isolated sentences, negation is typically used to deny something that is part of an invoked schema (e.g., “a whale is not a fish”). “Robin” does not invoke a schema which includes semantic features of “vehicle” so that “A robin is not a vehicle” is not an expected sentence meaning, even though it is true. On the other hand, “robin” does invoke a schema which includes semantic features of “bird” so that something that is part of the schema of “bird” might be expected to be denied (e.g., “A robin is not a bird that flies south during winter” is fine). Follow-ups taking the pragmatics of negation into account and providing more context showed that N400s are indeed modulated by sentence truth^82^ and plausibility^81^. Our model currently has no experience with negation and its pragmatics, but this could be incorporated in an extension of the model, allowing further research to investigate whether the pattern of semantic update seen in such sentences can be captured. A further observation that should be addressed by future simulations is the finding that discourse semantics can influence the N400 over and above the local sentence context^83^. This includes discourse contexts in which, for example, peanuts are fictively treated as human-like, completely reshaping expectations for plausible continuations of a sentence about the peanuts in question^84^. Addressing this issue will be crucial for determining how far the model can go in capturing N400 effects beyond local association and priming. Doing so will require extension of the model because it currently just processes one sentence at a time. However, the Story Gestalt model, which directly built upon the SG model already constitutes one possible approach towards such an extension in that it processes several sentences forming a coherent text^85^. Moreover, recent deep learning models trained to read natural language documents are based on a similar approach as the SG model, with models learning to answer questions posed about the document’s content^86^. These models have limitations in their current form. Nevertheless, they provide a starting point for addressing this important issue.

Finally, it remains to be explored how well the SG model can address behavioral measures of sentence processing. Given the extensive evidence reviewed above that the update of an implicit probabilistic representation of meaning is only one of the processes that occurs during language processing, it seems likely that a full account of overt behavioral responses would require a fuller model capturing these other processes. The beauty of ERPs is that they may index distinct aspects of these processes, and can thus speak to their neurocognitive reality even though several such processes might jointly influence a specific behavioral measure. To fully address behavior, the model will likely need to be integrated into a more complete account of the neuro-mechanistic processes that take place during language processing, including the more controlled and attention-related processes that may underlie the P600. In addition, the model’s query language and training corpus will need to be extended to address this issue, as well as the full range of relevant neurocognitive phenomena, including other ERP components (e.g., orthographic and syntactic ERPs) and signals that have been detected using other measurement modalities^62,87^.

While extending the model will be worthwhile, it nevertheless makes a useful contribution to understanding the brain processes underlying language comprehension in its current simple form. The model’s successes in capturing a diverse body of empirically observed neural responses suggest that the principles of semantic representation and processing it embodies may capture essential aspects of human language comprehension.

## Online Methods

We begin by describing the implicit probabilistic theory of meaning underlying the Sentence Gestalt (SG) model and relate the updates in the model to other probabilistic measures of surprise. Next we describe the new semantic update driven learning rule used in simulating the reduction in the incongruity effect due to repetition. We then provide details on the model’s training environment as well as the protocols used for training the model and for the simulations of empirical findings. Finally, we describe simulations conducted with an SRN. Figure 1 in the main text presents the SG network architecture and the processing flow in the model.

### Implicit probabilistic theory of meaning

The theory of meaning embodied in the Sentence Gestalt model holds that sentences constrain an implicit probabilistic representation of the meanings speakers intend to convey through these sentences. The representation is implicit in that no specific form for the representation is prescribed, nor are - in the general form of the theory - specific bounds set on the content of the representation of meaning. In any specific implementation of the theory, the content of the representation of meaning is prescribed by the range of possible probes and queries, which in the case of our implementation correspond to the vectors encoding the pairs of thematic roles and their fillers of described events. Sentences are viewed as conveying information about situations or events, and a representation of meaning is treated as a representation that provides the comprehender with a basis for estimating the probabilities of aspects of the situation or event the sentence describes. To capture this we characterize the ensemble of aspects as an ensemble of queries about the event, with each query associated with an ensemble of possible responses. The query-answer form is used instead of directly providing the complete event description at the output layer to keep the set of probes and fillers more open-ended and to suggest the broader framework that the task of sentence comprehension consists in building internal representations that can be used as a basis to respond to probes^13^. In the general form of the theory, the queries could range widely in nature and scope, encompassing, for example, whatever the comprehender should expect to observe via any sense modality or subsequent linguistic input, given the input received so far. This includes queries supporting expectations concerning the content of stative sentences (e.g., “Her hair is red.”) and probes supporting the anticipation of aspects of meaning of questions or commands based on representations concerning the current state of knowledge and intentions of the speaker, etc. For instance, the question “Where is the bathroom?” communicates that the speaker would like to know the location of the bathroom, and the command “Please close the door.” communicates that the speaker wants the listener to close the door. Thus, it is important to note that even though the current implementation focuses on sentences describing events, the theory is thought of as applying to language comprehension in general. In implementations to date, at least four different query formats have been considered^14,15,88^, including a natural language-based question and answer format (Fincham & McClelland, 1997, Abstract). Queries may also vary in their probability of being posed (hereafter called *demand probability*), and the correct answer to a particular query may be uncertain, since sentences may be ambiguous, vague or incomplete. An important aspect of the theory that receives little attention in many other theories of sentence comprehension is that aspects of meaning can often be estimated without being explicitly described in a sentence, due to knowledge acquired through past experience^14^. If events involving cutting steak usually involve a knife, the knife would be understood, even without ever having been explicitly mentioned in a sentence.

The theory envisions that sentences are uttered in situations where information about the expected responses to a probabilistic sample of queries is often available to constrain learning about the meaning of the sentence. When such information is available, the learner is thought to be (implicitly) engaged in attempting to use the representation derived from listening to the sentence to anticipate the expected responses to these queries and to use the actual responses provided with the queries to bring the estimates of the probabilities of these responses in line with their probabilities in the environment. This process is thought to occur in real time as the sentence unfolds; for simplicity it is modeled as occurring word by word as the sentence is heard.

As an example, consider the sequence of words ‘The man eats’ and the query, ‘What does he eat’? What the theory assumes is that the environment specifies a probability distribution over the possible answers to this and many other questions, and the goal of learning is to form a representation that allows the comprehender to match this probability distribution.

More formally, the learning environment is treated as producing sentence-event-description pairs according to a probabilistic generative model. The sentence consists of a sequence of words, while the event-description consists of a set of queries and associated responses. Each such pair is called an *example*. The words in the sentence are presented to the neural network in sequence, and after each word, the system can be probed for its response to each query, which is conditional on the words presented so far (we use *W_n_* to denote the sequence of words up to and including word *n*). The goal of learning is to minimize the expected value over the distribution of examples of a probabilistic measure (the Kullback-Leibler divergence, *D_KL_*) of the difference between the distribution of probabilities *p* over possible responses *r* to each possible query and the model’s estimates *ρ* of the distribution of these probabilities, summed over all of the queries *q* occurring after each word, and over all of the words in the sentence. In this sum, the contribution of each query is weighed by its demand probability conditional on the words seen so far, represented *p*(*q*|*w_n_*). We call this the *expected value E of the summed divergence measure*, written as:

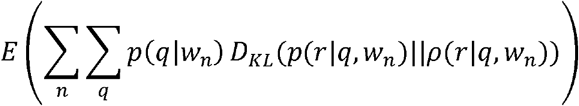

In this expression the divergence for each query, *D_KL_*(*p*(*r|q,W_n_*)/*ρ*(*r|q,W_n_*)), is given by

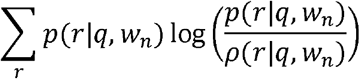

It is useful to view each combination of a query *q* and sequence of words *W_n_* as a context, henceforth called *C*. The sequence of words ‘the man eats’ and the query ‘what does he eat?’ is an example of one such context. To simplify our notation, we will consider each combination of *q* and *W_n_* as a context *C*, so that the divergence in context *C*, written *D_KL_*(*C*), is 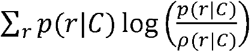. Note that *D_KL_*(*C*) equals 0 when the estimates match the probabilities (that is, when *p*(*r|C*) = *ρ*(*r|C*) for all *r*) in context *C*, since log(x/x) = log(1) = 0. Furthermore, the expected value of the summed divergence measure is 0 if the estimates match the probabilities for all *C*.

Because the real learning environment is rich and probabilistic, the number of possible sentences that may occur in the environment is indefinite, and it would not in general be possible to represent the estimates of the conditional probabilities explicitly (e.g. by listing them in a table). A neural network solves this problem by providing a mechanism that can process any sequence of words and associated queries that are within the scope of its environment, allowing it to generate appropriate estimates in response to queries about sentences it has never seen before^14^.

Learning occurs from observed examples by stochastic gradient descent: A training example consisting of a sentence and a corresponding set of query-response pairs is drawn from the environment. Then, after each word of the sentence is presented, each of the queries is presented along with the response that is paired with it in the example. This response is treated as the target for learning, and the model adjusts its weights to increase its probability of giving this response under these circumstances. This procedure tends to minimize the expected value of the summed divergence measure over the environment, though the model’s estimates will vary around the true values in practice as long as a non-zero learning rate is used. In that case the network will be sensitive to recent history and can gradually change its estimates if there is a shift in the probabilities of events in the environment.

### The implemented query-answer format and standard network learning rule

In the implementation of the model used here, the queries presented with a given training example can be seen as questions about attributes of the possible fillers of each of a set of possible roles in the event described by the sentence. There is a probe for each role, which can be seen as specifying a set of queries, one for each of the possible attributes of the filler of the role in the event. For example, the probe for the agent role can be thought of as asking, in parallel, a set of binary yes-no questions, one about each of several attributes or features *f* of the agent of the sentence, with the possible responses to the question being 1 (for yes the feature is present) or 0 (the feature is not present). For example, one of the features specifies whether or not the role filler is male. Letting *p*(*v|f,C*) represent the probability that the feature has the value *v* in context *C* (where now context corresponds to the role being probed in the training example after the *n*th word in the sentence has been presented), the divergence can be written as 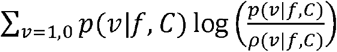. Writing the terms of the sum explicitly, this becomes 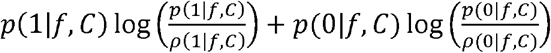. Using the fact that the two possible answers are mutually exclusive and exhaustive, the two probabilities must sum to 1, so that *p*(0|*f,C*) = 1 – *p*(1|*f,C*); and similarly, *ρ*(0|*f,C*) = 1 – *ρ*(1|*f,C*). Writing *p*(*f|C*) as shorthand for *p*(1|*f,C*) and *ρ*(*f|C*) for *ρ*(1|*f,C*), and using the fact that log(*a/b*) = log(*a*) – log(*b*) for all *a,b*, the expression for *D_KL_*(*f,C*) becomes

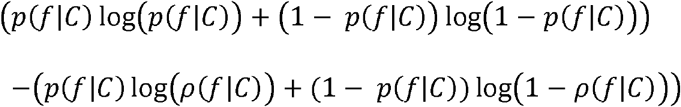

The first part of this expression contains only environmental probabilities and is constant, so that minimizing the expression as a whole is equivalent to minimizing the second part, called the *cross-entropy CE*(*f,C*) between the true and the estimated probability that the value of feature *f* = 1 in context *C*:

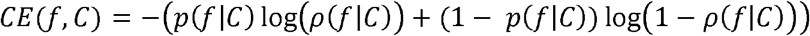

The goal of learning is then to minimize the sum of this quantity across all features and situations.

The actual value of the feature for a particular role in a randomly sampled training example *e* is either 1 (the filler of the role has the feature) or 0 (the filler does not have the feature). This actual value is the target value used in training, and is represented as *t*(*f|C_e_*), where we use *C_e_* to denote the specific instance of this context in the training example (note that the value of a feature depends on the probed role in the training example, but stays constant throughout the processing of each of the words in the example sentence). The activation *a* of a unit in the query network in context *C_e_, a*(*f|C_e_*), corresponds to the network’s estimate of the probability that the value of this feature is 1 in the given context; we use *a* instead of *ρ* to call attention to the fact that the probability estimates are represented by unit activations. The *cross-entropy* between the target value for the feature and the probability estimate produced by the network in response to the given query after word *n* then becomes:

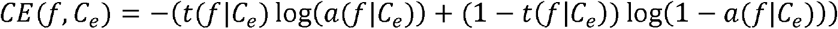

To see why this expression represents a sample that can be used to estimate *CE*(*f,C*) above, it is useful to recall that the value of a feature in a given context varies probabilistically across training examples presenting this same context. For example, for the context ‘the man eats …’, the value of a feature of the filler of the patient role can vary from case to case. Over the ensemble of training examples, the probability that *t*(*f|C_e_*) = 1 corresponds to *p*(*f|C*), so that the expected value of *t*(*f|C_e_*) over a set of such training examples will be *p*(*f|C*), and the average value of *CE*(*f,C_e_*) over such instances will approximate *CE*(*f,C*).

Now, the network uses units whose activation *a* is given by the logistic function of its net input, such that *a* = 1/(1 + *e*^−*net*^), where the net input is the sum of the weighted influences of other units projecting to the unit in question, plus its bias term. As has long been known^89^, the negative of the gradient of this cross-entropy measure with respect to the net input to the unit is simply *t*(*f|C_e_*) – *a*(*f|C_e_*). This is the signal back-propagated through the network for each feature in each context during standard network training (see section *simulation details/ training protocol* for more detail).

### Probabilistic measures of the surprise produced by the occurrence of a word in a sentence

Others have proposed probabilistic measures of the surprise produced by perceptual or linguistic inputs^19,30^. In the framework of our approach to the characterization of sentence meaning, we adapt one of these proposals^19^, and use it to propose measures of three slightly different conceptions of surprise: The normative surprise, the subjective explicit surprise, and the implicit surprise – the last of which corresponds closely to the measure we use to model the N400.

We define the normative surprise (NS) resulting from the occurrence of the nth word in a sentence *s* as the KL divergence between the environmentally determined distribution of responses *r* to the set of demand-weighted queries *q* before and after the occurrence of word *W_n_*:

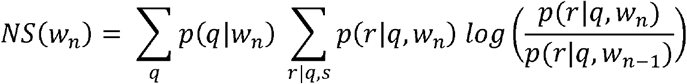

If one knew the true probabilities, one could calculate the normative surprise and attribute it to an ideal observer. In the case where the queries are binary questions about features as in the implemented version of the SG model this expression becomes:

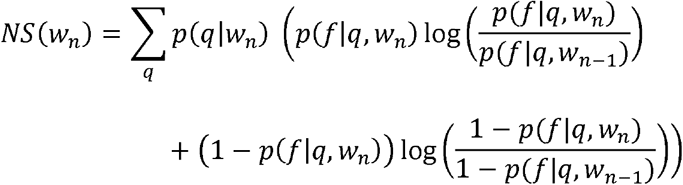

To keep this expression simple, we treat *q* as ranging over the features of the fillers of all of the probed roles in the sentence.

The explicit subjective surprise ESS treats a human participant or model thereof as relying on subjective estimates of the distribution of responses to the set of demand-weighted queries. In the model these are provided by the activations *a* of the output units corresponding to each feature:

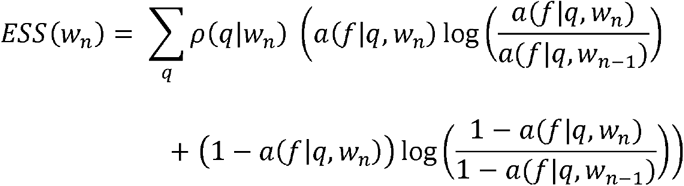

Our third measure, the implicit surprise (IS) is a probabilistically interpretable measure of the change in the pattern of activation over the learned internal meaning representation (corresponding to the SG layer in the model). Since the unit activations are constrained to lie in the interval between 0 and 1, they can be viewed intuitively as representing estimates of probabilities of implicit underlying meaning dimensions or *microfeatures*^90^ that together constrain the model’s estimates of the explicit feature probabilities. In this case we can define the implicit surprise as the summed KL divergence between these implicit feature probabilities before and after the occurrence of word *n*, using *a_i_* to represent the estimate of the probability that the feature characterizes the meaning of the sentence and (1 – *a_i_*) to represent the negation of this probability:

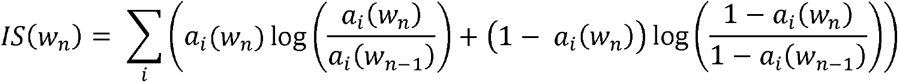

The actual measure we use for the semantic update (SU) as defined in the main text is similar to the above measure in being a measure of the difference or divergence between the activation at word *n* and word *n*−1, summed over the units in the SG layer:

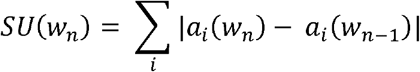

The SU and IS are highly correlated and have the same minimum (both measures are equal to 0 when the activations before and after word *n* are identical). We use the analogous measure over the outputs of the query network, called the explicit subjective update (ESU) to compare to the SU in the developmental simulation reported in the main text:

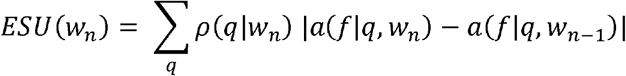

As before we treat *q* as ranging over all of the features of the fillers of all of the probed roles in the sentence. In calculating the ESU or the ESS, the queries associated with the presented sentences are all used, with *ρ*(*q|w_n_*) = 1 for each one.

The simulation results presented in the main text show the same pattern in all cases if the ESS and IS are used rather than the SU and ESU.

### Semantic update driven learning rule

The semantic update driven learning rule introduced in this article for the Sentence Gestalt model is motivated by the idea that later-coming words in a sentence provide information that can be used to teach the network to optimize the probabilistic representation of sentence meaning it derives from words coming earlier in the sentence. We briefly consider how this idea could be applied to generate signals for driving learning in the query network, in a situation where the teaching signal (in the form of a set of queries and corresponding feature values) corresponding to the actual features of an event are available to the model only after the presentation of the last word of the sentence (designated word *N*). In that situation, the goal of learning for the last word can be treated as the goal of minimizing the KL divergence between the outputs of the query network after word *N* and the target values of the features of the event *t*(*f|q,e*). As in the standard learning rule, this reduces to the cross-entropy, which for a single feature is given by

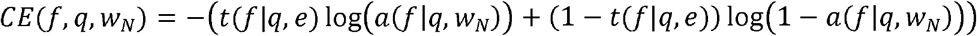

A single {*sentence, event*} pair chosen from the environment would then provide a sample from this distribution. As is the case in the standard training regime, the negative of the gradient with respect to the net input to a given output feature unit in the query network after a given probe is simply *t*(*f*|*q*, *e*) − *a*(*f* | *q*, *W_N_*). This is then the error signal propagated back through the network. To train the network to make better estimates of the feature probabilities from the next to last word in the sentence (word *N*−1), we can use the difference between the activations of the output units after word *N* as the teaching signal for word *N*−1, so for a given feature unit the estimate of the gradient with respect to its net input simply becomes *a*(*f|q,w_N_*) – *a*(*f|q, w*_*N*−1_). Using this approach, as *a*(*f|q,W_N_*) comes to approximate *t*(*f|q,e*) it thereby comes to approximate the correct target for *a*(*f|q, N*−1). This cycle repeats for earlier words, so that as *a*(*f|q, N*−1) comes to approximate *a*(*f|q, N*) and therefore *t*(*f|q, e*) it also comes to approximate the correct teacher for *a*(*f|q, N*−2), etc. This approach is similar to the temporal difference (TD) learning method used in reinforcement learning^91^ in situations where reward becomes available only at the end of an episode, except that here we would be learning the estimates of the probabilities for all of the queries rather than a single estimate of the final reward at the end of an episode. This method is known to be slow and can be unstable, but it could be used in combination with learning based on episodes in which teaching information is available throughout the processing of the sentence, as in the standard learning rule for the SG model.

The semantic update based learning rule we propose extends the idea described above, based on the observation that the pattern of activation over the SG layer of the update network serves as the input pattern that allows the query network to produce estimates of probabilities of alternative possible responses to queries after it has seen some or all of the words in a sentence. Consider for the moment an ideally trained network in which the presentation of each word produces the optimal update to the SG representation based on the environment it had been trained on so far, so that the activations at the output of the query network would correspond exactly to the correct probability estimates. Then using the SG representation after word *n*+1 as the target for training the SG representation after word *n* would allow the network to update its implicit representation based on word *n* to capture changes in the environmental probabilities as these might be conveyed in a sentence. More formally, we propose that changing the weights in the update network to minimize the Implicit Surprise allows the network to make an approximate update to its implicit probabilistic model of sentence meaning, providing a way for the network to learn from linguistic input alone. The negative of the gradient of the Implicit Surprise with respect to the net input to SG unit *i* after word *n* is given by *a_i_*(*w_n_*) – *a_i_*(*w*_*n*−1_). This is therefore the signal that we back propagate through the update network to train the connections during implicit temporal difference learning. As noted in the main text, the sum over the SG units of the absolute value of this quantity also corresponds to the SU, our model’s N400 correlate. The model would not be able to learn language based on this semantic update driven learning rule alone. We assume that language learning proceeds by a mixture of experience with language processed in the context of observed events (as in the standard training regime) and processed in isolation (as with the semantic update driven learning rule), possibly with changing proportions across development. Future modeling work should explore this issue in more detail.

### Simulation Details

#### Environment

The model environment consists of {sentence, event} pairs probabilistically generated online during training according to constraints embodied in a simple generative model (see Fig. 9a). The sentences are single clause sentences such as “At breakfast, the man eats eggs in the kitchen”. They are stripped of articles as well as inflectional markers of tense, aspect, and number, and are presented as a sequence of constituents, each consisting of a content word and possibly one closed class word such as a preposition or passive marker. A single input unit is dedicated to each word in the model’s vocabulary. In the example above, the constituents are “at breakfast”, “man”, “eats”, “eggs”, “in kitchen”, and presentation of the first constituent corresponds to activating the input units for “at” and “breakfast”. The events are characterized as sets of role filler pairs, in this case: agent – man, action – eat, patient – eggs, location – kitchen, situation - breakfast. Each thematic role is represented by a single unit at the probe and output layer. For the filler concepts, we used feature-based semantic representations such that each concept was represented by a number of units (at the probe and output layer) each corresponding to a semantic feature. For instance, the concept “daisy” was represented by five units. The units have labels that allow the reader to keep track of their roles but the model is not affected by the labels themselves, only by the similarity relationships induced by these labels. For example, the semantic features of “daisy” are labeled “can grow”, “has roots”, “has petals”, “yellow”, and “daisy”. The feature-based representations were handcrafted to create graded similarities between concepts roughly corresponding to real world similarities as in other models of semantic representation^92,93^.

**Supplementary Figure 12.**
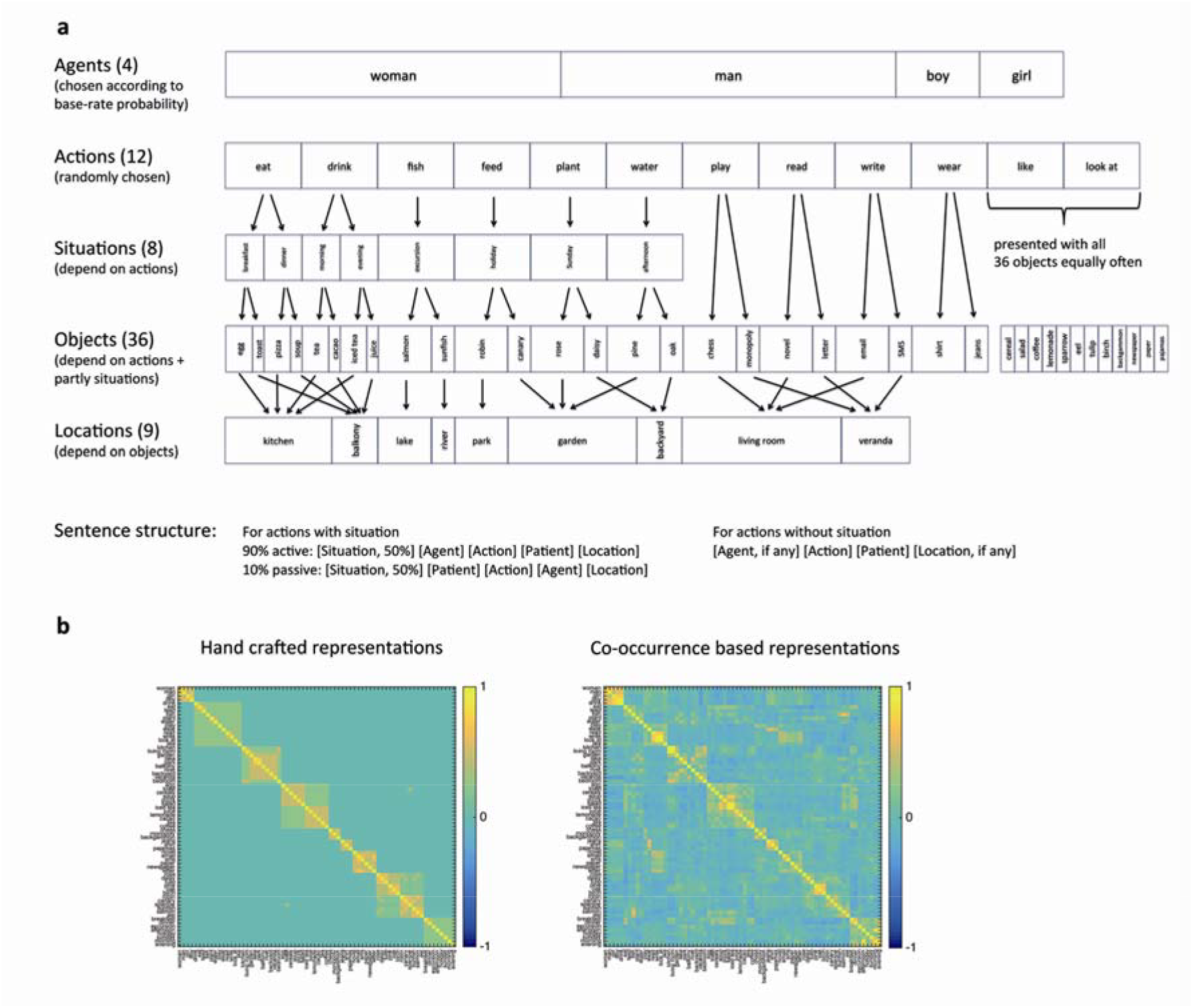
a. The sentence/ event generator used to train the model. Bar width corresponds to relative probability. First, one out of twelve actions is chosen with equal probability. Then, for every action except one (“look at”) an agent is chosen (‘woman “ and “man” each with a probability of .4, “boy” and “girl” with a probability of. 1). Next, a situation is chosen depending on the action. Some actions can occur in two possible situations, some in one, and some without a specified situation. Even if an action occurs in a specific situation, the corresponding word is presented only with a probability of .5 in the sentence while the situation is always part of the event representation. Then, depending on the action (and in the case that an action can occur in two possible situations, depending on the situation) an object/patient is chosen. For each action or situation (except for “like” and “look at” for which all 36 objects are chosen equally often) there is a high probability and a low/probability object (if the agent is “man” or “woman”, the respective high/low probabilities are.7/.3, if the agent is “girl” or “boy”, the probabilities are.6/4). The high and low probability objects occurring in the same specific action context are always from the same semantic category, and for each category, there is a third object which is never presented in that action context and instead only occurs in the unspecific “like” or “look at” contexts (to enable the simulation of categorically related incongruities; these are the twelve rightmost objects in the figure; here bar width is larger than probability to maintain readability). Possible sentence structures are displayed below. b. Similarity matrices of the hand-crafted semantic representations used for the current model (left) and representations based on a principal component analysis on word vectors derived from co-occurrences in large text corpora^94^. The correlation between the matrices is r = .73.

For instance, all living things shared a semantic feature (“can grow”), all plants shared an additional feature (“has roots”), all flowers shared one more feature (“has petals”) and then the daisy had two individuating features (“yellow” and its name “daisy”) so that the daisy and the rose shared three of their five semantic features, the daisy and the pine shared two features, the daisy and the salmon shared only one feature, and the daisy and the email did not share any features (see the Supplementary Table 1 for a complete list of concepts and features). Comparison of a similarity matrix of the concepts based on our hand-crafted semantic representations and representations based on a principal component analysis (PCA) performed on semantic word vectors derived from co-occurrences in large text corpora^94^ showed a reasonable correspondence (*r* = .73; see Fig. 9b), suggesting that the similarities among the hand-crafted conceptual representations roughly matched real world similarities (as far as they can be derived from co-occurrence statistics).

#### Training protocol

The training procedure approximates a situation in which a language learner has observed an event and thus has a complete representation of the event available, and then hears a sentence about it so that learning can be based on a comparison of the current output of the comprehension mechanism and the event. It is important to note that this is not meant to be a principled theoretical assumption but is rather just a practical consequence of the training approach. In general, we do not assume that listeners can only learn when they simultaneously experience a described event, first, because neural networks can generalize^13^ and second, because the SG model can also learn simply from listening or reading based on the new learning rule driven by the semantic update (see section *Semantic update driven learning rule*, above). Also, observed events can be ambiguous and language can provide a particular disambiguating perspective on an event that cannot be gleaned directly from the event itself^95^. The SG model implements a simplification of the situation in the sense that events in the model are always unambiguous and complete. In addition, the training procedure implements the assumption that listeners anticipate the full meaning of each presented sentence as early as possible^96,97^, so that the model can learn to probabilistically preactivate the semantic features of all role fillers involved in the described event based on the statistical regularities in its environment.

Each training trial consists in randomly generating a new {*sentence, event*} pair based on the simple generative model depicted in Fig. 9a, and then going through the following steps: At the beginning of a sentence, all units are set to 0. Then, for each constituent of the sentence, the input unit or units representing the constituent are turned on and activation flows from the input units and – at the same time via recurrent connections - from the SG units to the units in the first hidden layer (Hidden 1), and from these to the units in the SG layer where the previous representation (initially all 0’s) is replaced by a new activation pattern which reflects the influence of the current constituent. The activation pattern at the SG layer is then frozen while the model is probed concerning the event described by the sentence in the query part of the model. Specifically, for each probe question, a unit (representing a thematic role) or units (corresponding to feature-based representations of fillers concepts) at the probe layer are activated and feed into the hidden layer (Hidden 2) which at the same time receives activation from the SG layer. Activation from the SG and the probe layer combine and feed into the output layer where the units representing the complete role-filler pair (i.e., the unit representing the thematic role and the units corresponding to the feature-based representation of the filler concept) should be activated. After each presented constituent, the model is probed once for the filler of each role and once for the role of each filler involved in the described event, and for each response, the model’s activation at the output layer is compared with the correct output. After each response, the gradient of the cross-entropy error measure for each connection weight and bias term in the query network is back-propagated through this part of the network, and the corresponding weights and biases are adjusted accordingly.

At the SG layer, the gradient of the cross-entropy error measure for each connection weight and bias term in the update network is collected for the responses on all the probes for each constituent before being back-propagated through this part of the network and adjusting the corresponding weights and biases. We used a learning rate of 0.00001 and momentum of 0.9 throughout.

#### Simulation of empirical findings

Because the model’s implicit probabilistic representation of meaning and thus also the semantic update at any given point is determined by the statistical regularities in the training set, in the description of the simulations below we try to make clear how the observed effects depend on the training corpus (please refer to Fig. 7a).

#### Main simulations

For the simulations of semantic incongruity, cloze probability, and categorically related semantic incongruity, for each condition one agent (“man”) was presented once with each of the ten specific actions (excluding only “like” and “look at”). The agent was not varied because the conditional probabilities for the later sentence constituents depend very little on the agents (the only effect of the choice of agent is that the manipulation of cloze probability is stronger for “man” and “woman”, namely .7 vs. .3, than for “girl” and “boy”, namely .6 vs. .4; see Fig. 7a). For the simulation of semantic incongruity, the objects were the high probability objects in the congruent condition (e.g., “The man plays chess.”) and unrelated objects in the incongruent condition (e.g., “The man plays *salmon*”). For the simulation of cloze probability, the objects/patients were the high probability objects in the high cloze condition (e.g., “The man plays *chess*.”) and the low probability objects in the low cloze condition (e.g., “The man plays *monopoly*.”). For the simulation of categorically related semantic incongruities, the congruent and incongruent conditions from the semantic incongruity simulation were kept the same and there was an additional condition where the objects were from the same semantic category as the high and low probability objects related to the action (and thus shared semantic features at the output layer, e.g., “The man plays *backgammon*”), but were never presented as patients of that specific action during training (so that their conditional probability to complete the presented sentence beginnings was 0).

Instead, these objects only occurred as patients of the unspecific “like” and “look at” actions (Fig. 7a). For all these simulations, there were 10 items in each condition, and semantic update was computed based on the difference in SG layer activation between the presentation of the action (word *n*−1) and the object (word *n*).

For the simulation of the influence of a word’s position in the sentence, we presented the longest possible sentences, i.e. all sentences that had occurred during training with a situation and a location, including both the version with the high probability ending and the version with the low probability ending of these sentences. There were 12 items in each condition, and semantic update was computed over the course of the sentences, i.e. the difference in SG layer activation between the first and the second word provided the basis for semantic update induced by the second word (the agent), the difference in SG layer activation between the second and the third word provided the basis for semantic update induced by the third word (the action), the difference in SG layer activation between the third and the fourth word provided the basis for semantic update induced by the fourth word (the object/ patient), and the difference in SG layer activation between the fourth and the fifth word provided the basis for semantic update induced by the fifth word (the location). It is interesting to consider the conditional probabilities of the constituents over the course of the sentence: Given a specific situation, the conditional probability of the presented agent (“man”; at the second position in the sentence) is .36 (because the conditional probability of that agent is overall .4, and the probability of the sentence being an active sentence such that the agent occurs in the second position is .9; see Fig. 7a). The conditional probability of the action (at the third position) is 1 because the actions are determined by the situations (see section on reversal anomalies, below, for the rationale behind this predictive relationship between the situation and the action). The conditional probability of the objects (at the fourth position) is either .7 (for high probability objects) or .3 (for low probability objects) so that it is .5 on average, and the conditional probability of the location (at the fifth position) is 1 because the locations are determined by the objects. Thus, the constituents’ conditional probabilities do not gradually decrease across the course of the sentences. The finding that semantic update nonetheless gradually decreased over successive words in these sentences (see *Results*) suggests that the SG layer activation does not perfectly track conditional probabilities. Even if an incoming word can be predicted with a probability of 1.0 so that an ideal observer could in principle have no residual uncertainty, the presentation of the item itself still produces some update, indicating that the model retains a degree of uncertainty, consistent with the ‘noisy channel’ model^98^. In this situation, as we should expect, the SG anticipates the presentation of the item more strongly as additional confirmatory evidence is accumulated, so that later perfectly predictable constituents are more strongly anticipated than earlier ones. In summary, the model’s predictions reflect accumulation of predictive influences, rather than completely perfect instantaneous sensitivity to probabilistic constraints in the corpus.

For the simulation of lexical frequency, the high frequency condition comprised the high probability objects from the ten semantic categories, the two high probability agents (“woman” and “man”) and two high probability locations (“kitchen” and “living room”). The low frequency condition contained the ten low probability objects, the two low probability agents (“girl” and “boy”) and two low probability locations (“balcony” and “veranda”). The high and low frequency locations were matched pairwise in terms of the number and diversity of object patients they are related to (“kitchen” matched with “balcony”, “living room” matched with “veranda”). Before presenting the high versus low frequency words, we presented a blank stimulus to the network (i.e., an input pattern consisting of all 0) to evoke the model’s default activation which reflects the encoding of base-rate probabilities in the model’s connection weights. There were 14 items in each condition, and semantic update was computed based on the difference in SG layer activation between the blank stimulus (word *n*−1) and the high or low frequency word (word *n*).

To simulate semantic priming, for the condition of semantic relatedness, the low and high probability objects of each of the ten semantic object categories were presented subsequently as prime-target pair (e.g., “monopoly chess”). For the unrelated condition, primes and targets from the related pairs were re-assigned such that there was no semantic relationship between prime and target (e.g., “sunfish chess”). For the simulation of associative priming, the condition of associative relatedness consisted of the ten specific actions as primes followed by their high probability patients as targets (e.g., “play chess”). For the unrelated condition, primes and targets were again re-assigned such there was no relationship between prime and target (e.g., “play eggs”). To simulate repetition priming, the high probability object of each semantic category was presented twice (e.g., “chess chess”). For the unrelated condition, instead of the same object, a high probability object from another semantic category was presented as prime. For all priming simulations, there were 10 items in each condition, and semantic update was computed based on the difference in SG layer activation between the prime (word *n*−1) and the target (word *n*).

For the simulation of reversal anomalies, each of the eight situations was presented, followed by the high probability object related to that situation and the action typically performed in that situation (e.g., “At breakfast, the eggs *eat*…”). For the congruent condition, the situations were presented with a possible agent and the action typically performed in that situation (e.g., “At breakfast, the man *eats*…”) and for the incongruent condition, with a possible agent and an unrelated action (e.g., “At breakfast, the man *plants*…”). There were eight items in each condition, and semantic update was computed based on the difference in SG layer activation between the presentation of the second constituent which could be an object or an agent (e.g., “eggs” or “man”; word *n*−1) and the action (word *n*). Please note that in the model environment, the situations predict specific actions with a probability of 1. This prevented the critical words (i.e., the actions) from being much better predictable in the reversal anomaly condition where they are preceded by objects (which in the model environment also predict specific actions with a probability of 1) as compared to the congruent condition where they are preceded by agents (which are not predictive of specific actions at all). Of course, situations do not completely determine actions in the real world. However, the rationale behind the decision to construct the corpus in that way to simulate the reversal anomaly experiment by Kuperberg and colleagues^37^ was that the range of plausibly related actions might be similar for specific situations and specific objects such that actions are not much better predictable in the reversal anomaly than in the congruent condition. A relevant difference between both conditions was that in the reversal anomaly condition the model initially assumed the sentences to be in passive voice, because during training, sentences with the objects presented before the actions had always been in passive voice (see Fig. 7a). Thus, when the critical word was presented without passive marker (i.e., “by”), the model revised its initial assumptions in that regard in the reversal anomaly condition while there was no need for revision in the congruent condition. An oversimplification contained in this implementation is that the model never experiences the eggs in any other role than the patient role, even though eggs can occupy other roles in real language environments such as in the sentence “At breakfast, the eggs ruined the omelet.”. This shortcoming is addressed in the simulation of a third type of reversal anomaly discussed below.

#### Additional simulations of reversal anomalies

We also simulated a second type of reversal anomaly where a relationship between two noun phrases is established prior to encountering the verb^70^ (e.g. “De speer heft de atleten *geworpen*”, lit: “The javelin has the athletes *thrown*”, relative to “De speer werd door de atleten *geworpen*”, lit: “The javelin was by the athletes *thrown*”). For this simulation, we used the same stimuli as for the other reversal anomaly simulation, but with Dutch word order. This makes the sentence structures relevant to examining whether the same mechanism that allows the model to account for the semantic illusion effects reported by Kuperberg et al. would also hold when the verb occurs at the end of the sentence. Thus, the relevant experimental conditions contained sentences such as “The pine was by the man watered.” (i.e., “The pine was watered by the man.” with Dutch word order; congruent condition), “The pine has the man watered.” (i.e. “The pine has watered the man.” with Dutch word order; reversal anomaly condition) and “The pine was by the man drunken.” (i.e., “The pine was drunken by the man.” with Dutch word order; incongruent condition). For this simulation, we trained the model on the same training environment used in the main simulations, with the following modifications. The sentence structures were adjusted such that active sentences were changed from e.g., “The man waters the pine.” to “The man has the pine watered.” and passive sentences were changed from “The pine was watered by the man.” to “The pine was by the man watered.” We added an additional input unit representing “has” and used a single unit for “was by” because both words now always occurred in direct succession (e.g., “ … *was by* the man watered.” instead of “… *was* watered *by* the man.”). Apart from these adjustments, all parameters of the model and training were kept the same. This implementation does not completely correspond to the empirical experiment^70^ in that in our simulation there was no specific relationship between the agent and the action (i.e., the man in the model environment is equally likely to perform all 12 actions and thus was equally likely to water something as he was to drink something, for instance) while in the stimulus material of the empirical experiment there was a specific probabilistic relationship between the agents and the actions (i.e., athletes might be more likely to throw something than to summarize something). However, important for current purposes, this implementation allowed to test whether the way the model accounts for the slight N400 increase in reversal anomalies would be robust to changes in word order, i.e. the presentation of two noun phrases prior to the presentation of the verb. For the simulation, there were eight items in each experimental condition, and semantic update was computed as the difference in SG layer activation between the third constituent (“man”, word *n*-1) and the fourth constituent (the action, word *n*). The results are displayed in Supplementary Figure 3. As expected and consistent with the experimental findings, the SU at the verb in the reversal anomaly sentences is only slightly larger than the SU in the congruent control condition, and the SU for the incongruent verb condition is much larger.

Finally, we simulated a type of reversal anomaly where both noun phrases could be agents in events such as in the example “De vos die op de stroper *joeg*” (lit: The fox that on the poacher *hunted*)^40^ or “De zieken die in de chirurg sneden” (lit: The patient that into the surgeon *cut*). As discussed in the main text, both participants in such sentences can be agents, even in events involving the relevant action, and in events involving both of them and different actions. Although the original experiment used embedded clauses, we used single clause sentences with Dutch word order, e.g., “The fox on the poacher hunted.” and “The patient into the surgeon cut.” For this simulation, we increased the percentage of passive sentences in the model’s environment from 10% to 30% (the implementation of the retrieval-integration model used 50% passive sentences^39^). We do not assume that there are more passive sentences in Dutch than in English, but take the increase of the rate of passive sentences to be a simple approximation to the situation that a major grammatical difference between English and Dutch lies in the number of permissible word orders so that word order is a less reliable cue to meaning in Dutch. Specifically, a study found SV word order to be a valid cue to the agent role in 95/100 of sentences in English but only 35/100 sentences in Dutch^41^. As we assume that the model simultaneously uses all available constraints to map from incoming words to sentence meaning, we assume this variability in terms of word order in Dutch also plays a role in its interpretation of reversal anomaly sentences with two noun phrases that can both be agents. To capture the relevant features, we extended the training environment for this simulation by adding eight analogous scenarios to the main simulation environment (Supplementary Fig. 13). To keep the overall size of the training environment roughly similar to the main simulation, we eliminated six actions with their respective objects and situations from the environment when adding these new scenarios. In keeping with the characteristics of the materials used in the experiment, which were often built around a typical event that quickly comes to mind when particular participants are involved (for example, surgeons typically operate on patients), we constructed each of the new scenarios around a probable event involving a central agent doing a central action to a central patient (e.g., the poacher hunting the fox or the surgeon operating on the patient), but alternative events can occur with lower probability as well. In particular, the central patient can also perform the central action, but not towards the central agent. For instance, the fox can also hunt (though not the poacher) and the patient can also cut into something (though not into the surgeon). Furthermore, there are alternative less specific actions as well (such as approaching, watching, standing or sitting in front of) that can be performed by all sorts of agents (including the central patients) towards all sorts of patients (including the central agents). Thus, the central patients can sometimes also be agents in events involving the central agents, e.g., the fox can watch the poacher and the patient can stand in front of the surgeon.

As noted in the main text, there is considerable variability among the materials used in the empirical experiment^40^ as is apparent from the different examples involving the fox/poacher and patient/ surgeon, and that the ERP is averaged across all these materials. Thus, we designed our scenarios (see Suppl. Fig. 13 for details) based on our examination of the entire set of the experimental materials, instead of trying to capture any particular scenario exactly. Furthermore, we cannot claim to have exactly matched the average probabilities of actions performed by the various participants across the full set of materials used in the actual experiment. The richness and diversity of the experimental materials along a number of relevant dimensions makes it difficult to determine how well we have approximated the factors that influence the construction of a representation of meaning in these sentences. The current scenarios thus provide a proof of concept that the model can capture the empirical data when taking into account the elements of reversal anomalies described above. This proof-of-concept approach in light of the complexity of the issue is somewhat similar to the approach taken in implementing the training environment for the retrieval integration model^39^.

**Suppl. Fig. 13.**
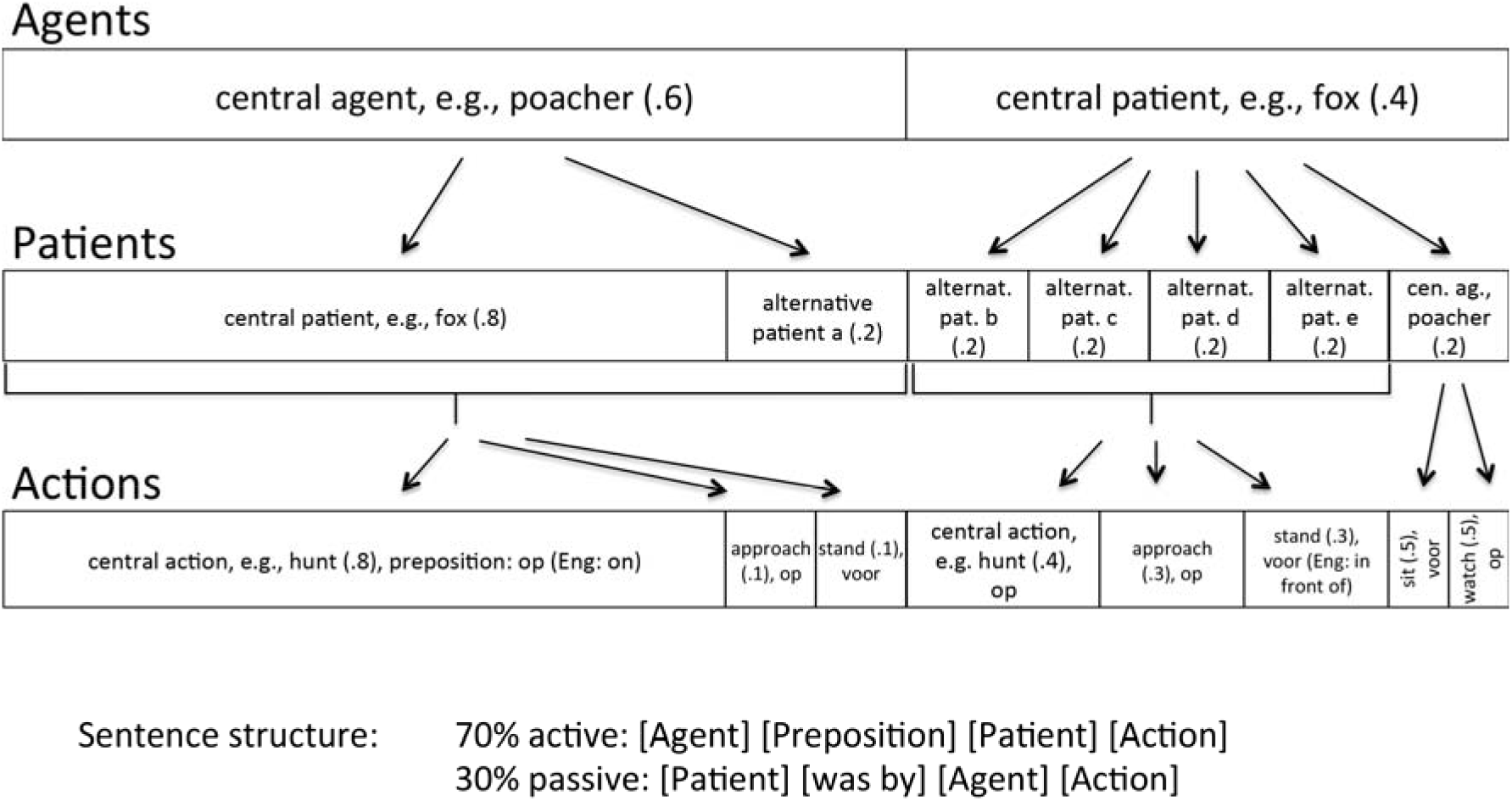
Example scenario from the training environment designed to simulate N400 amplitudes during reversal anomalies with two animate event participants such as “The fox on the poacher hunted.” Eight analogous scenarios were added to the training environment (with Dutch word order) to be able to assess the reliability of the effects across items. English words are used as labels for the scenario participants to help the reader align the design of the scenarios with a natural sentence example; the central agent, patient, and action are the most frequent filler of each role, here labeled ‘poacher’, ‘fox’ and ‘hunt’ respectively. The alternative actions (approach, stand, sit, and watch) were shared across all eight scenarios, with counterbalanced assignment to the different slots in the scheme above (e.g., in another scenario, ‘watch’ and ‘sit’ switch positions with ‘approach’ and ‘stand’). The alternative actions are intended to capture the existence of unspecific actions that can be performed by a wide variety of agents towards a wide variety of patients. To capture the impression that many of the central agents in the experimental sentences were relatively unlikely to be patients in events involving the respective central patient, but could very well be patients in a variety of alternative events, the central agents filled the patient role in different scenarios. Specifically, in two groups of four scenarios each, for each scenario, the central agents from the remaining three scenarios within the group, and additionally one of the central agents from a scenario from the other group, were used as alternative patients b, c, d, and e (see scheme above). Furthermore, all the central agents also occurred as patients in events involving the generic actions ‘like’ and ‘look at’ from the pre-existing part of the environment. Similarly, while the central patients were slightly less likely to occur as agents than the central agents within the scenarios, they could additionally occur as agents in events involving the generic action ‘like’. This is intended to capture the impression that on average the central patients in the experimental sentences were fairly likely to be agents (with some e.g. ‘the fox’ more likely than others, e.g. ‘the patient’). When the central agent was the agent of the event, the alternative patient (alternative patient a in the scheme above) was shared across two scenarios (i.e., there were overall four such patients). Each new concept is represented by four semantic features at the output layer. For current purposes, the crucial point in assigning the semantic features was to avoid any systematic differences between the central agents and the central patients. The central agent and the central patient (‘fox’ and ‘poacher’, above) each have three unique features and one feature labeled ‘animate’ which is shared with the agents in the pre-existing part of the training environment. All actions have three unique features and one feature labeled ‘action’, which is shared with the actions in the pre-existing part of the training environment. The prepositions are not associated with any semantic features. There were two different propositions (these can be thought of as corresponding to the Dutch ‘op’ and ‘voor’, which were used frequently in the experimental materials). Both occurred with half of the central actions and overall occurred equally frequently. The resulting model has 71 units at the input layer and 182 units at the probe and output layer.

Apart from these changes, all parameters of the model and training were kept the same, and again, 10 instances of the model were trained for 800,000 sentences each.

For the congruent condition we presented a sentence describing the most typical event, e.g., “The poacher on the fox hunted”; for the incongruent condition, we presented an unrelated action instead of the most probable action, e.g., “The poacher on the fox planted”; and for the reversal anomaly condition, we presented the most typical action with agent and patient reversed, e.g., “The fox on the poacher hunted”. There were eight items in each condition, and semantic update was computed based on the difference in SG layer activation between the third word, which could be the typical agent or the typical patient (e.g., “poacher” or “fox”; word *n*−1) and the action (word *n*).

#### Supplementary analysis of third reversal anomaly simulation

As noted in the main text, the model exhibited uncertainty in its interpretation of reversal anomaly sentences. Here we describe the details. Consistent with the original SG simulations^14^, the model’s interpretations are sensitive both to event probability constraints and word-order constraints. (As discussed in the main text, we use the phrase ‘event probability constraints’ to refer to the probability distribution of role fillers in events consistent with the words so far encountered, independent of the order of the words. For example, at the occurrence of the second noun in ‘the poacher on the fox’ and ‘the fox on the poacher’, the words so far encountered are the same, and so by this usage, the event probability constraints on the fillers of the Agent, Patient, and Action roles would be the same as well.) In the reversal anomaly sentence, word order and event probabilities conflict. The model’s representation correspondingly reflects uncertainty and ends up in an inconclusive state (see Suppl. Fig. 14a).

**Supplementary Figure 14.**
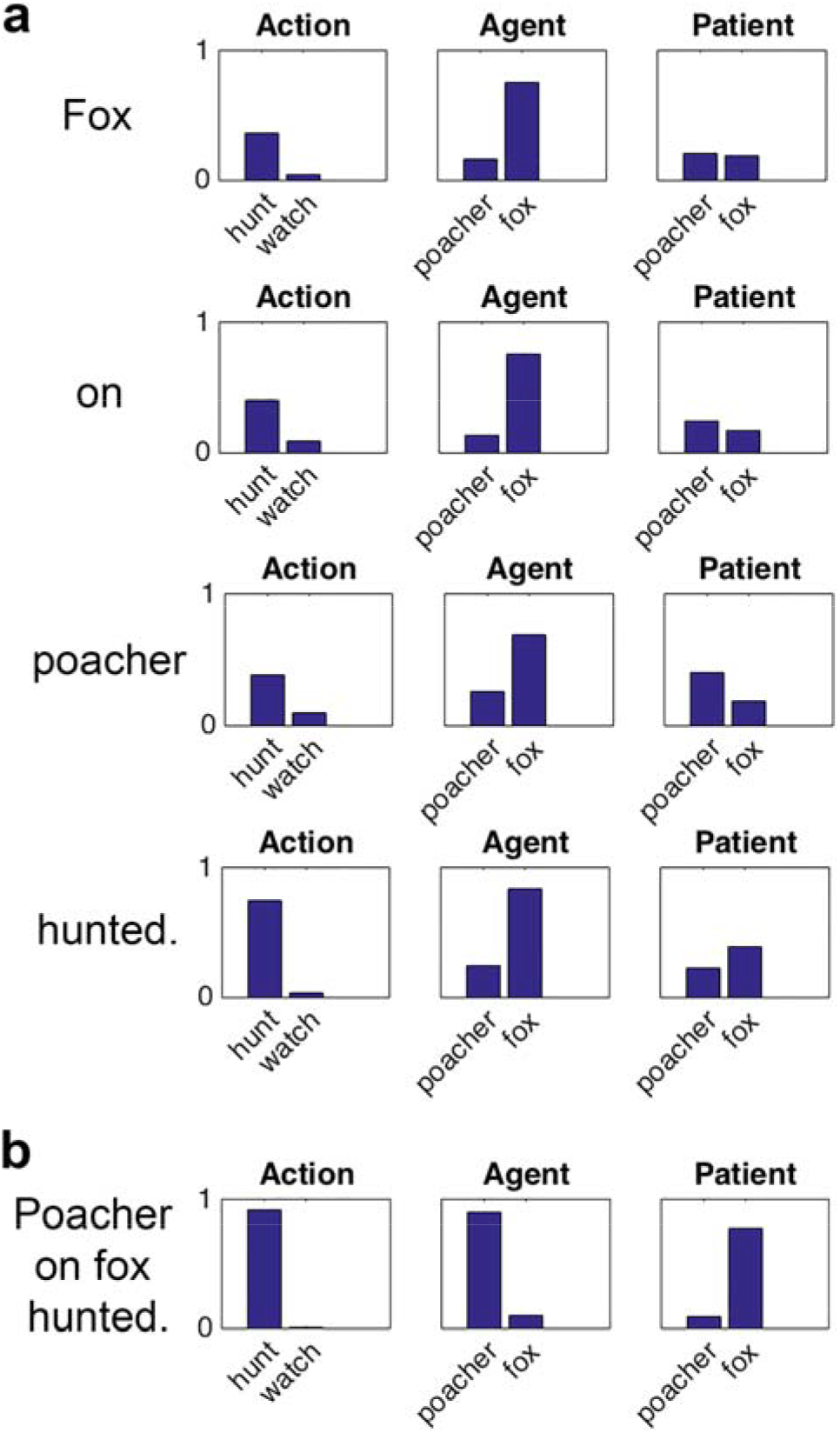
The model’s interpretation of sentences from the simulation of a type of reversal anomaly where both event participants can be agents and can perform the action of interest (see supplementary text for details). The simulation was conducted with a model trained with Dutch word order and the example sentences shown are literal translations from Dutch. For the visualization of the model’s interpretation, English words are used to help the reader map the displayed activations to natural sentence examples. However, it is important to note that the activations are averaged across the respective event participants (‘fox’ representing the central patient, ‘poacher’ representing the central agent, and ‘hunt’ representing the central action) in eight analogous scenarios (see online methods for details). **a.** The model’s average activation of units representing the central agent (‘poacher’ in the example), the central patient (‘fox’ in the example), the central action (‘hunt’ in the example) as well as an alternative action (‘watch’ in the example) when probed for the Action, Agent and Patient role over the course of a reversal anomaly sentence describing an event where the central patient does the central action to the central agent (‘The fox on the poacher hunted.’). **b.** The model’s average activation of the same units after the presentation of the verb in a congruent sentence describing an event where the central agent does the central action to the central patient (‘The poacher on the fox hunted.’). Please note that the model’s representation differs between the congruent and the reversal anomaly sentence. While the interpretation in the congruent sentence is unambiguous and clear, the representation in the reversal anomaly sentence reflects a state of unresolved conflict between different cues, demonstrating the model’s joint sensitivity to event probability and word order constraints.

The model starts by favoring the interpretation of the fox as the agent of the event and is uncertain about the patient. When “the poacher” is presented, the model slightly prefers the poacher as the patient (which makes sense based on syntax, i.e. word order and the preposition “on” without passive marker) but at the same time keeps “hunted” as the most probable action (which makes sense based on event probability) and also still maintains the possibility that the poacher may be the agent and the fox may be the patient (counter to the syntactic cues but consistent with event probability). When the final word “hunted” is presented, the model continues to exhibit cue conflict. There is a shift in the probabilities for the filler of the patient role from a tendency towards the poacher to a slight preference for the fox, which makes sense based on event probabilities (the fox is always the patient in events additionally involving a poacher and hunting). At the same time the model maintains a high probability for the fox being the agent based on syntactic cues (word order and active voice) and also maintains the possibility that the poacher could be either the agent (in line with event probabilities) or the patient (in line with the syntax). When the constraints based on word order and event probabilities agree, as in the control sentences, there is a strong preference for one specific interpretation, with very high activations for the correct role fillers and very low activations for alternative role filler pairs (Suppl. Fig 14b). When the constraints conflict, the model may remain in a state of relative uncertainty and indetermination, in line with the notion that representations during human language comprehension can remain underspecified^57^ (Suppl. Fig. 14a).

We also examined the model’s capacity to assign roles correctly when the reversal anomaly context (e.g., ‘the fox on the poacher’) was followed by a verb that it had experienced in such contexts during training (e.g. ‘watched’; see Supplementary Fig. 13 for details on the training environment). The model performed correctly across all of the scenarios tested, in that the correct filler was most active in each of the Agent, Patient, and Action roles. The model does not strongly pre-activate ‘watch’ upon presentation of ‘the fox on the poacher…’ (which would result in a larger SU upon presentation of ‘hunted’) due to the event probability constraints, which favor hunting in events involving poachers and foxes.

#### Additional simulations

To simulate the developmental trajectory of N400 effects we examined the effect of semantic incongruity on semantic update (as described above) at different points in training, specifically after exposure to 10000, 100000, 200000, 400000, and 800000 sentences. To examine the relation between update at the SG layer and update at the output layer (reflecting latent and explicit estimates of semantic feature probabilities, respectively), at each of the different points in training (see above) we computed the update of activation at the output layer (summed over all role filler pairs) analogously to the activation update at the SG layer.

To simulate semantic priming effects on N400 amplitudes during near-chance lexical decision performance in a second language, we examined the model early in training when it had been presented with just 10000 sentences. As illustrated in Figure 5a, at this point the model fails to understand words and sentences, i.e. to activate the corresponding units at the output layer. The only knowledge that is apparent in the model’s performance at the output layer concerns the possible filler concepts for the agent role and their relative frequency, as well as a beginning tendency to activate the correct agent slightly more than the others. Given the high base-rate frequencies of the possible agents, it does not seem surprising that the model learns this aspect of its environment first. At this stage in training, we simulated semantic priming as described above. In addition, even though this has not been done in the empirical study, we also simulated associative priming and influences of semantic incongruity in sentences (as described above).

For the simulation of the interaction between semantic incongruity and repetition, all sentences from the simulation of semantic incongruity (see above) were presented twice, in two successive blocks (i.e., running through the first presentation of all the sentences before running through the second presentation) with connection weights being adapted during the first round of presentations (learning rate = .01). Sentences were presented in a different random order for each model with the restrictions that the presentation order was the same in the first and the second block, and that the incongruent and congruent version of each sentence directly followed each other. The order of conditions, i.e. whether the incongruent or the congruent version of each sentence was presented first was counterbalanced across models and items (i.e., for half of the models, the incongruent version was presented first for half of the items, and for the other half of the models, the incongruent version was presented first for the other half of the items).

It is often assumed that learning is based on prediction error^49–51^. Because the SG layer activation at any given time represents the model’s implicit prediction or probability estimates of the semantic features of all aspects of the event described by a sentence, the change in activation induced by the next incoming word can be seen as the prediction error contained in the previous representation (at least as far as it is revealed by that next word). Thus, in accordance with the widely shared view that prediction errors drive learning, we used a temporal difference (TD) learning approach, assuming that in the absence of observed events, learning is driven by this prediction error concerning the next internal state. Thus, the SG layer activation at the next word serves as the target for the SG layer activation at the current word, so that the error signal becomes the difference in activation between both words, i.e. SG_*n*+1_ – SG_*n*_ (also see section *Semantic update driven learning rule*, above). There were 10 items in each condition, and semantic update was computed during the first and second presentation of each sentence as the difference in SG layer activation between the presentation of the action (word *n*−1) and the object (word *n*).

For the simulation of the influence of violations of word order (phrase structure)^42^, we presented two types of word order changes for each sentence, focusing on sentences starting with a situation, because in these sentences it is easier to keep changes in conditional probabilities of semantic event features relatively low when changing word order. For each sentence, we presented (1) a version where we changed the position of the action and the patient (e.g., “On Sunday, the man *the robin* feeds” compared to “On Sunday, the man *feeds* the robin”; with semantic update computed as the difference in SG layer activation between the presentation of the agent (word *n*−1) and the patient or action, respectively (word *n*)), and (2) a version where we changed the position of the agent and the action (e.g., “On Sunday, *feeds* the man the robin” compared to “On Sunday, *the man* feeds the robin”; with semantic update computed as the difference in SG layer activation between the presentation of the situation (word *n*−1) and the action or agent, respectively (word *n*)). For type (1), changing position of action and patient, the conditional probability of the semantic features associated with the critical word (not at this position in the sentence but in general within the described event) is .7 in the condition with the changed word order and 1.0 in the condition with the normal word order. For type (2), changing position of agent and action, the conditional probability of the semantic features associated with the critical word (again, crucially, not at this position in the sentence but in general within the described event) is 1.0 in the condition with the changed word order and .4 in the condition with the normal word order. Thus, while changes in word order also entail changes in the amount of semantic update of event features, the design of the simulation ensures that influences of word order (syntax) and semantic update can be dissociated. Specifically, the surprise concerning the semantic features of the described event was on average .15 in the condition with the changed word order (.3 for type (1) and 0.0 for type (2)) while it was on average .3 in the condition with the normal word order (0.0 for type (1), and .6 for type (2)). There were 16 items (8 of each type) in each condition (i.e., normal vs. changed word order).

For the simulation of the influence of constraint on unexpected endings, we presented semantically incongruent sentences in the high constraint condition (e.g., “The man eats *the email*.”) and sentences containing an action which was presented with all 36 objects equally often in the low constraint condition (e.g., “The man likes *the email*.”). This captures the crucial point that in both conditions, the presented object is unexpected; in the high constraint condition, another object is highly expected and in the low constraint condition, no specific object is expected. While the situation is slightly different from the empirical experiment^43^ where both continuations were low cloze but plausible, it is the best way to approximate the experimental situation within our training environment. There were 10 items in each condition, and semantic update was computed as the difference in SG layer activation between the presentation of the action (word *n*−1) and the object (word *n*).

### Simple recurrent network model simulations

We trained a classic simple recurrent network^99^ (consisting of an input and output layer with 74 units each, as well as a hidden and context layer with 100 units each) on the same original training corpus as the SG model. Except for the architectural difference, all parameters were kept the same. We then simulated influences of violations of word order (phrase structure), reversal anomalies, and development, as described above for the SG model. The measure for surprisal that we set in relation to N400 amplitudes consists in the summed magnitude of the cross-entropy error induced by the current word (word *n*).

### Statistics

All reported statistical results are based on ten runs of the model each initialized independently (with initial weights randomly varying between +/− .05) and trained with independently-generated training examples as described in section *Simulation Details/ Environment* (N=800000, unless otherwise indicated). In analogy to subject and item analyses in empirical experiments, we performed two types of analyses on each comparison, a model analysis with values averaged over items within each condition and the 10 models treated as random factor, and an item analysis with values averaged over models and the items (N ranging between 8 and 16; please see the previous section for the exact number of items in each simulation experiment) treated as random factor. There is much less noise in the simulations as compared to empirical experiment such that the relatively small sample size (10 runs of the model and 8 to 16 items per condition) should be sufficient. There was no blinding. We used two-sided paired t-tests to analyze differences between conditions; when a simulation experiment involved more than one comparison, significance levels were Bonferroni-corrected within the simulation experiment. Effect size (Cohen’s d) was computed using the function computeCohen_d in MATLAB. To test for the interaction between repetition and congruity, we used a repeated measures analysis of variance (rmANOVA) with factors Repetition and Congruity. To analyze whether our data met the normality assumption for these parametric tests, we tested differences between conditions (for the t-tests) and residuals (for the rmANOVA) for normality with the Shapiro-Wilk test. Using study-wide Bonferroni correction to adjust significance levels for the multiple performed tests, results did not show significant deviations from normality (all *ps* > .15 for the model analyses and > .32 for the item analyses) except for the model analysis of the difference between congruent and incongruent sentences in the third reversal anomaly simulation (Fig. 3; *p* = .0064) and the item analysis of the change in word order (*p* = .066; this might be due to the items in this simulation experiment consisting of two types with slightly different characteristics; see section *Simulation of empirical findings* above). Both effects did not change when using the Wilcoxon signed rank test, which does not depend on the normality assumption. Specifically, the model analysis of the difference between congruent and incongruent sentences remained significant (*p* = .0002; see also caption of Fig. 3), while the item analysis of the change in word order did not reach significance neither in the t-test (see caption of Fig. 4) nor in the Wilcoxon signed rank test (*p* = .10). To further corroborate our results we additionally tested all comparisons with deviations from normality at uncorrected significance levels <.05 using the Wilcoxon signed rank test; all results remained significant. Specifically, in the model analyses deviations from normality at uncorrected significance levels were detected for the semantic incongruity effect (Fig. 2a; *p* = .043), the frequency effect (Fig. 2e; *p* = .044), the difference between categorically related incongruities and congruent completions (Fig. 2d; *p* = .0053), as well as the difference between reversal anomaly and incongruent sentences (Fig. 3; *p* = .034) in the third reversal anomaly simulation. Wilcoxon signed rank tests confirmed significant effects of semantic incongruity (Fig. 2a; *p* = .002), lexical frequency (Fig. 2e; *p* = .037), a significant difference between categorically related incongruities and congruent sentence continuations (Fig. 2d; *p* = .002), as well as significant differences between reversal anomaly and incongruent sentences in the third reversal anomaly simulation (Fig.3; *p* = .002). In the item analyses, deviations from normality at an uncorrected significance level were detected for the difference between low constraint unexpected endings and expected endings (Supplementary Fig. 6; *p* = .03), the difference between incongruent completions and reversal anomalies in the first reversal anomaly simulation in the SG model (Supplementary Fig. 1i; *p* = .012) as well as in the SRN (Supplementary Fig. 7; *p* = .043), and for the difference between changed and normal word order in the SRN (Supplementary Fig. 7; *p* = .011). Again, Wilcoxon signed rank tests confirmed significant differences between low constraint unexpected endings and expected endings (Supplementary Fig. 6; *p* = .002), between the incongruent completions and reversal anomalies in the SG model (Supplementary Fig. 1i; *p* = .0078) and the SRN (Supplementary Fig. 7; *p* = .039), as well as a significant influence of word order in the SRN (*p* = .0004).

Using Levene’s test, we detected violations of the assumption of homogeneity of variances (required for the rmANOVA used to analyze the interaction between repetition and congruity; Fig. 6 and Supplementary Fig. 4) in the item analysis, *F_2_(3)* = 12.05, *p* < .0001, but not in the model analysis, *F_1_* < 1. We nonetheless report the ANOVA results for both analyses because ANOVAs are typically robust to violations of this assumption as long as the groups to be compared are of the same size. However, we additionally corroborated the interaction result from the item ANOVA by performing a two-tailed paired t-test on the repetition effects in the incongruent versus congruent conditions, i.e. we directly tested the hypothesis that the size of the difference in the model’s N400 correlate between the first presentation and the repetition was larger for incongruent than for congruent sentence completions: incongruent (first – repetition) > congruent (first – repetition). Indeed, the size of the repetition effects significantly differed between congruent and incongruent conditions, t_2(9)_ = 10.99, *p* < .0001, and the differences between conditions did not significantly deviate from normality, *p* = .44, thus fulfilling the prerequisites for performing the t-test.

In general, systematic deviations from normality are unlikely for the results by-model (where apparent idiosyncrasies are most probably due to sampling noise), but possible in the by-item data. Thus, while we present data averaged over items in the figures in the main text in accordance with the common practice in ERP research to analyze data averaged over items, for transparency we additionally display the data averaged over models as used for the by-item analyses (see Supplementary Fig. 1 and 3-10).

### Code availability

All computer code used to run the simulations and analyze the results will be made available on github at the time of publication.

## Author contributions

M.R. developed the idea for the project, including the idea of linking the N400 to the updating of SG layer activation in the model. S.S.H. re-implemented the model for the current simulations. M.R. and J.L.M. formulated the training environment. J.L.M. formulated the new learning rule and developed the probabilistic formulation of the model with input from M.R. M.R. adjusted the model implementation, implemented the training environment, formulated and implemented the simulations, trained the networks and conducted the simulations, and performed the analyses with input from J.L.M. J.L.M. and M.R. discussed the results and wrote the manuscript.

## Acknowledgements

This project has received funding from the European Union’s Horizon 2020 research and innovation programme under the Marie Sklodowska-Curie grant agreement No 658999 to Milena Rabovsky. We thank Roger Levy, Stefan Frank, and the members of the PDP lab at Stanford for helpful discussion.

---

**Supplementary Table 1.** Words (i.e. labels of input units) and their semantic representations (i.e., labels of the output units by which the concepts that the words refer to are represented)

| **Words** | **Semantic representations** |
| --- | --- |
| Woman | person, animate, adult, female, woman |
| Man | person, animate, adult, male, man |
| Girl | person, animate, child, female, girl |
| Boy | person, animate, child, male, boy |
| Drink | action, consume, done with liquids, drink |
| Eat | action, consume, done with foods, eat |
| Feed | action, done to animals, done with food, feed |
| Fish | action, done to fishes, done close to water, fish |
| Plant | action, done to plants, done with earth, plant |
| Water | action, done to plants, done with water, water |
| Play | action, done with games, done for fun, play |
| Wear | action, done with clothes, done for warming, wear |
| Read | action, done with letters, perceptual, read |
| Write | action, done with letters, productive, write |
| Look at | action, visual look at |
| Like | action, positive, like |
| Kitchen | location, inside, place to eat, kitchen |
| Living room | location, inside, place for leisure, living room |
| Bedroom | location, inside, place to sleep, bedroom |
| Garden | location, outside, place for leisure, garden |
| Lake | location, outside, place with animals, lake |
| Park | location, outside, place with animals, park |
| Balcony | location, outside, place to step out, balcony |
| River | location, outside, place with water, river |
| Backyard | location, outside, place behind house, backyard |
| Veranda | location, outside, place in front of house, veranda |
| Breakfast | situation, food related, in the morning, breakfast |
| Dinner | situation, food related, in the evening, dinner |
| Excursion | situation, going somewhere, to enjoy, excursion |
| Afternoon | situation, after lunch, day time, afternoon |
| Holiday | situation, special day, no work, holiday |
| Sunday | situation, free time, to relax, Sunday |
| Morning | situation, early, wake up, morning |
| Evening | situation, late, get tired, evening |
| Egg | consumable, food, white, egg |
| Toast | consumable, food, brown, toast |
| Cereals | consumable, food, healthy, cereals |
| Soup | consumable, food, in bowl, soup |
| Pizza | consumable, food, round, pizza |
| Salad | consumable, food, light, salad |
| Iced tea | consumable, drink, from leaves, iced tea |
| Juice | consumable, drink, from fruit, juice |
| Lemonade | consumable, drink, sweet, lemonade |
| Cacao | consumable, drink, with chocolate, cacao |
| Tea | consumable, drink, hot, tea |
| Coffee | consumable, drink, activating, coffee |
| Chess | game, entertaining, strategic, chess |
| Monopoly | game, entertaining, with dice, monopoly |
| Backgammon | game, entertaining, old, backgammon |
| Jeans | garment, to cover body, for legs, jeans |
| Shirt | garment, to cover body, for upper part, shirt |
| Pajamas | garment, to cover body, for night, pajamas |
| Novel | contains language, contains letters, art, novel |
| Email | contains language, contains letters, communication, email |
| SMS | contains language, contains letters, communication, short, SMS |
| Letter | contains language, contains letters, communication, on paper, letter |
| Paper | contains language, contains letters, scientific, paper |
| Newspaper | contains language, contains letters, information, newspaper |
| Rose | can grow, has roots, has petals, red, rose |
| Daisy | can grow, has roots, has petals, yellow, daisy |
| Tulip | can grow, has roots, has petals, colorful, tulip |
| Pine | can grow, has roots, has bark, green, pine |
| Oak | can grow, has roots, has bark, tall, oak |
| Birch | can grow, has roots, has bark, white bark, birch |
| Robin | can grow, can move, can fly, red, robin |
| Canary | can grow, can move, can fly, yellow, canary |
| Sparrow | can grow, can move, can fly, brown, sparrow |
| Sunfish | can grow, can move, can swim, yellow, sunfish |
| Salmon | can grow, can move, can swim, red, salmon |
| Eel | can grow, can move, can swim, long, eel |
| By | passive voice (activated together with the deep subject, e.g., 'by the man') |
| Was | passive voice (activated together with the verb, e.g., 'was played') |
| During/at | no output units (activated together with situation words, e.g., 'at breakfast') |
| In | no output units (activated together with location words, e.g., 'in the park') |

**Supplementary Figure 1.**
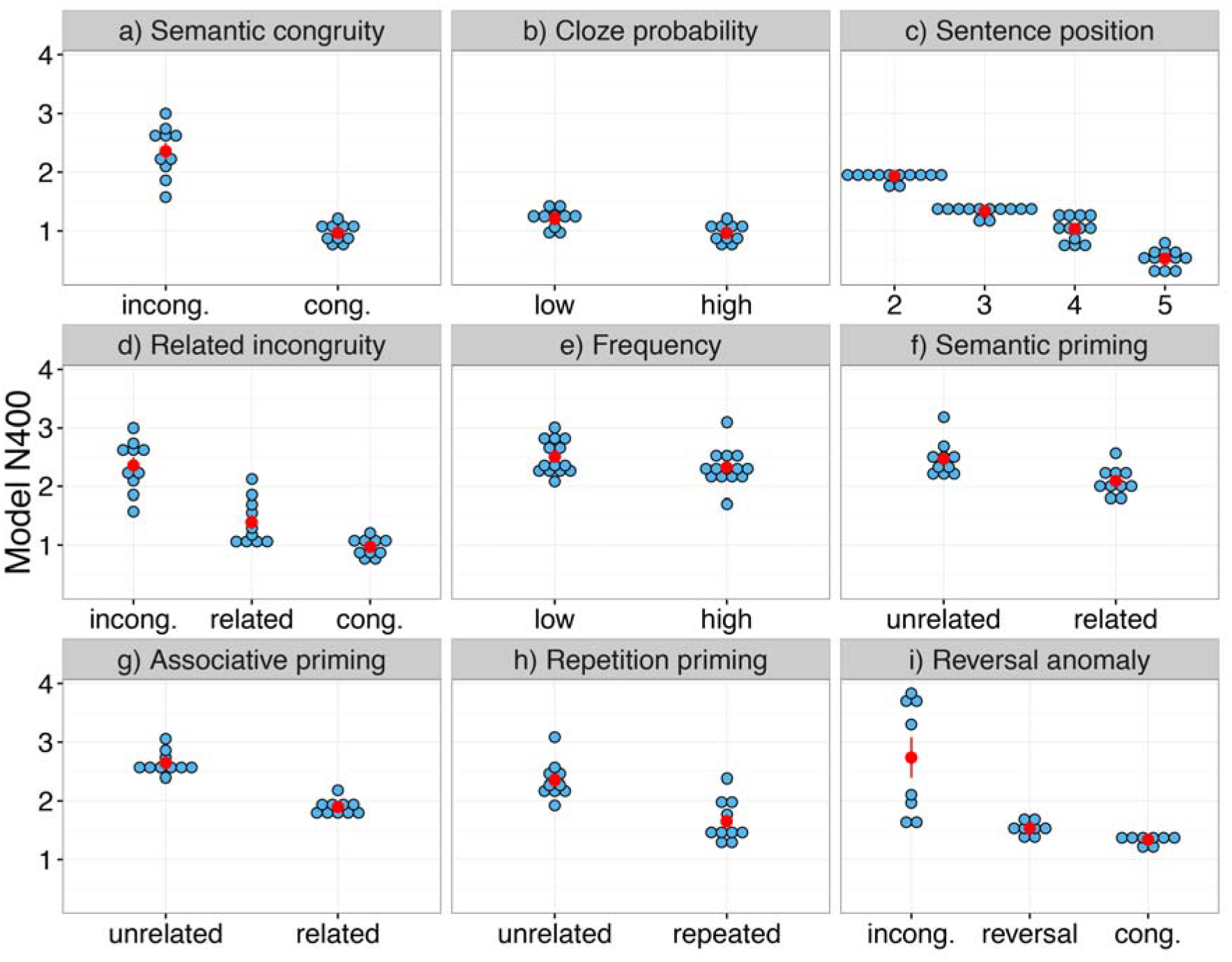
Simulation results for the basic effects (by item). Displayed is the model’s N400 correlate, i.e. the update of the Sentence Gestalt layer activation – the model’s probabilistic representation of sentence meaning - induced by the new incoming word. Cong., congruent; incong., incongruent. See text for details of each simulation. Here, each blue dot represents the results for one item, averaged across 10 independent runs of the model; the red dots represent the means for each condition, and red error bars represent +/− SEM (sometimes invisible because bars may not exceed the area of the red dot). Statistical results are reported in the caption of Fig. 2 in the main text.

**Supplementary Figure 2.**
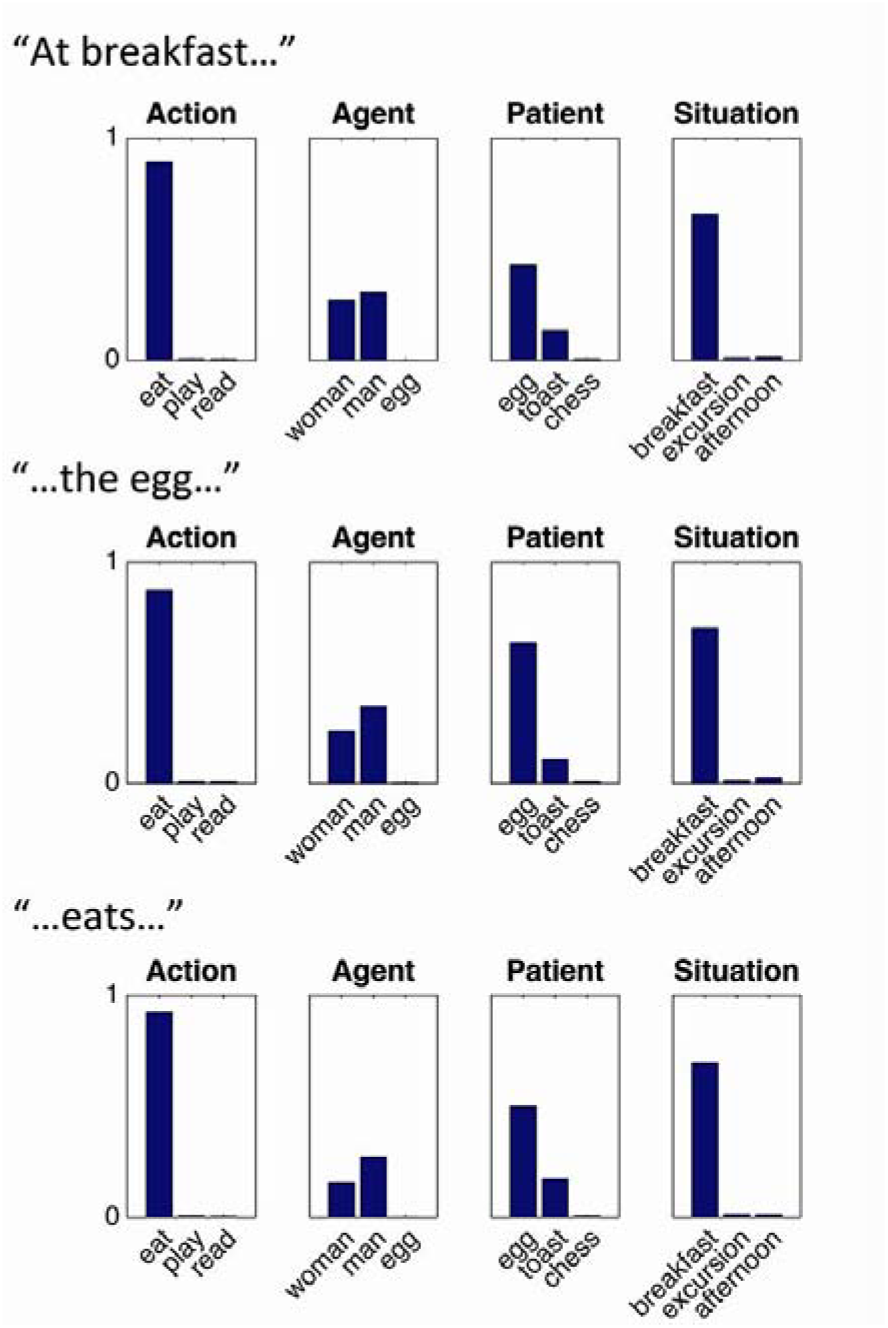
Processing reversal anomalies. Activation of selected output units while the model processes a sentence from the semantic illusion simulation: “At breakfast, the egg eats …”. Note that the model continues to represent the egg as the patient (not the agent) of eating, even after the word “eat” has been presented.

**Supplementary Figure 3.**
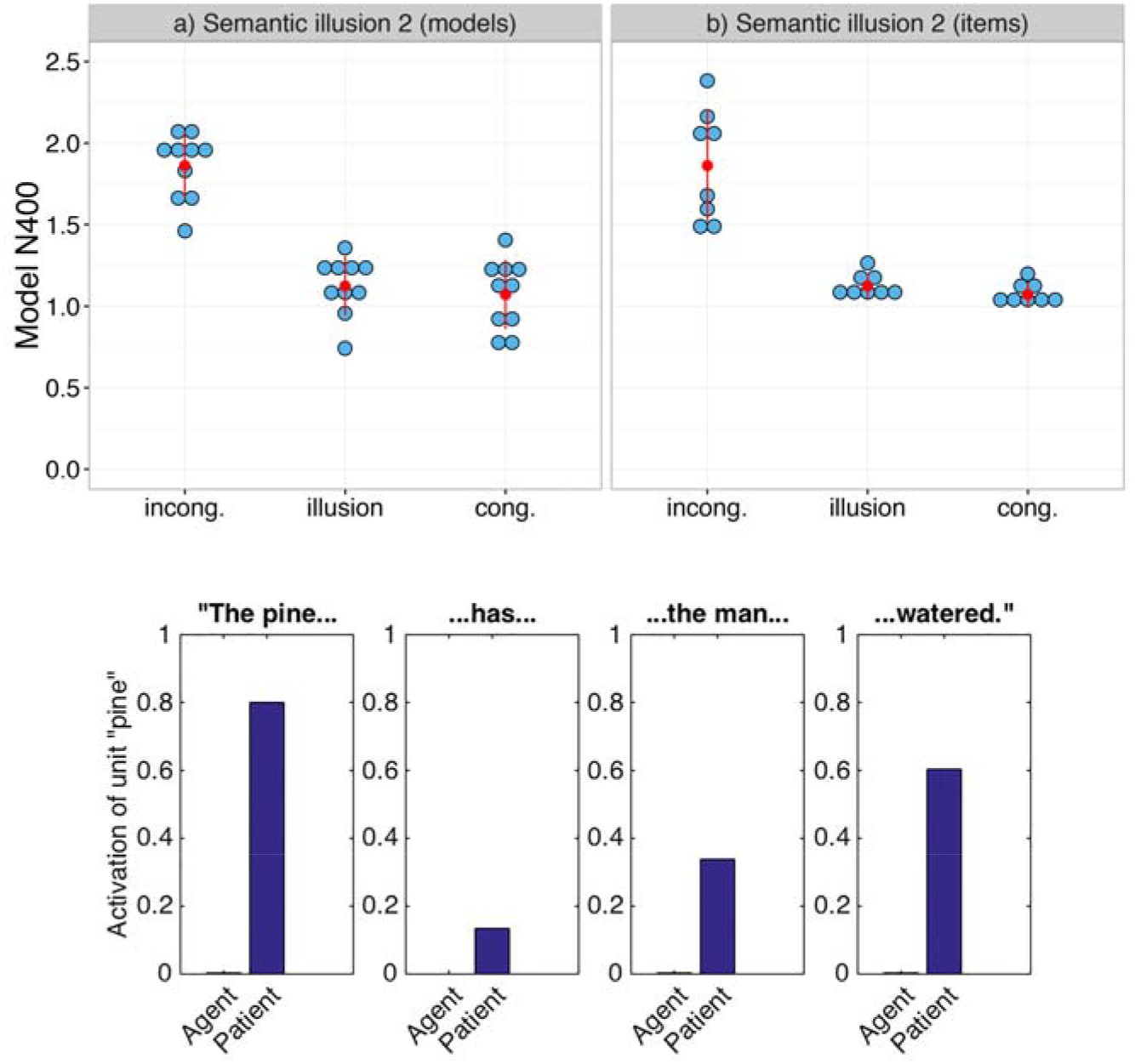
Simulation results for a second type of reversal anomaly where the relationship between two noun phrases is established prior to encountering the verb (see text for more details)^70^; the simulation was conducted with a model trained with Dutch word order. Cong., congruent; incong., incongruent. Top left. Each blue dot represents the results for one independent run of the model, averaged across items per condition. Top right. Each blue dot represents the results for one item, averaged across 10 independent runs of the model. The red dots represent the means for each condition, and red error bars represent +/− SEM. Results are similar as for the other reversal anomaly simulation: t_1(9)_ = 1.69, p = .38, t_2(7)_ = 12.67, p < .0001, for the comparison between congruent condition and reversal anomaly; t_1(9)_ = 13.31, p < .0001, t_2(7)_ = 6.76, p < .001, for the comparison between reversal anomaly and incongruent condition, and t_1(9)_ =12.18, p < .0001, t_2(7)_ = 7.36, p < .001, for the comparison between congruent and incongruent condition. Bottom. Activation of the unit “pine” in response to the Agent and Patient probe while the model processes a sentence from this reversal anomaly simulation, literally “The pine has the man watered.” (i.e., “The pine has watered the man.” with Dutch word order). As for with the “eggs” type anomaly sentences, the model represents the pine as the patient instead of the agent of the event throughout the sentence.

**Supplementary Figure 4.**
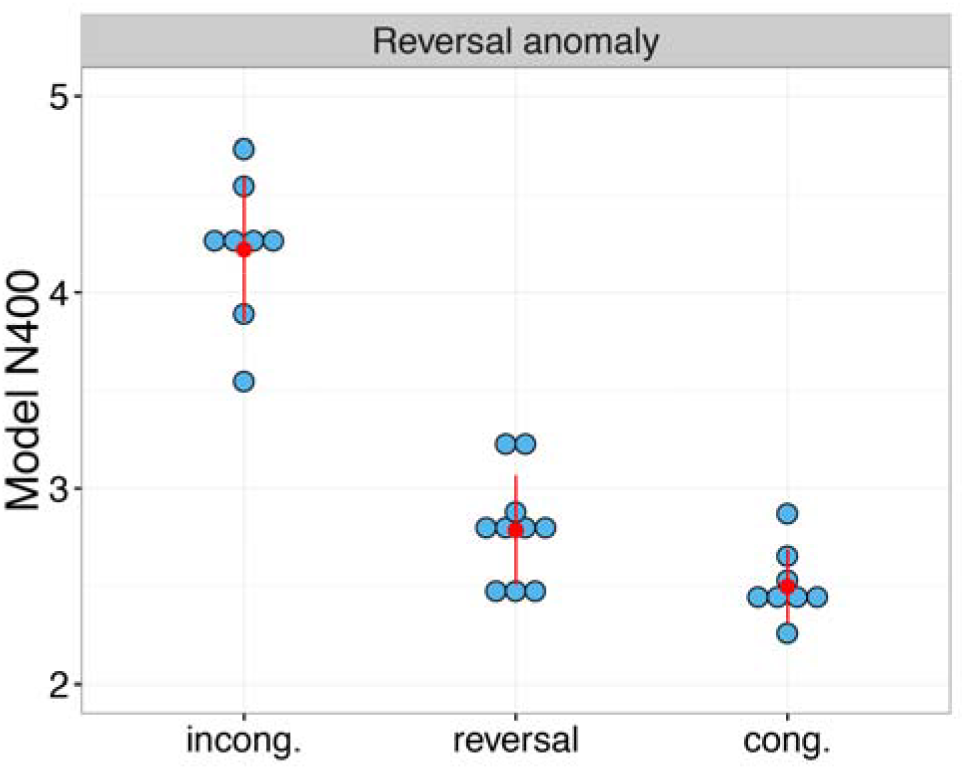
Simulation results of a type of semantic illusion where both event participants can be agents and can perform the action of interest (by item). Here, each blue dot represents the results for one item, averaged across 10 independent runs of the model; the red dots represent the means for each condition, and red error bars represent +/− SEM. Statistical results are reported in the caption of Fig. 3 in the main text:

**Supplementary Figure 5.**
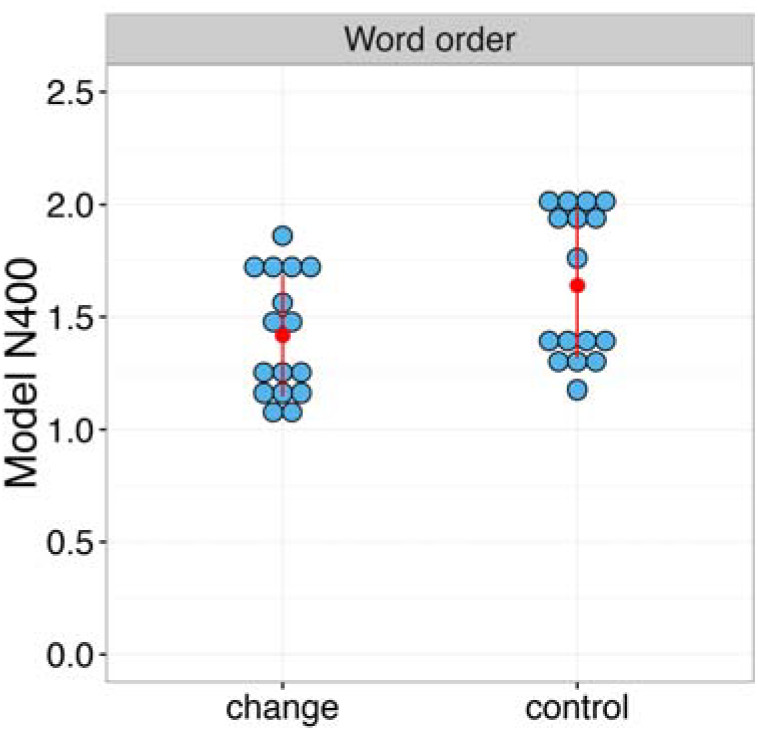
Simulation of the influence of a change in word order (by item). Change, changed word order; control, normal word order. Each blue dot represents the results for one item, averaged across 10 runs of the model; red dots represent means for each condition, and red error bars represent +/− SEM. Statistical results are reported in the caption of Fig. 4.

**Supplementary Figure 6.**
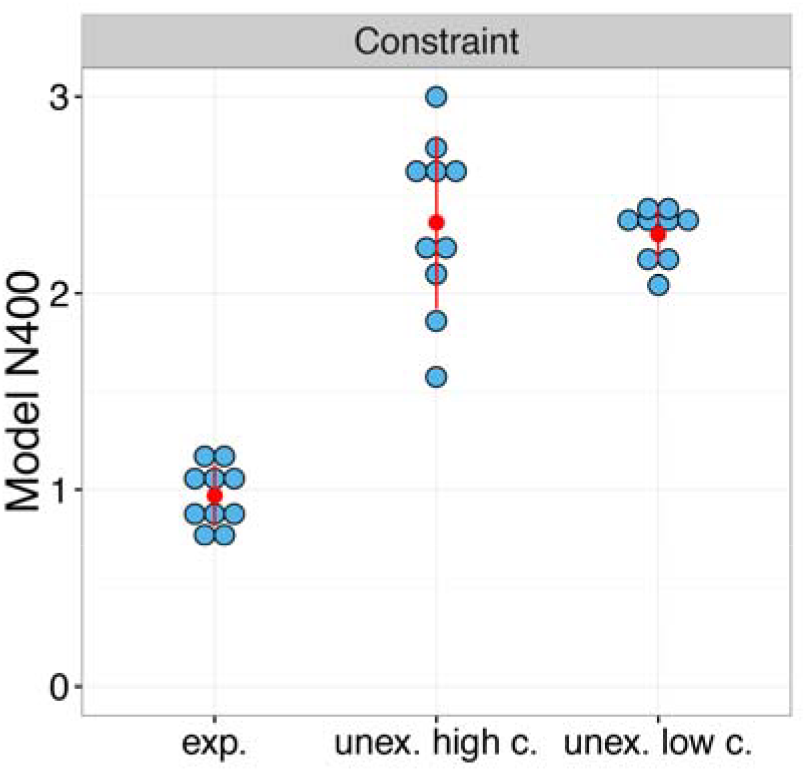
Simulation of the influence of constraint for unexpected endings (by item). Exp., expected; unex., unexpected; c., constraint. Each blue dot represents the results for one item, averaged across 10 runs of the model; red dots represent means for each condition, and red error bars represent +/− SEM. Statistical results are reported in the caption of Fig. 6.

**Supplementary Figure 7.**
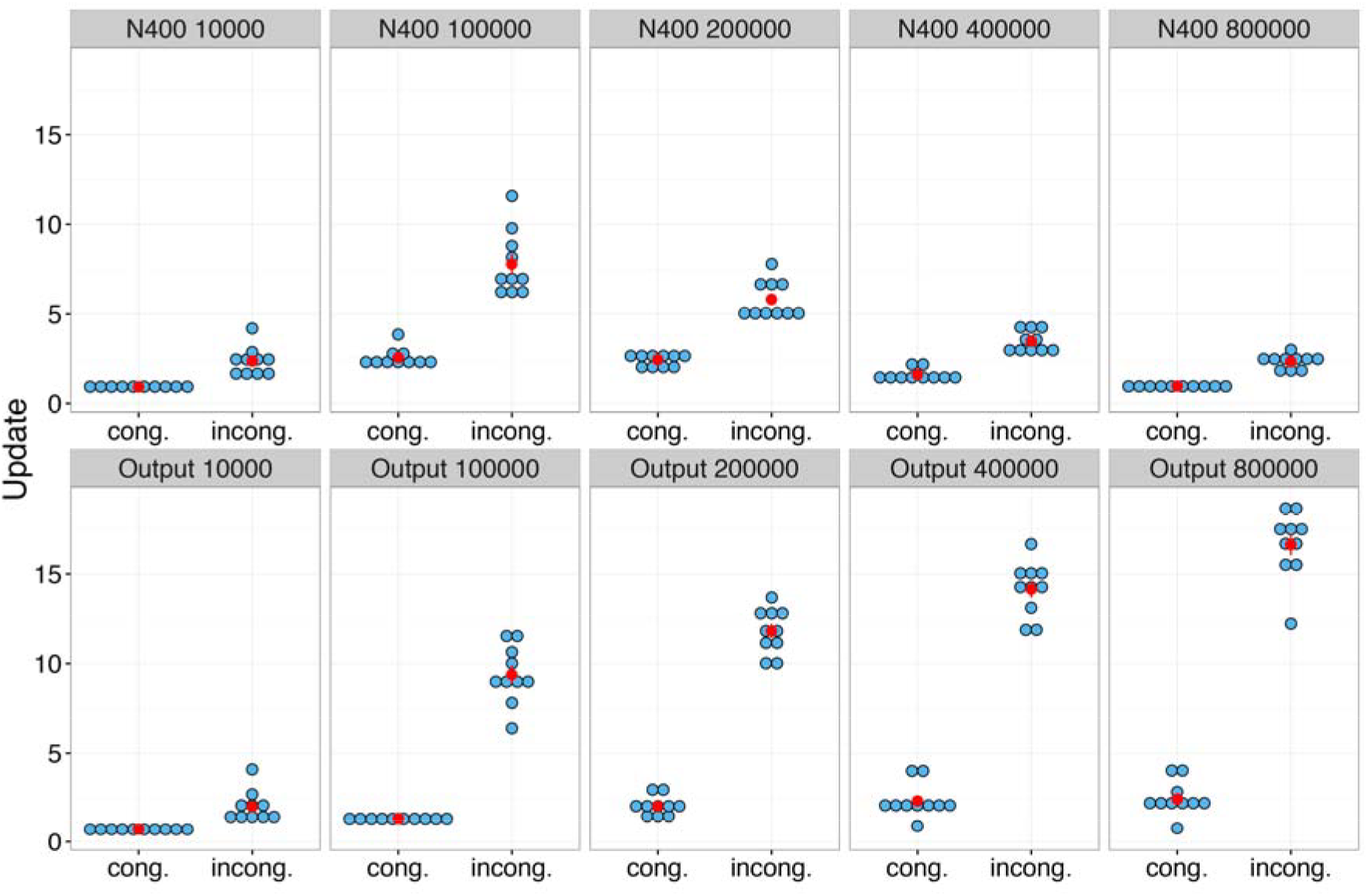
Development across training (by item). Semantic incongruity effects as a function of the number of sentences the model has been exposed to. Top. Semantic update at the model’s hidden Sentence Gestalt layer shows at first an increase and later a decrease with additional training, in line with the developmental trajectory of the N400. Each blue dot represents the results for one item, averaged across 10 independent runs of the model; the red dots represent the means for each condition, and red error bars represent +/− SEM. Statistical results are reported in the caption of Fig. 5 in the main text. Bottom. Activation update at the output layer steadily increases with additional training, reflecting closer and closer approximation to the true conditional probability distributions embodied in the training corpus.

**Supplementary Figure 8.**
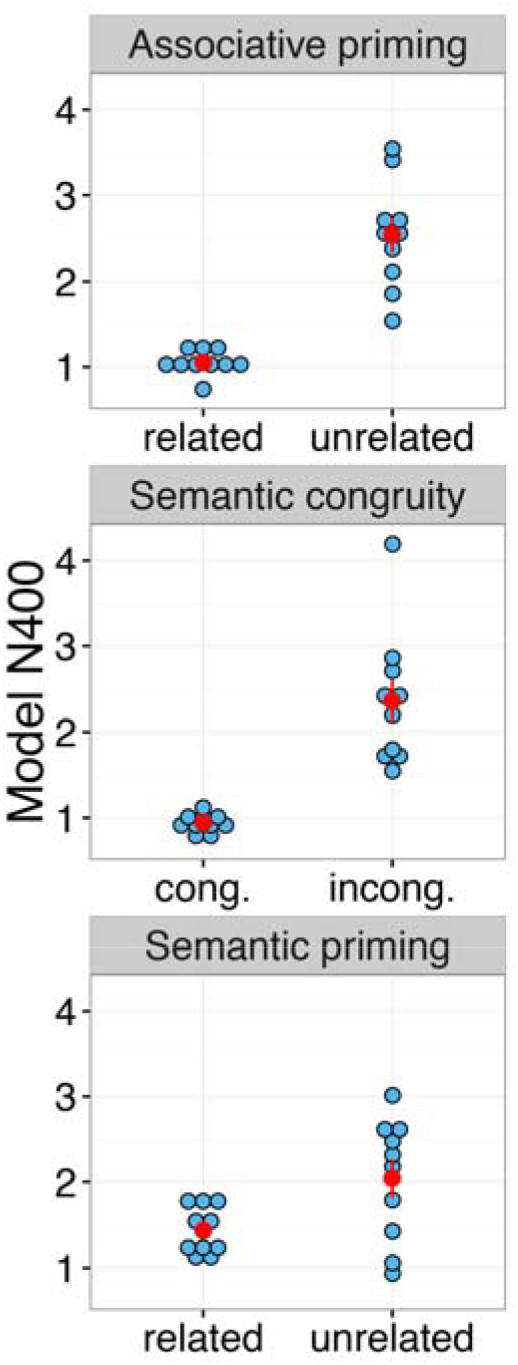
Comprehension performance and semantic update effects at a very early stage in training (by item). Cong., congruent; incong., incongruent. Even at a low level of performance (see Fig. 5a in the main text for illustration), there are robust effects of associative priming(top), semantic congruity in sentences (middle), and semantic priming (bottom). Here, each blue dot represents the results for one item, averaged across ten independent runs of the model; the red dots represent the means for each condition, and red error bars represent +/− SEM. Statistical results are reported in the caption of Fig. 6 in the main text.

**Supplementary Figure 9.**
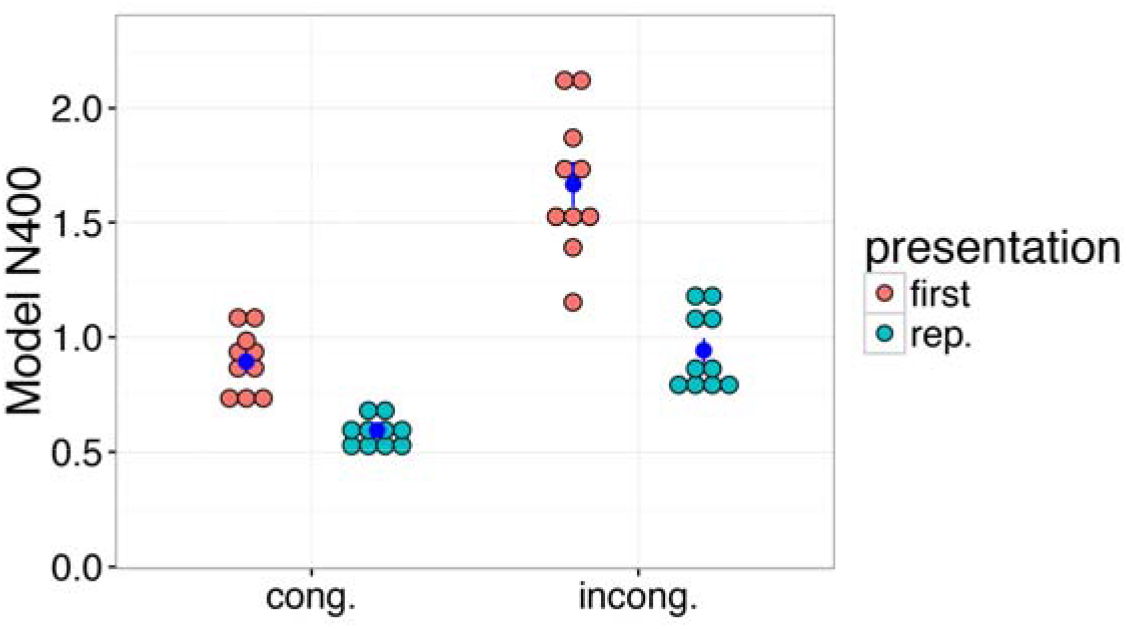
Simulation of the interaction between delayed repetition and semantic incongruity (by item). Each red or green dot represents the results for one item, averaged across 10 runs of the model; blue dots represent means for each condition, and blue error bars represent +/− SEM. Statistical results are reported in the caption of Fig. 7.

**Supplementary Figure 10.**
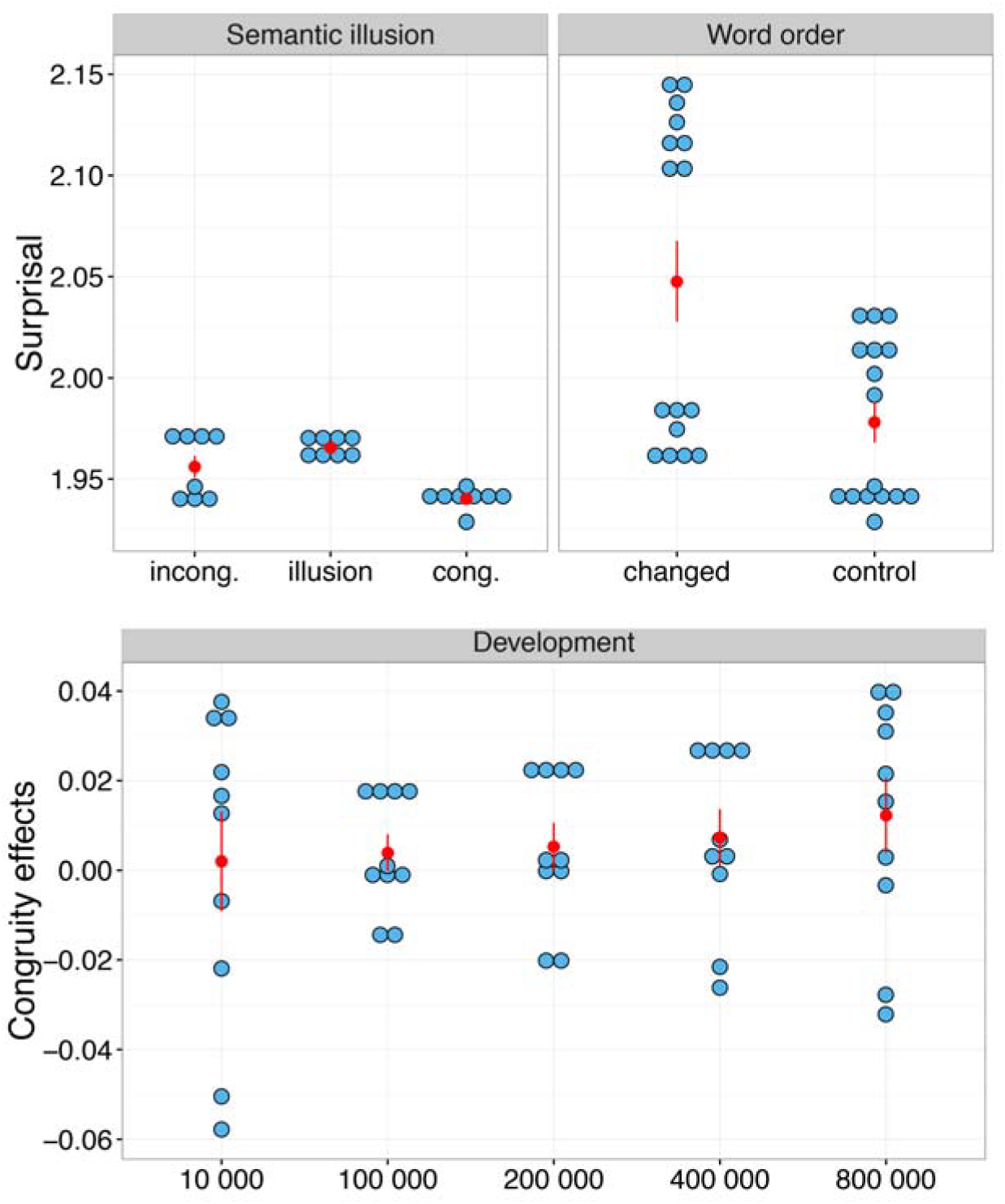
Simulation results from a simple recurrent network model (SRN) trained to predict the next word based on the preceding context. Each blue dot represents the results for one item, averaged across 10 runs of the model; red dots represent means for each condition, and red error bars represent +/− SEM. Statistical results are reported in the caption of Fig. 8. Top left, semantic illusion. Top right, word order. Bottom, congruity effect on surprisal as a function of the number of sentences the model has been exposed to.

**Supplementary Figure 11.**
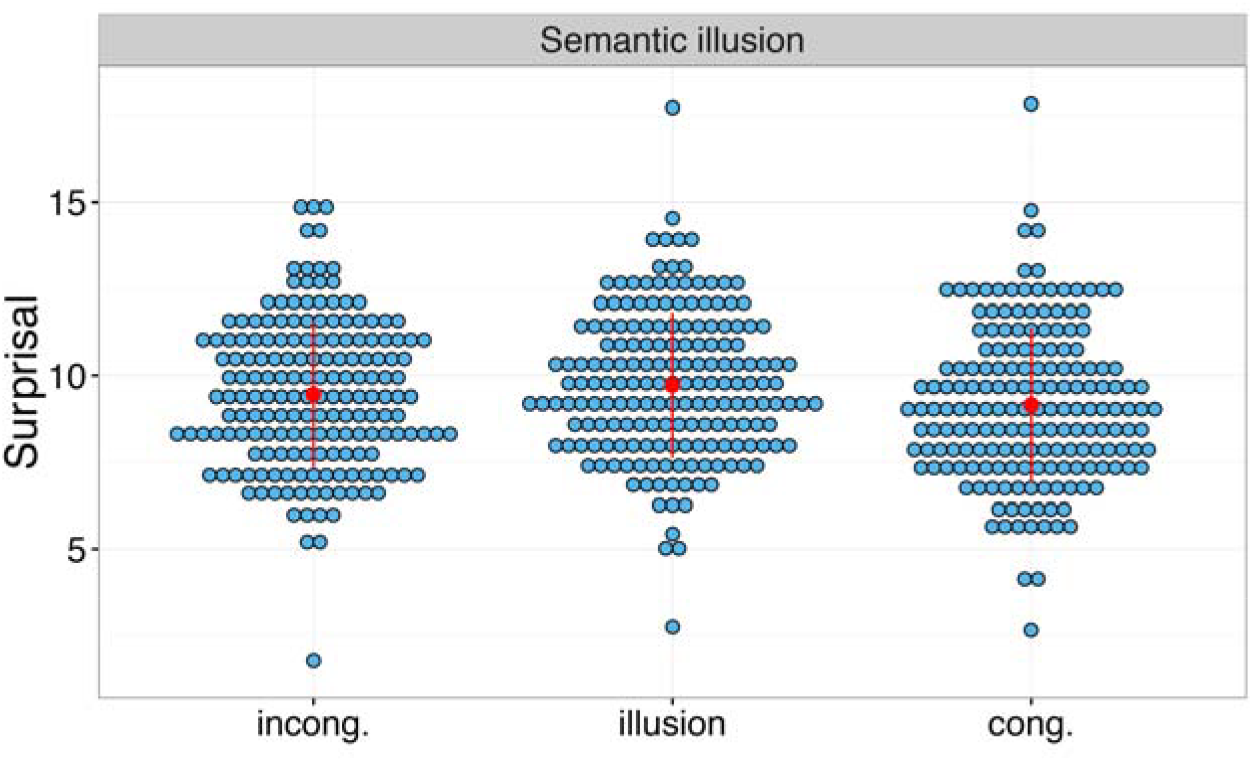
Simulation results from a simple recurrent network (SRN) implementation by T. Mikolov^100^ trained by S. Frank on 23M sentences from a web corpus. Incong., incongruent; cong., congruent. The simulation experiment consisted in the presentation of materials from the semantic illusion experiment by Kuperberg and colleagues which we requested from the authors (there are a few slight differences in the materials due to an issue with retrieving the original stimuli, but the materials largely overlap and resulted in the same pattern of results; G. Kuperberg, personal communication). Each blue dot represents the results for one item, averaged over three runs of the model; the red dots represent the means for each condition, and red error bars represent +/− SEM. Results resemble those from the SRN that we trained on the same corpus as the SG model (Fig. 9 and Supplementary Fig. S10) in that word surprisal was large in the semantic illusion condition, numerically even larger than in the incongruent condition. There were 3 runs of the model and 180 items in each condition (1 less in the incongruent condition because the model did not know one of the words in this condition, “curtseys”) so that we report statistical results from the item analyses: t_2(179)_ = 11.76, *p* < .0001, for the comparison between congruent condition and semantic illusion; t_2(178)_ = 1.29, *p* = .59, for the comparison between semantic illusion and incongruent condition, and t_2(178)_ =1.45, *p*= .45, for the comparison between congruent and incongruent condition. We thank Stefan Frank for performing the simulation and sharing the results with us!

## Footnotes

1 In our model’s N400 correlate, the effect of congru¡ty is much larger than the effect of cloze probability. This result seems to contrast with studies directly comparing low cloze plausible and low cloze implausible continuations and reporting that the N400 is mainly influenced by cloze probability and much less by plausibility^101^. However, note that in the model’s environment, the probability in the low cloze condition was comparatively high (.3), considerably higher than the near-to-zero cloze probability in the mentioned studies^101^, and that low cloze and high cloze continuations in the model environment share semantic features, contributing to the comparatively small effect of cloze probability.

2 We use the phrase ‘event probability constraints’ to refer to the probability distribution of role fillers in events consistent with the words so far encountered, independent of the order of the words. For example, at the occurrence of the second noun in ‘the poacher on the fox’ and ‘the fox on the poacher’, the words so far encountered are the same, and so by this usage, the event probability constraints on the fillers of the Agent, Patient, and Action roles would be the same as well.

3 Specifically, in instances where participants understand the dog to be the agent and the man to be the patient, N400 amplitudes on *man* should be small. For interpretations in which the understanding is in line with syntactic conventions, the situation is more complicated. Depending on participant’s prior experience and details of the relevant word order and event probability information, there might be instances where the syntactically-specified assignment was understood immediately and instances where participants temporarily process the sentence in line with event probability constraints or experience uncertainty that gets resolved later in the process. Thus, in these cases, one might expect to either see a large N400 (reflecting immediate interpretation in line with syntactic conventions and thus considerable semantic update) ora small N400 (reflecting a temporary event probability based interpretation or uncertainty) accompanied by an indication of an enhanced controlled update process later in the process (which might be reflected in increased P600 amplitudes).

## References

1. Kutas, M. & Hillyard, S. A. Reading senseless sentences: Brain potentials reflect semantic incongruity. Science (80-.). 207, 203–205 (1980).

2. Kutas, M. & Federmeier, K. D. Thirty years and counting: finding meaning in the N400 component of the event-related brain potential (ERP). Annu. Rev. Psychol. 62, 621–647 (2011).

3. Lau, E. F., Phillips, C. & Poeppel, D. A cortical network for semantics: (de)constructing the N400. Nat. Rev. Neurosci. 9, 920–933 (2008).

4. Debruille, J. B. The N400 potential could index a semantic inhibition. Brain Res. Rev. 56, 472–477 (2007).

5. Federmeier, K. D. & Laszlo, S. Chapter 1 Time for Meaning. Electrophysiology Provides Insights into the Dynamics of Representation and Processing in Semantic Memory. Psychology of Learning and Motivation - Advances in Research and Theory 51, (2009).

6. Baggio, G. & Hagoort, P. The balance between memory and unification in semantics: A dynamic account of the N400. Lang. Cogn. Process. 26, 1338–1367 (2011).

7. Brown, C. & Hagoort, P. The processing nature of the N400: Evidence from masked priming. J. Cogn. Neurosci. 5, 34–44 (1993).

8. Kuperberg, G. R. Separate streams or probabilistic inference? What the N400 can tell us about the comprehension of events. Lang. Cogn. Neurosci. 3798, (2015).

9. Chomsky, N. Syntactic structures. (Mouton, 1957).

10. Fodor, J. Modularity of Mind. (MIT Press, 1981).

11. Fodor, J. & Pylyshyn, Z. W. Connectionism and cognitive architecture: A critical analysis. Cognition 28, 3–71 (1988).

12. Jackendoff, R. Foundations of Language: Brain, Meaning, Grammar, Evolution. (Oxford University Press, 2002).

13. McClelland, J. L., St. John, M. & Taraban, R. Sentence comprehension: A parallel distributed processing approach. Lang. Cogn. Process. 4, 287–336 (1989).

14. St. John, M. F. & McClelland, J. L. Learning and applying contextual constraints in sentence comprehension. Artif. Intell. 46, 217–257 (1990).

15. Rohde, D. L. T. A Connectionist Model of Sentence Comprehension and Production. (Carnegie Mellon University, 2002).

16. Laszlo, S. & Plaut, D. C. A neurally plausible Parallel Distributed Processing model of Event-Related Potential word reading data. Brain Lang. 120, 271–281 (2012).

17. Laszlo, S. & Armstrong, B. C. PSPs and ERPs: Applying the dynamics of post-synaptic potentials to individual units in simulation of temporally extended Event-Related Potential reading data. Brain Lang. 132, 22–27 (2014).

18. Cheyette, S. J. & Plaut, D. C. Modeling the N400 ERP component as transient semantic over-activation within a neural network model of word comprehension. Cognition 162, 153–166 (2017).

19. Itti, L. & Baldi, P. Bayesian Surprise Attracts Human Attention. 1–8 (2006). doi:10.1016/j.visres.2008.09.007

20. Bransford, J. D. & Johnson, M. K. Contextual Prerequisites for U nderstanding: Some Investigations of Comprehension and Recall 1. J. Verbal Learning Verbal Behav. 11, 717–726 (1972).

21. Griffiths, T. L., Steyvers, M. & Tenenbaum, J. B. Topics in Semantic Representation. 114, 211–244 (2007).

22. Andrews, M., Vigliocco, G. & Vinson, D. Integrating experiential and distributional data to learn semantic representations. Psychol. Rev. 116, 463–498 (2009).

23. Wu, Y. et al. Google’s neural machine translation system: Bridging the gap between human and machine translation. arXiv:1609.08144 (2016).

24. Seidenberg, M. S. & McClelland, J. L. A Distributed, Developmental Model of Word Recognition and Naming. Psychol. Rev. 96, 523–568 (1989).

25. McClelland, J. L. in The Handbook of Language Emergence (eds. MacWhinney, B. & O’Grady, W.) 54–80 (John Wiley & Sons, 2015).

26. Barsalou, L. W. Grounded Cognition. (2008). doi:10.1146/annurev.psych.59.103006.093639

27. Pulvermüller, F. Words in the brain’s language. Behav. Brain Sci. 253–336 (1999).

28. Kutas, M. & Hillyard, S. A. Brain potentials during reading reflect word expectancy and semantic association. Nature 307, 101–103 (1984).

29. Van Petten, C. & Kutas, M. Influences of semantic and syntactic context on open- and closed-class words. Mem. Cogn. 19, 95–112 (1991).

30. Levy, R. Expectation-based syntactic comprehension. Cognition 106, 1126–1177 (2008).

31. Frank, S. L., Galli, G. & Vigliocco, G. The ERP response to the amount of information conveyed by words in sentences. Brain Lang. 140, 1–25 (2015).

32. Federmeier, K. D. & Kutas, M. A Rose by Any Other Name: Long-Term Memory Structure and Sentence Processing. J. Mem. Lang. 41, 469–495 (1999).

33. Hagoort, P., Baggio, G. & Willems, R. M. in The Cognitive Neurosciences (ed. Gazzaniga, M. S.) 819–836 (MIT Press, 2009).

34. Barber, H., Vergara, M. & Carreiras, M. Syllable-frequency effects in visual word recognition: evidence from ERPs. Neuroreport 15, 545–548 (2004).

35. Koivisto, M. & Revonsuo, A. Cognitive representations underlying the N400 priming effect. Cogn. Brain Res. 12, 487–490 (2001).

36. Rugg, M. D. The effects of semantic priming and word repetition on event-related potentials. Psychophysiology 22, 642–647 (1985).

37. Kuperberg, G. R., Sitnikova, T., Caplan, D. & Holcomb, P. J. Electrophysiological distinctions in processing conceptual relationships within simple sentences. Cogn. Brain Res. 17, 117–129 (2003).

38. Kim, A. & Osterhout, L. The independence of combinatory semantic processing: Evidence from event-related potentials. J. Mem. Lang. 52, 205–225 (2005).

39. Brouwer, H., Crocker, M. W., Venhuizen, N. j & Hoeks, J. C. J. A Neurocomputational Model of the N400 and the P600 in Language Comprehension. Cogn. Sci.

40. Herten, M. Van, Kolk, H. H. J. & Chwilla, D. J. An ERP study of P600 effects elicited by semantic anomalies. 22, 241–255 (2005).

41. Mcdonald, J. L. The Development of Sentence Comprehension Strategies in English and Dutch. 335, 317–335 (1986).

42. Hagoort, P. & Brown, C. M. ERP effects of listening to speech compared to reading: the P600 / SPS to syntactic violations in spoken sentences and rapid serial visual presentation. Neuropsychologia 38, 1531–1549 (2000).

43. Federmeier, K. D., Wlotko, E. W., De Ochoa-Dewald, E. & Kutas, M. Multiple effects of sentential constraint on word processing. Brain Res. 1146, 75–84 (2007).

44. Friedrich, M. & Friederici, A. D. N400-like semantic incongruity effect in 19-month-olds: Processing known words in picture contexts. J. Cogn. Neurosci. 16, 1465–77 (2004).

45. Atchley, R. A. et al. A comparison of semantic and syntactic event related potentials generated by children and adults. Brain Lang. 99, 236–246 (2006).

46. Kutas, M. & Iragui, V. The N400 in a semantic categorization task across 6 decades. Electroencephalogr. Clin. Neurophysiol. - Evoked Potentials 108, 456–471 (1998).

47. Gotts, S. J. Incremental learning of perceptual and conceptual representations and the puzzle of neural repetition suppression. Psychon. Bull. Rev. (2015). doi:10.3758/s13423-015-0855-y

48. McLaughlin, J., Osterhout, L. & Kim, A. Neural correlates of second-language word learning: minimal instruction produces rapid change. Nat. Neurosci. 7, 703–704 (2004).

49. Schultz, W., Dayan, P. & Montague, P. R. A neural substrate of prediction and reward. Science (80-.). 275, 1593–1599 (1997).

50. Friston, K. A theory of cortical responses. Philos. Trans. R. Soc. Lond. B. Biol. Sci. 360, 815–36 (2005).

51. McClelland, J. L. The interaction of nature and nurture in development: A parallel distributed processing perspective. Int. Perspect. Psychol. Sci. Vol. 1 Lead. Themes (1994).

52. Besson, M., Kutas, M. & Petten, C. Van. An Event-Related Potential (ERP) Analysis of Semantic Congruity and Repetition Effects in Sentences. J. Cogn. Neurosci. 4, 132–149 (1992).

53. Schott, B., Richardson-Klavehn, A., Heinze, H.-J. & Düzel, E. Perceptual priming versus explicit memory: dissociable neural correlates at encoding. J. Cogn. Neurosci. 14, 578–592 (2002).

54. Rumelhart, D. E. in Metaphor and Thought (ed. Ortony, A.) 71–82 (Cambridge University Press, 1979).

55. McCarthy, G., Nobre, A. C., Bentin, S. & Spencer, D. D. Language-Related Field Potentials in the Anterior-Medial Temporal Lobe: I. Intracranial Distribution and Neural Generators. J. Neurosci. 15, 1080–1089 (1995).

56. Nobre, A. C. & McCarthy, G. Language-Related Field Potentials in the Anterior-Medial Temporal Lobe: II. Effects of Word Type and Semantic Priming. J. Neurosci. 15, 1090–1098 (1995).

57. Sanford, A. J. & Sturt, P. Depth of processing in language comprehension: Not noticing the evidence. Trends Cogn. Sci. 6, 382–386 (2002).

58. Ferreira, F., Bailey, K. G. D. & Ferraro, V. Good-Enough Representations in Language Comprehension. Curr. Dir. Psychol. Sci. 11, 11–15 (2002).

59. Dronkers, N. F., Wilkins, D. P., Valin, R. D. Van, Redfern, B. B. & Jaeger, J. J. Lesion analysis of the brain areas involved in language comprehension. 92, 145–177 (2004).

60. Turken, A. U. & Dronkers, N. F. The neural architecture of the language comprehension network: converging evidence from lesion and connectivity analyses. Front. Syst. Neurosci. 5, 1–20 (2011).

61. Bookheimer, S. Functional MRI of language: New approaches to understanding the cortical organization of semantic processing. Annu. Rev. Neurosci. 25, 151–188 (2002).

62. Friederici, A. D. Towards a neural basis of auditory sentence processing. Trends Cogn. Sci. 6, 78–84 (2002).

63. Thompson-Schill, S. L., D’Esposito, M., Aguirre, G. K. & Farah, M. J. Role of left inferior prefrontal cortex in retrieval of semantic knowledge: A reevaluation. Proc. Natl. Acad. Sci. 94, 14792–14797 (1997).

64. Clayards, M., Tanenhaus, M. K., Aslin, R. N. & Jacobs, R. A. Perception of speech reflects optimal use of probabilistic speech cues. Cognition 108, 804–809 (2008).

65. Van Petten, C., Coulson, S., Rubin, S., Plante, E. & Parks, M. Time course of word identification and semantic integration in spoken language. J. Exp. Psychol. Learn. Mem. Cogn. 25, 394–417 (1999).

66. van den Brink, D., Brown, C. M. & Hagoort, P. The Cascaded Nature of Lexical Selection and Integration in Auditory Sentence Processing. J. Exp. Psychol. Learn. Mem. Cogn. 32, 364–372 (2006).

67. Rao, R. P. N. & Ballard, D. H. Predictive coding in the visual cortex: a functional interpretation of some extra-classical receptive-field effects. Nature 2, 79–87 (1999).

68. Rabovsky, M. & McRae, K. Simulating the N400 ERP component as semantic network error: Insights from a feature-based connectionist attractor model of word meaning. Cognition 132, 68–89 (2014).

69. Swinney, D. A. Lexical Access during Sentence Comprehension: (Re)Consideration of Context Effects. J. Verbal Learn. Behav. 18, 645–659 (1979).

70. Hoeks, J. C. J., Stowe, L. A. & Doedens, G. Seeing words in context: the interaction of lexical and sentence level information during reading. Cogn. Brain Res. 19, 59–73 (2004).

71. Osterhout, L. & Holcomb, P. J. Event-Related Brain Potentials Elicited by Syntactic Anomaly. J. Mem. Lang. 31, 785–806 (1992).

72. Brouwer, H. & Hoeks, J. C. J. A time and place for language comprehension: mapping the N400 and the P600 to a minimal cortical network. Front. Hum. Neurosci. 7, 758 (2013).

73. Coulson, S., King, J. W. & Kutas, M. Expect the Unexpected: Event-related Brain Response to Morphosyntactic Violations. Lang. Cogn. Process. 13, 21–58 (1998).

74. Sassenhagen, J., Schlesewsky, M. & Bornkessel-Schlesewsky, I. The P600-as-P3 hypothesis revisited: Single-trial analyses reveal that the late EEG positivity following linguistically deviant material is reaction time aligned. Brain Lang. 137, 29–39 (2014).

75. Polich, J. Updating P300 □: An integrative theory of P3a and P3b. Clin. Neurophysiol. 118, 2128–2148 (2007).

76. Schacht, A., Sommer, W., Shmuilovich, O., Casado Martinez, P. & Martin-Loeches, M. Differential Task Effects on N400 and P600 Elicited by Semantic and Syntactic Violations. PLoS One 9, 1–7 (2014).

77. Luck, S. J., Vogel, E. K. & Shapiro, K. L. Word meanings can be accessed but not reported during the attentional blink. Nature 383, 616–618 (1996).

78. Gleitman, L. R. & Gleitman, H. Phrase and paraphrase. (Norton, 1970).

79. Just, M. A. & Carpenter, P. A. A capacity theory of comprehension: Individual differences in working memory. Psychol. Rev. 99, 122–149 (1992).

80. Fischler, I., Bloom, P. A., Childers, D. G., Roucos, S. E. & Perry, N. W. Brain potentials related to stages of sentence verification. Psychophysiology 20, 400–409 (1983).

81. Staab, J. et al. Negation Processing in Context Is Not (Always) Delayed. 20, (2008).

82. Nieuwland, M. S. & Kuperberg, G. R. When the Truth Is Not Too Hard to Handle. Psychol. Sci. 19, 1213–1218 (2008).

83. van Berkum, J. J., Hagoort, P. & Brown, C. M. Semantic integration in sentences and discourse: evidence from the N400. J. Cogn. Neurosci. 11, 657–671 (1999).

84. Nieuwland, M. S. & Van Berkum, J. J. a. When peanuts fall in love: N400 evidence for the power of discourse. J. Cogn. Neurosci. 18, 1098–1111 (2006).

85. St. John F. M. The Story Gestalt: A Model of Knowledge-Intensive Processes’ in Text Comprehension. Cogn. Sci. 16, 271–306 (1992).

86. Hermann, K. M. et al. Teaching Machines to Read and Comprehend. in Proceedings of the 28th International Conference on Neural Information Processing Systems 1693–1701 (2015).

87. McCandliss, B. D., Cohen, L. & Dehaene, S. The visual word form area: Expertise for reading in the fusiform gyrus. Trends Cogn. Sci. 7, 293–299 (2003).

88. Bryant, B. D. & Miikkulainen, R. From Word Stream to Gestalt: A Direct Semantic Parse for Complex Sentences. (2001).

89. Hinton, G. I. Connectionist Learning Procedures. Mach. Learn. – an Artif. Intell. Approach III, 555–610 (1990).

90. Hinton, G. E., McClelland, J. L. & Rumelhart, D. E. in Parallel Distributed Processing (eds. Rumelhart, D. E. & McClelland, J. L.) 77–109 (MIT Press, 1986). doi:10.1146/annurev-psych-120710-100344

91. Sutton, R. S. & Barto, A. G. Reinforcement Learning: An Introduction. (MIT Press, 1998).

92. Rumelhart, D. E. & Todd, P. M. Learning and connectionist representations. Atten. Perform. XIV Synerg. Exp. Psychol. Artif. Intell. Cogn. Neurosci. 3–30 (1993).

93. McClelland, J. L. & Rogers, T. T. The parallel distributed processing approach to semantic cognition. Nat. Rev. Neurosci. 4, 310–322 (2003).

94. Pennington, J., Socher, R. & Manning, C. Glove: Global vectors for word representation. Emnlp2014.Org at <http://emnlp2014.org/papers/pdf/EMNLP2014162.pdf>

95. Gleitman, L. R., January, D., Nappa, R. & Trueswell, J. C. On the give and take between event apprehension and utterance formulation. Mem. Lang. 57, 544–569 (2007).

96. Altmann, G. T. M. & Kamide, Y. Incremental interpretation at verbs: restricting the domain of subsequent reference. Cognition 73, 247–264 (1999).

97. Kamide, Y., Altmann, G. T. M. & Haywood, S. L. The time-course of prediction in incremental sentence processing: Evidence from anticipatory eye movements. Mem. Lang. 49, 133–156 (2003).

98. Levy, R., Bicknell, K., Slattery, T. & Rayner, K. Eye movement evidence that readers maintain and act on uncertainty about past linguistic input. Proc. Natl. Acad. Sci. 106, 21086–21090 (2009).

99. Elman, J. L. Finding Structure in Time. Cogn. Sci. 14, 179–211 (1990).

100. Mikolov, T., Deoras, A., Povey, D., Burget, L. & Cernocky, J. H. Strategies for Training Large Scale Neural Network Language Models. in IEEE Workshop on Automatic Speech Recognition and Understanding (2011).

101. Delong, K. A., Quante, L. & Kutas, M. Predictability, plausibility, and two late ERP positivities during written sentence comprehension. Neuropsychologia 61, 150–162 (2014).

